# Functional Organization of Glycolytic Metabolon on Mitochondria

**DOI:** 10.1101/2023.08.26.554955

**Authors:** Haoming Wang, John Vant, Youjun Wu, Richard Sánchez, Mary Lauren Micou, Andrew Zhang, Vincent Luczak, Seungyoon Blenda Yu, Mirna Jabbo, Seokjun Yoon, Ahmed Abdallah Abushawish, Majid Ghassemian, Eric Griffis, Marc Hammarlund, Abhishek Singharoy, Gulcin Pekkurnaz

## Abstract

Glucose, the primary cellular energy source, is metabolized through glycolysis initiated by the rate-limiting enzyme Hexokinase (HK). In energy-demanding tissues like the brain, HK1 is the dominant isoform, primarily localized on mitochondria, crucial for efficient glycolysis-oxidative phosphorylation coupling and optimal energy generation. This study unveils a unique mechanism regulating HK1 activity, glycolysis, and the dynamics of mitochondrial coupling, mediated by the metabolic sensor enzyme O-GlcNAc transferase (OGT). OGT catalyzes reversible O-GlcNAcylation, a post-translational modification, influenced by glucose flux. Elevated OGT activity induces dynamic O-GlcNAcylation of HK1’s regulatory domain, subsequently promoting the assembly of the glycolytic metabolon on the outer mitochondrial membrane. This modification enhances HK1’s mitochondrial association, orchestrating glycolytic and mitochondrial ATP production. Mutations in HK1’s O-GlcNAcylation site reduce ATP generation, affecting synaptic functions in neurons. The study uncovers a novel pathway that bridges neuronal metabolism and mitochondrial function via OGT and the formation of the glycolytic metabolon, offering new prospects for tackling metabolic and neurological disorders.

## INTRODUCTION

In cellular environments, metabolic reactions are more complex than a simple cascade of enzymes and metabolites operating within diffusion limits. Rather, an advanced level of organization leverages the compartmentalization of metabolic enzymes to organelles such as mitochondria and other subcellular structures to optimize biochemical transformations. This complex spatial organization, often referred to as a “metabolon,” delineates the physical boundaries of metabolic fluxes within the cytoplasm^1-3^. Metabolons enhance metabolic efficiency by co-compartmentalizing enzymes with their sequential substrates, thereby minimizing diffusional transit times, promoting metabolite channeling, providing an optimized microenvironment for individual pathways, and limiting counterproductive metabolite-enzyme interactions. Despite the pivotal role of this spatial organization in ensuring efficient substrate channeling and metabolic flux regulation, our mechanistic insight into the regulation of metabolon formation, particularly within key metabolic pathways like glycolysis, remains sparse^4,5^.

Glycolysis mediates the conversion of glucose to pyruvate, which is then further metabolized to generate ATP. A key enzyme in this process is Hexokinase (HK), which catalyzes the initial, rate-limiting step of phosphorylating glucose into glucose-6-phosphate. Importantly, this step consumes ATP. Hexokinase 1 (HK1), one of the distinct isoforms of HK expressed in metabolically demanding cells such as neurons, is predominantly localized on the outer mitochondrial membrane^6-8^. This subcellular proximity to mitochondria implies a role for HK1 in modulating metabolic efficiency^9^. Although previous studies have explored the role of HK1 in glycolysis, the specific molecular pathway that dictate HK1 activity, its mitochondrial localization, and its potential involvement in glycolytic metabolon assembly remain elusive.

This study investigates a novel molecular mechanism that modulates the activity and mitochondrial localization of HK1 through the action of the metabolic sensor enzyme O-GlcNAc transferase (OGT). OGT catalyzes a unique post-translational modification, O-GlcNAcylation, by attaching a GlcNAc sugar moiety to serine and threonine residues on proteins. Intracellular UDP-GlcNAc concentrations, modulated by glucose flux through the hexosamine biosynthetic pathway, regulate this process^10^. Our findings reveal that HK1 undergoes dynamic O-GlcNAcylation at its regulatory domain when OGT activity is elevated. This modification enhances HK1’s localization to the mitochondria, leading to increased rates of both glycolytic and mitochondrial ATP production. This is achieved by facilitating the formation of a glycolytic metabolon on the mitochondrial outer membrane. In contrast, mutations at the O-GlcNAcylation site of HK1 result in reduced ATP production rates and dysregulation of the energy-demanding process of presynaptic vesicle recycling in neurons. In conclusion, our study uncovers novel molecular pathways linking neuronal metabolism to mitochondrial function via OGT-mediated modulation of HK1. We identify HK1 O-GlcNAcylation as a new mechanism regulating glycolysis and metabolon formation that is critical for metabolic efficiency, offering potential implications for the development of targeted interventions for metabolic and neurological disorders.

## RESULTS

### Glucose modulates mitochondrial localization of Hexokinase 1

To characterize the Hexokinase 1 (HK1) localization at the subcellular level, we carried out immunofluorescence analysis in disassociated rat hippocampal neurons. Using hippocampal neurons transfected with Mito-DsRed as a mitochondrial marker, we observed over 80% of endogenous HK1 resided alongside mitochondria within the soma, dendrites, and axon (Figure 1A) as previously reported^11^.

**Figure 1.**
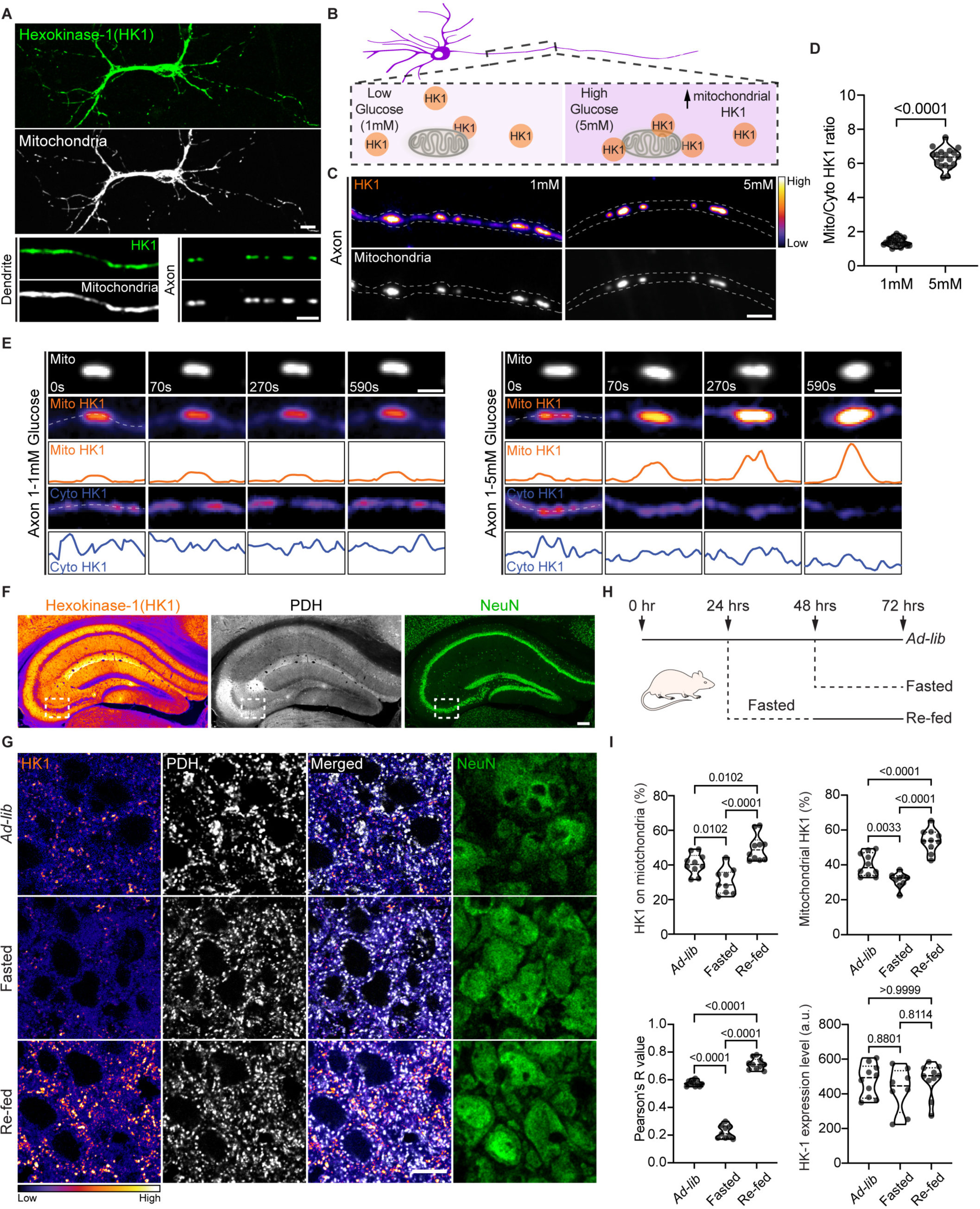
Glucose-dependent regulation of Hexokinase 1 localization. (A) Immunofluorescence staining of disassociated rat hippocampal neurons for endogenous Hexokinase 1 (HK1; green) and mitochondrial marker (MitoDsRed; gray), illustrating subcellular localization of HK1 in the somatodendritic region, as well as along axonal and dendritic lengths. (B-E) Hippocampal neurons cultured in 5mM glucose, co-transfected with MitoDsRed (gray), wild-type human (WT) HK1 tagged with eGFP, and rat shRNA-HK1 to achieve endogenous level of HK1 (pseudocolor, fire), then transferred to 1mM glucose for 72 hours. (C) Representative axonal images were captured following a 2-hour exposure to 5mM glucose or at 1mM glucose, as summarized in (B) (also see Figure S1A for schematic of extracellular glucose manipulation paradigm). Scale bar represents 5µm, pseudocolor scale indicates low to high intensity. (D) Quantification of mitochondrial (Mito) and cytoplasmic (Cyto) HK1 intensity ratios along axons. Data are presented as a violin plot with individual data points and associated p-value. n = 94-103 mitochondria, 9-10 axons from three independent experiments (Mann-Whitney *U* test) (E) Spatiotemporal changes in immunofluorescence intensity of HK1 was measured by live-cell imaging for 10 minutes along the length of an axon within cytoplasmic (Cyto; blue intensity plot) and mitochondrial (Mito; orange intensity plot) compartments during a medium switch from 1 to 1mM glucose or 1 to 5mM glucose (Scale bar represents 2µm). (F-I) Immunofluorescence staining of adult mouse brain coronal slices (Bregma, -2mm) with antibodies against HK1 (pseudocolor, fire), mitochondrial marker Pyruvate Dehydrogenase (PDH; gray) and neuronal marker NeuN (green). (F) The hippocampal section at ad libitum fed state (Ad-lib). Scale bar represents 100µm. (G) Enlarged images of white dashed boxes in (F) display the HK1 distribution pattern in the CA3 region of the hippocampus in Ad-lib, after 24 hours fasting (Fasted) and 24 hours of re-feeding (Re-fed) states. Scale bar represents 10µm. (H) Schematic illustration of the experimental regimen for altering blood glucose levels through fasting and refeeding in mice, used to compare HK1 localization in the brain. (I) Co-localization analysis to measure the percent intensity of HK1 on mitochondria and mitochondrial HK1, as well as the Pearson’s correlation coefficient (R value) and quantification of HK1 expression level for each condition. Data are presented as violin plots with individual data points and associated p-values. n = 27-30 hippocampal CA3 regions, 9 mice from three independent experiments (one-way ANOVA with post hoc Kruskal-Wallis multiple comparison test). See also Figure S1.

Given that glucose serves as the substrate for HK1, and mitochondria are ATP-rich, we hypothesized that glucose flux could be a key regulator of HK1’s mitochondrial positioning (Figure 1B). To test the effect of glucose flux on HK1’s positioning, we cultured primary neurons in media containing 5 mM glucose. These neurons were then subjected to a 72-hour glucose reduction to 1 mM, followed by a subsequent increase back to 5 mM, before imaging (Figure S1A)^12^. We also established a system for live imaging of HK1 and mitochondria using neurons co-expressing Mito-DsRed and shRNA resistant HK1-GFP, together with a short hairpin RNA (shRNA) designed to deplete endogenous HK1. The efficacy of this shRNA had already been validated in Neuro-2A cells (Figures S1B and S1C). By using this approach, we ensured that the HK1 level remained consistent with endogenous levels (Figures S1C, and as detailed in Figure S4F and S4G). Our results revealed a direct response to glucose flux: an increase in extracellular glucose prompted mitochondrial enrichment of HK1 in neurons (Figure 1B-E). To further examine the role of glucose flux, we subjected the neurons to a 2-hour glucose deprivation period, followed by an immediate reintroduction of glucose to 5 mM before imaging (Figure S1D-F). Pyruvate (1 mM) was present throughout these experiments as an alternate fuel source. Remarkably, a transient decrease in extracellular glucose resulted in the downregulation of mitochondrial HK1. Upon reintroduction of glucose to 5 mM, however, we observed a rebound recruitment of HK1 to the mitochondria within the same neuron (Figures S1D-F). The size of the mitochondria remained unaffected by these fluctuations in glucose (Figure S1G).

### Feeding state influences Hexokinase 1 positioning *in vivo* in mouse brain

Expanding our investigation *in vivo*, we performed feeding and fasting experiments with mice (Figure 1F-H). We confirmed that fluctuations in blood glucose levels mirrored the feeding cycles, as previously reported^12^. Mice subjected to a 24-hour fasting exhibited a drop in blood glucose, which normalized upon a subsequent 24-hour feeding period (Figure S1H). Upon examining HK1 localization within the CA3 region of the hippocampus in ad-libitum fed (*Ad-lib*), fasted, and re-fed mice, we found a clear correlation with blood glucose levels (Figure 1F-I). Fasting, and the consequent drop in blood glucose, resulted in a downregulation of mitochondrial HK1. In contrast, refeeding, and the subsequent rise in blood glucose, led to an upregulation of mitochondrial HK1 compared to *ad-lib* conditions (Figures 1I). Importantly, neither feeding nor fasting altered overall HK1 expression levels (Figure 1I).

### O-GlcNAc transferase activity promotes Hexokinase 1 recruitment to mitochondria

The hexosamine biosynthetic pathway integrates glucose flux signals, culminating in the production of UDP-GlcNAc, an essential substrate for O-GlcNAcylation. Acting as a metabolic sensor that reflects the cell’s nutritional status, this pathway dynamically regulates O-GlcNAc addition by OGT. This addition is counterbalanced by its removal through the enzyme O-GlcNAcase (OGA) (Figure 2A). Given the pathway’s previously recognized role in mitochondrial anchoring, we hypothesized that OGT and OGA might collaboratively dictate the positioning of HK1 on the mitochondria in response to glucose flux changes to enhance metabolic efficiency^12,13^. To determine whether OGT and OGA activity influences HK1 localization in hippocampal neurons, we pharmacologically and genetically manipulated this enzyme pair and modulated O-GlcNAcylation levels (Figure S2A-C). Both biochemical analysis of mitochondrial fractions (Figure 2B and 2C) and imaging data suggested (Figure 2D and 2E) that enhancing O-GlcNAcylation increased mitochondrial HK1, while reducing O-GlcNAcylation decreased mitochondrial HK1 in neurons. Importantly, mitochondrial size remained unaffected by O-GlcNAcylation level fluctuations (Figure S2B). This phenotype was also observed in HEK293T cells, a cell type expressing HK1 as predominant hexokinase isoform, further confirming our findings (Figure S2D and S2E).

**Figure 2.**
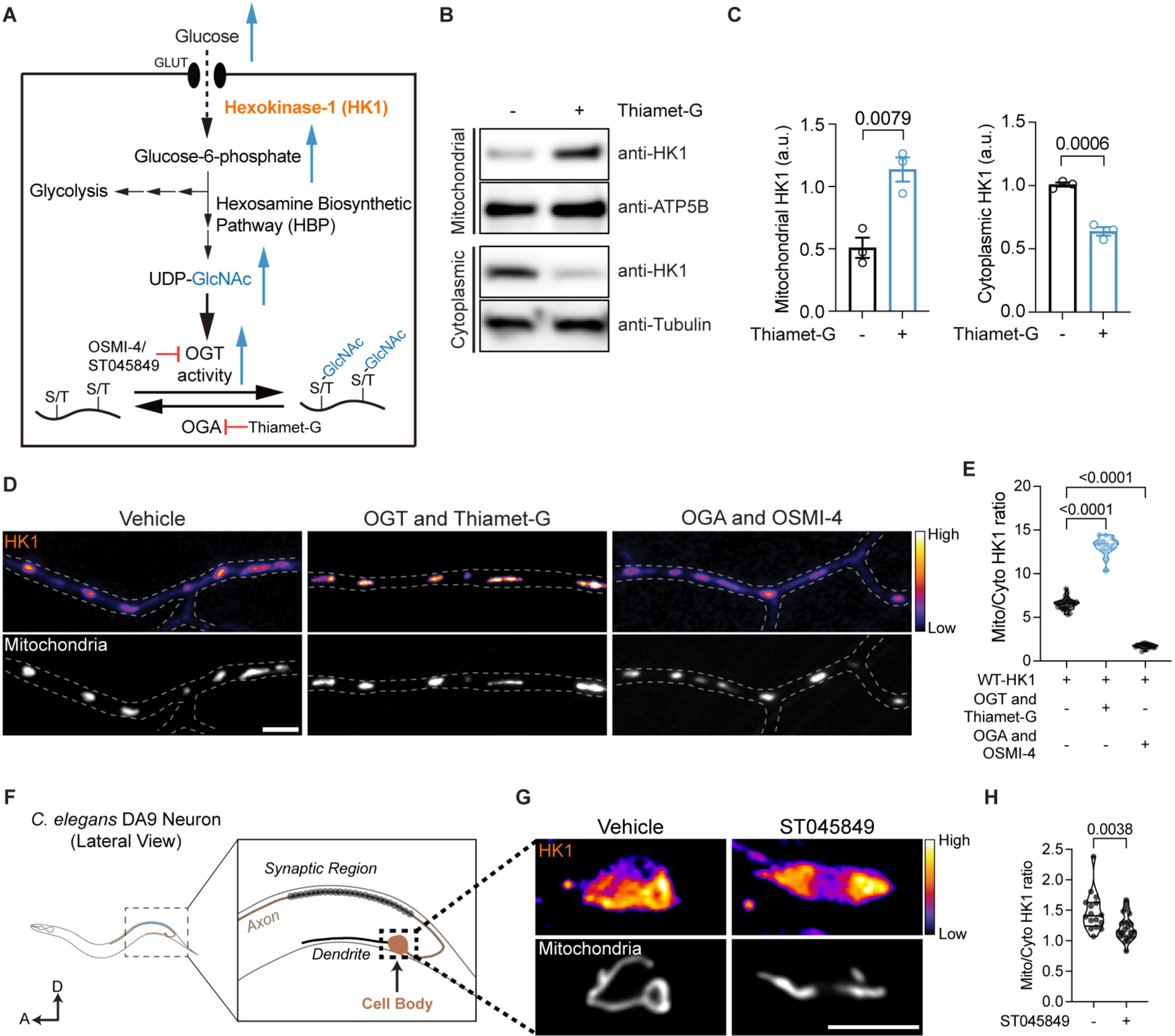
O-GlcNAcylation regulates mitochondrial localization of Hexokinase 1. (A) Schematic representation of glucose metabolism via Hexosamine Biosynthetic Pathway (HBP). Rate-limiting steps and inhibitors used in this study are also indicated. Increased glucose flux upregulates HBP pathway, UDP-GlcNAc synthesis, and OGT activity. (B and C) Mitochondrial and cytoplasmic fractions, obtained from rat cortical neurons treated overnight with OGA inhibitor Thiamet-G to enrich neuronal O-GlcNAcylation or vehicle (DMSO), were separated by SDS gel electrophoresis and probed with anti-HK1, anti-ATP5β (mitochondrial marker), and anti-tubulin (cytoplasmic marker). (C) Quantification of HK1 bands normalized to the intensity of ATP5β (mitochondrial fraction) or tubulin (cytoplasmic fraction) for each condition. n = 3 independent experiments (All values are shown as mean ± SEM; unpaired *t*-test). (D and E) Manipulating O-GlcNAcylation affects HK1 localization in neurons. (D) Axonal segments of hippocampal neurons cultured in 5mM glucose, co-transfected with MitoDsRed (gray), rat shRNA-HK1, and wild-type human (WT) HK1 tagged with eGFP(pseudocolor, fire) to achieve endogenous expression level of HK1. O-GlcNAcylation level was upregulated by ectopic expression of OGT and Thiamet-G treatment, and downregulated by expression of OGA and OSMI-4 treatment. Scale bar represents 5µm. (E) Quantification of mitochondrial (Mito) and cytoplasmic (Cyto) HK1 intensity ratios along axons. Data are presented as a violin plot with individual data points and associated p-values. n = 83-120 mitochondria from 11-13 axons, three independent experiments (one-way ANOVA with post hoc Kruskal-Wallis multiple comparison test). (F-H) In vivo imaging of endogenously-GFP tagged HK1 in C. elegans neurons. (F) Schematic representation of the DA9 neuron cell body in C. elegans nervous system. (G) DA9 neurons expressing (hxk-1) HK1-7xSplitGFP (pseudo-color, fire) and mito-TagRFP (gray) following vehicle (DMSO) or OGT inhibitor ST045849 treatments. Scale bar represents 5μm. (H) Mitochondrial and cytoplasmic HK1-7xSplitGFP ratios were quantified from DA9 neuron cell bodies for each condition. Data are presented as a violin plot with individual data points and associated p-value. n = 16-23 cell bodies, three independent experiments (unpaired *t*-test). See also Figure S2.

### O-GlcNAcylation modulates Hexokinase 1 positioning *in vivo* in *C. elegans*

Building on our findings that O-GlcNAcylation plays a key role in controlling HK1’s mitochondrial localization in mammalian cells, we expanded our inquiry to *C. elegans*. Given the conservation of OGT and HK1 in *C. elegans*, we investigated whether O-GlcNAcylation could regulate HK1 positioning *in vivo*. We engineered *C. elegans* to express HK1 endogenously tagged with 7xSplitGFP in the DA9 neuron (Figure S2F). To attenuate O-GlcNAcylation without overexpressing OGA, we treated the nematodes with an OGT inhibitor (ST045849), that was previously validated specifically in *C. elegans*^14^. Upon imaging the cell body of the DA9 neuron (Figure 2F), we observed that a reduction in O-GlcNAcylation corresponded to a decrease in mitochondrial HK1 (Figures 2G and 2H). These *in vivo* observations parallel the outcomes seen with OSMI-4 treatments in cultured rodent hippocampal neurons, highlighting the conserved role of O-GlcNAcylation in regulating HK1 positioning across different genera.

### Hexokinase 1 undergoes regulated O-GlcNAcylation

To elucidate whether HK1 is regulated via O-GlcNAcylation directly, we employed a CRISPR-based strategy to add a GFP tag to the C-terminus of endogenous HK1 in HEK293T cells (Figure S3A), a tagging strategy known not to hinder HK1’s function^11^. We then immunoprecipitated (IP) HK1 from these genetically engineered cells and resolved proteins on an SDS-PAGE gel (Figure 3A and 3B, S3B). To determine whether the extent of HK1 O-GlcNAcylation was susceptible to changes in OGT and OGA activity, we upregulated OGT activity by ectopic expression and applied the OGA inhibitor Thiamet-G. Anti-GlcNAc antibodies detected a band co-migrating with HK1 in anti-GFP IP from the CRISPR-modified HEK293T cells (Figure 3A). This band’s intensity was amplified when OGT was overexpressed and OGA was concurrently inhibited (Figures 3A and 3B). Further, we performed IP on O-GlcNAcylated mitochondrial proteins from rat cortical neurons and compared the samples with and without Thiamet-G-induced OGA inhibition (Figure S3C and S3D). A significant amplification of O-GlcNAcylated mitochondrial HK1 was observed upon OGA inhibition (Figure S3D) in neurons, suggesting that under basal conditions, HK1’s O-GlcNAcylation sites are not saturated, and the modification level is likely determined by the dynamic equilibrium between OGT and OGA activity. We extended our investigation to Hexokinase 2 (HK2), examining its O-GlcNAcylation cycling status. Despite the detection of an O-GlcNAcylated HK2 band in anti-GFP immunoprecipitants from HEK293T cells (Figure S3E and S3F), this band’s intensity did not increase upon Thiamet-G-mediated OGA inhibition or OGT overexpression (Figures S3E and S3F). These results imply that while HK2 is subject to baseline O-GlcNAcylation, the extent of this modification is unaffected by alterations in OGT or OGA activity.

**Figure 3.**
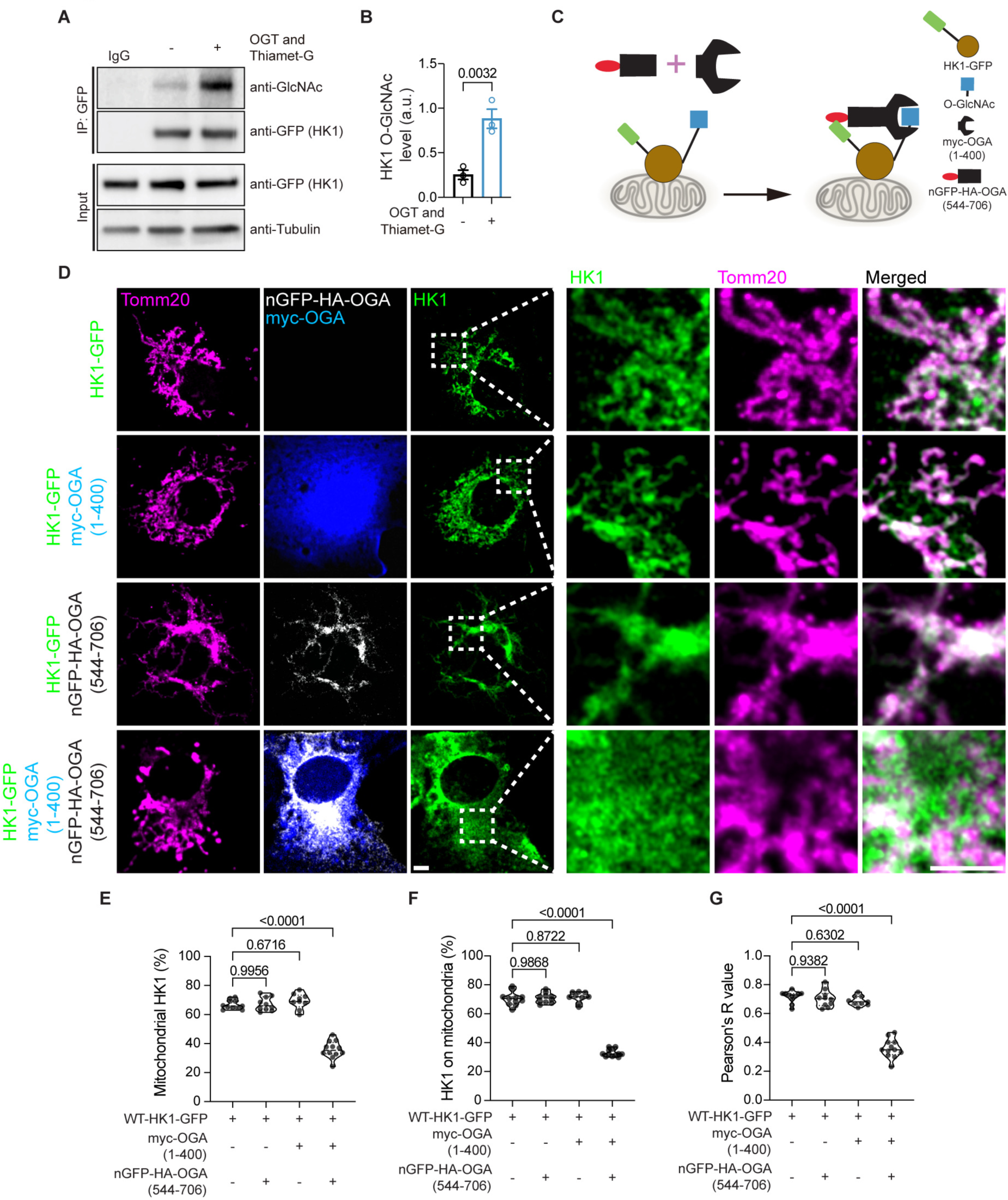
OGT and OGA regulate Hexokinase 1 O-GlcNAcylation. (A and B) The impact of O-GlcNAcylation upregulation, elicited by ectopic OGT expression and Thiamet-G treatment, on HK1 was investigated using immunoprecipitation (IP) of endogenous HK1 from HEK293T-EGFP-HK1 cells (CRISPR-edited cells expressing endogenously GFP-tagged HK1). HK1 was immunoprecipitated using an anti-GFP antibody from cells treated overnight with either vehicle (DMSO) or Thiamet-G, with or without OGT expression. Rabbit IgG served as a negative IP control. Input lanes, which were loaded with 3% of cell lysates used for IP, were also probed with an anti-tubulin antibody to confirm equal loading. (B) HK1 O-GlcNAcylation was quantified by normalizing the intensity of each GlcNAc band to the corresponding HK1 (GFP) band. OGT overexpression resulted in a significant increase in HK1 O-GlcNAcylation (B). n = 3 independent experiments (mean ± SEM; unpaired *t*-test). (C-G) Selective downregulation of HK1 O-GlcNAcylation by a nanobody-fused split OGA eraser and evaluation of subcellular localization. (C) Schematic illustrating the approach to selectively downregulate HK1 O-GlcNAcylation using eGFP-tagged HK1 (HK1-GFP) and GFP-nanobody (nGFP)-directed split O-GlcNAc eraser (nGFP-HA-OGA (544-706) and myc-OGA (1-400)). The OGA enzyme was engineered into a split and truncated form with limited substrate activity. nGFP-HA-OGA (544-706) recognizes HK1-GFP and redirects the catalytic half of OGA myc-OGA (1-400), to remove the O-GlcNAc modification from HK1. (D) Representative images of COS-7 cells expressing HK1-GFP (green), myc-OGA (1-400) (blue) and nGFP-HA-OGA (544-706) (gray). The subcellular localization of HK1 was examined using immunofluorescence staining with anti-Tomm20 (mitochondrial marker; magenta), anti-myc (myc-OGA (1-400); blue) and anti-HA (nGFP-HA-OGA (544-706); gray) antibodies. The HK1 distribution pattern is highlighted in the enlarged images corresponding to the white dashed boxes in (D). Scale bars represents 5µm. (E-G) Quantitative analysis of HK1 co-localization to assess the percentage of mitochondrial HK1 intensity (E), HK1 intensity on mitochondria (F), and the Pearson’s correlation coefficient (R value) for each condition (G). Data are presented as violin plots with individual data points and associated p-values. n = 42 cells from three independent experiments (one-way ANOVA with post hoc Kruskal-Wallis multiple comparison test). See also Figure S3.

### Nanobody-mediated control of O-GlcNAcylation regulates Hexokinase 1 localization

To investigate the role of HK1 O-GlcNAcylation in determining its subcellular localization, we adopted an approach to selectively attenuate HK1’s O-GlcNAcylation. In COS-7 cells, we co-expressed GFP-tagged HK1 along with a GFP nanobody-fused, split O-GlcNAcase (nGFP-OGA)^15-17^(Figure 3C). The enzymatic activity of OGA was only present when both myc-OGA (1-400) and nGFP-HA-OGA (544-706) were concurrently expressed (Figure 3C and 3D). To ensure consistency, we confirmed that the expression levels of split OGA plasmids and HK1 were equal across conditions (Figure S3G-I). Following cell fixation, we performed staining with Tomm20, a protein that serves as a mitochondrial marker (Figures 3D). The targeted reduction of HK1’s O-GlcNAcylation led to a decrease in mitochondrial HK1 within COS-7 cells (Figures 3D-G). Our findings, therefore, highlight that the level of HK1 O-GlcNAcylation modulates its mitochondrial localization in both neurons and COS-7 cells.

### Identification and Confirmation of O-GlcNAcylation Sites in Hexokinase 1

To elucidate the presence and location of O-GlcNAcylation sites within Hexokinase 1 (HK1), we IP’ed this protein from HEK293T cells. The cells were subjected to both basal and enhanced O-GlcNAcylation conditions. The resulting HK1 protein was then resolved on an SDS-PAGE gel and probed with anti-GlcNAc antibodies. This process confirmed that HK1 O-GlcNAcylation levels indeed rise in response to an upregulated O-GlcNAcylation environment (Figure 4A and 4B). Proteins with a molecular weight approximating that of HK1 were subsequently extracted from the SDS-PAGE gel. Peptides were isolated from these proteins through a trypsin and gluC co-digestion and subsequently analyzed using tandem mass spectrometry. This in-depth analysis yielded the identification of a single O-GlcNAc modification site within HK1 (Figure S4A). Notably, the site of this modification, T259, is unique to HK1 among the three hexokinase isoforms. It is, however, conserved across most mammalian HK1 and *C. elegans* (Figure 4C). This finding underscores the potentially critical role of O-GlcNAcylation at T259 in the functionality of HK1.

**Figure 4.**
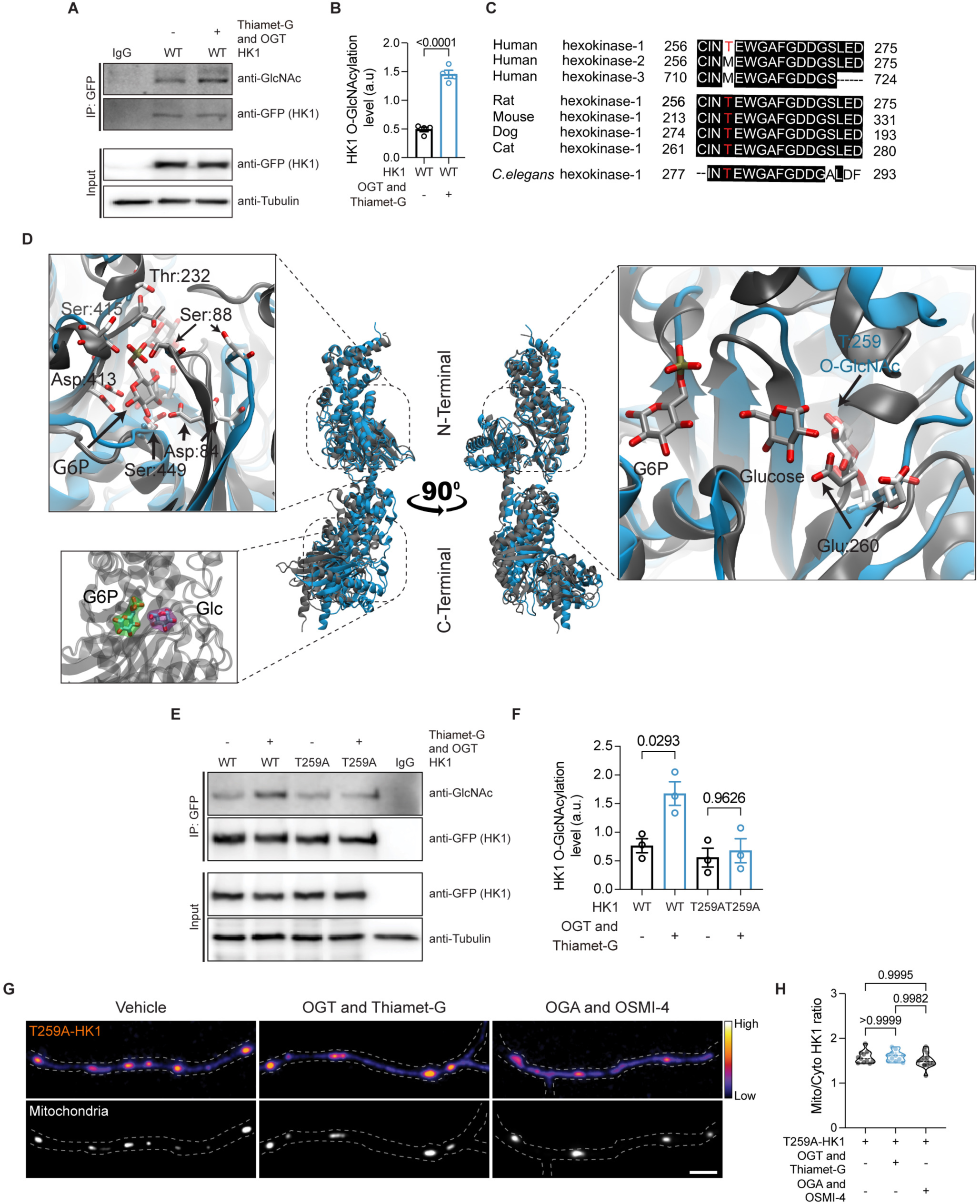
OGT-dependent regulation of Hexokinase 1 localization requires O-GlcNAcylation. (A-F) Identification and characterization of HK1 O-GlcNAcylation site. (A) To identify the O-GlcNAc modification site, eGFP tagged HK1 was overexpressed with or without OGT in HEK293T cells, and immunoprecipitated (IP) with a GFP antibody following overnight vehicle (DMSO) or Thiamet-G treatment. Input lanes contained 3% of the cell lysates used for the IP and were also probed with anti-tubulin antibody as a loading control. HK1 O-GlcNAcylation levels were quantified by normalizing the intensity of each GlcNAc band to the intensity of HK1 (GFP) bands. OGT overexpression and Thiamet-G treatment increased HK1 O-GlcNAcylation (B), this condition was selected for mass spectrometry sample preparation. n = 4 independent experiments (mean ± SEM and associated p-values; unpaired *t*-test). (C) Sequence alignment of human HK1 (amino acid residues 256-275) from different species showing conserved O-GlcNAcylated threonine (T) residue (red) (T259), and human hexokinase isoforms 2 and 3. Conserved amino acids are indicated as black box. (D) Structural representation of the conformational differences between the equilibrated structure of the full unmodified HK1 (gray) and O-GlcNAcylated HK1 (blue). The Cα atoms in residues 20 to 460 were used for alignment. Top left inset shows key residues (Ser88, Thr232, and Ser415) responsible for coordinating the phosphate of G6P and the 2’ hydroxyl (Asp84, Asp413, and Ser449). Bottom left inset shows the binding pockets for G6P (green VDW iso-surface) and glucose (purple VDW iso-surface). Right inset shows HK1 with O-GlcNAcylation at T259, the single-turn alpha-helix (residues 260 to 264) forms a flexible loop to accommodate the bulky O-GlcNAc with the positions of G6P and glucose from the crystal structure shown for reference. (E) eGFP-tagged HK1, either wild type (WT) or with the putative O-GlcNAc site mutated (T259A), was expressed in HEK293T cells with or without OGT. Following overnight vehicle (DMSO) or Thiamet-G treatment, HK1 immunoprecipitated (IP) using a GFP antibody. HK1 immunoprecipitants were analyzed with anti-GlcNAc and anti-GFP antibodies, and GlcNAcylation levels on HK1 were quantified. (F) The intensity of each GlcNAc band was normalized to the intensity of the GFP band for quantification of WT and T259A HK1 O-GlcNAcylation levels. n = 3 independent experiments (mean ± SEM and associated p-values; One-way ANOVA with post hoc Kruskal-Wallis test). (G) Hippocampal neurons cultured in 5mM glucose, co-transfected with MitoDsRed (gray), rat shRNA-HK1, and O-GlcNAc site mutated T259A HK1 tagged with eGFP to achieve endogenous expression level of HK1 (pseudocolor, fire). Neuronal O-GlcNAcylation levels were upregulated by ectopic expression of OGT and Thiamet-G treatment, and downregulated by expression of OGA and OSMI-4 treatment. (H) The mitochondrial (Mito) and cytoplasmic (Cyto) HK1 intensity ratios were quantified along axons. Data are presented as a violin plot with individual data points and associated p-values. n = 81-86 mitochondria from 10-13 axons, three independent experiments (one-way ANOVA with post hoc Kruskal-Wallis multiple comparison test, Scale bar represents 5µm). See also Figure S4.

Having established the specific O-GlcNAc modification site within HK1, we next sought to understand how this modification might influence the protein’s structural conformation. To determine the effect of T259 O-GlcNAcylation on the structure of HK1, we turned to computational modeling (Figure 4D). The HK1 structure is composed of N- and C-terminal halves: the N-terminal half is the regulatory domain, while the C-terminal half exhibits catalytic activity, participating in glucose binding and glucose phosphorylation via ATP. When glucose-6-phosphate (G6P) binds at the N-terminal of HK1, it impedes HK1’s interaction with mitochondria, creating a negative feedback loop. A linker region joining the two halves is primarily composed of a 28 residue-long (residues 448 to 476) alpha helix^18^. This linker region is implicated in regulating the coupling between the G6P-occupancy of the N-terminal with the glucose reactivity of the C-terminal half^18^ by perturbing the C-terminal binding pocket through a network of allosteric interactions and altering the conformation of the catalytic binding pocket to favor the binding of G6P over ATP. However, little has been reported on the relevance of N-terminal G6P binding to mitochondrial membrane-hexokinase interactions. Interestingly, the T259 O-GlcNAcylation residue is located near the non-catalytic N-terminal pocket. Through Resolution Exchange Solute Tempering (REST2) molecular dynamics simulations^19,20^, we revealed a conformational transition state of the O-GlcNAc modified HK1, which results in a loss of secondary structure and a looser binding pocket compared to the unmodified HK1 (Figure 4D). At the site of O-GlcNAcylation, the single-turn alpha-helix (residues 260 to 264) forms a flexible loop to accommodate the bulky O-GlcNAc. Additionally, key residues (Ser88, Thr232, and Ser415) responsible for coordinating the phosphate and the 2’ hydroxyl (Asp84, Asp413, and Ser449) of G6P are offered more significant conformational flexibility to accommodate the bulky O-GlcNAc moiety without disrupting packing of the pocket. This enhanced conformational freedom of the residues prevents a stable interaction with G6P. Thus, HK1 has a largescale conformational rearrangement upon O-GlcNAcylation (Figure 4D right) with an N-terminal Root Mean Squared Deviation (RMSD) of 3.5 Å (aligned and calculated with Cα atoms in residues 20 to 460 using the crystal structure as a reference).

### OGT-dependent mitochondrial localization of Hexokinase 1 requires T259 O-GlcNAcylation

In order to elucidate the significance of T259 O-GlcNAcylation in regulating the mitochondrial localization of HK1, we employed site-directed mutagenesis to substitute Threonine 259 with Alanine, thereby abolishing O-GlcNAcylation at this specific residue. Subsequently, we expressed and immunoprecipitated both the wild-type (WT) HK1 and the O-GlcNAc-deficient T259A-HK1 variant from HEK239T cells. The immunoprecipitants were subjected to SDS-PAGE analysis and probed with anti-GlcNAc antibodies to validate the successful disruption of O-GlcNAcylation in the T259A-HK1 mutant (Figure 4E and 4F). To investigate the impact of O-GlcNAcylation at T259 on HK1’s mitochondrial localization, we conducted imaging experiments using HEK293T cells expressing either WT-HK1 or the T259A-HK1 mutant, in conjunction with Mito-DsRed. To simulate conditions of heightened glucose flux, we employed OGT overexpression and O-GlcNAcase (OGA) inhibition using Thiamet-G. We observed a pronounced increase in the mitochondrial localization of WT-HK1 (Figure S4B and S4C), whereas no such effect was observed in cells expressing the T259A-HK1 mutant (Figure S4D and S4E).

Expanding our investigation to hippocampal neurons, we expressed T259A-HK1-GFP using our shRNA approach (Figure S4F and S4G). By modulating O-GlcNAcylation levels through ectopic OGT expression and OGA inhibition or ectopic OGA expression and OGT inhibition using OSMI-4, we aimed to replicate conditions of increased or decreased glucose flux, respectively, as outlined in Figure S2C. Analysis of the mitochondrial to cytoplasmic HK1 ratio along the axon of the neuron revealed that the absence of O-GlcNAcylation at the T259 site rendered HK1’s mitochondrial localization unresponsive to changes in O-GlcNAcylation levels (Figure 4G and 4H), without affecting mitochondrial size (Figure S4H).

### O-GlcNAcylation modulates Hexokinase 1 activity and ATP Production Rates

Mitochondrial binding allows HK1 to have prime access to mitochondrially generated ATP, thus promoting more effective glucose phosphorylation^21^. To investigate the role of O-GlcNAcylation in modulating HK1 activity, we measured the levels of G6P, which is the product of HK1-mediated glucose phosphorylation (Figure 5A). We expressed WT-HK1 or the T259A-HK1 in HEK239T cells and manipulated O-GlcNAcylation levels by the ectopic expression of OGT and inhibition of OGA using Thiamet-G. Our results revealed that increasing O-GlcNAcylation levels led to elevated G6P levels in cells expressing WT-HK1, while this effect was absent in cells expressing T259A-HK1 (Figure 5B). Importantly, the expression levels of both WT-HK1 and T259A-HK1 remained constant (Figure S5A and S5B), indicating that the observed differences in G6P levels were specifically attributed to O-GlcNAcylation of HK1. Upregulation of O-GlcNAcylation in HEK293T cells also enhanced production of G6P, without altering endogenous HK1 expression level (Figure S5C-E).

**Figure 5.**
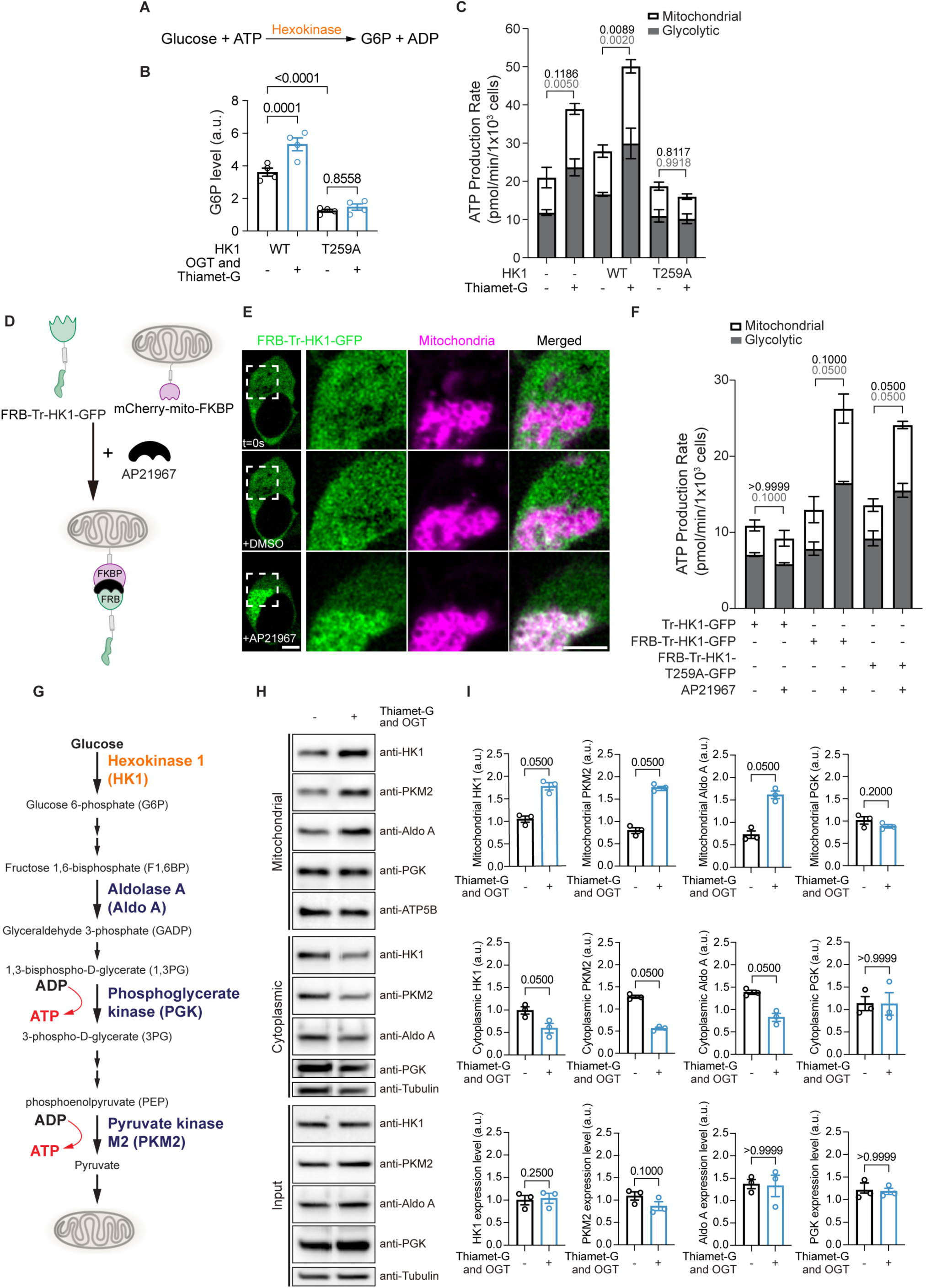
Hexokinase 1 O-GlcNAcylation enhances metabolic efficiency. (A) The hexokinase enzyme catalyzes the first step of glucose metabolism, where glucose and ATP are converted into glucose-6-phosphate (G6P) and ADP. (B) Quantification of G6P levels in HEK239T cells expressing eGFP-tagged WT or T259A HK1 with or without ectopic OGT expression and overnight Thiamet-G treatment. n = 4 independent experiments (mean ± SEM and associated p-values; One-way ANOVA with post hoc Kruskal-Wallis test). (C) Mitochondrial and glycolytic ATP production rates calculated from HEK293T cells expressing control vector, WT or T259A HK1, following overnight vehicle or Thiamet-G treatments. n= 15–17 wells, three independent experiments. (D) The schematic illustrates the strategy for inducing the mitochondrial relocation of truncated HK1 (Tr-HK1), which naturally resides in the cytoplasm due to the absence of a mitochondrial localization sequence. Relocation is accomplished with the use of the dimerizer AP21967, which promotes the dimerization of the FRB domain on Tr-HK1 and the FKBP domain on a mitochondria-targeted mCherry tag. (E) HEK293T cells expressing FRB-Tr-HK1-GFP (green) and mCherry-mito-FKBP (magenta) were treated with either vehicle (DMSO) or dimerizer AP21967. 10-15 minutes treatment with AP21967 was found to be sufficient to induce recruitment of HK1 to mitochondria. Scale bars represents 5µm. (F) Mitochondrial and glycolytic ATP production rates were calculated for HEK293T cells expressing either control Tr-HK1-GFP (localized in the cytoplasm), FRB-Tr-HK1-GFP (chemically-inducible cytoplasmic wild-type HK1 initially in the cytoplasm but recruitable to mitochondria), or FRB-Tr-HK1-T259A-GFP (chemically-inducible cytoplasmic HK1 with O-GlcNAc site mutation initially in the cytoplasm but recruitable to mitochondria), with or without 15-17 minutes AP21967 treatments. n=11–18 wells, three independent experiments (All ATP measurement values are shown as mean ± SEM, one-way ANOVA with post hoc Kruskal-Wallis multiple comparison test). (G) Schematic diagram illustrating glycolysis, highlighting the rate-limiting glycolytic enzymes and substrates. The two steps of glycolysis that generate ATP are indicated by red arrows. (H and I) Analysis of glycolytic enzymes in mitochondrial and cytoplasmic fractions from HEK293T cells. (H) Mitochondrial and cytoplasmic fractions, isolated from HEK293T cells treated overnight with either the OGA inhibitor Thiamet-G to enrich O-GlcNAcylation or vehicle (DMSO), were subjected to SDS-PAGE and analyzed by Western blotting using antibodies against HK1, Pyruvate Kinase M2 (PKM2), Aldolase A (Aldo A), Phosphoglycerate kinase (PGK), ATP5B (mitochondrial marker), and tubulin (cytoplasmic marker). (I) Quantification of the total intensity of glycolytic enzymes in mitochondrial, cytoplasmic fractions, and total cell lysates (input), normalized to the intensity of ATP5B (for mitochondrial fractions) and tubulin (for cytoplasmic fractions and input). Data are presented as mean ± SEM from n = 3 independent experiments (unpaired t-test). See also Figure S5.

To assess the functional consequences of O-GlcNAcylation on HK1 activity and cellular metabolism, we examined glycolytic and mitochondrial oxidative phosphorylation (OXPHOS) ATP production rates using respirometry measurements in HEK293T cells. We found that upregulating O-GlcNAcylation levels, either by OGA inhibition or by expressing WT-HK1 with Thiamet-G treatment, resulted in increased glycolytic and mitochondrial OXPHOS ATP production rates (Figure 5C). In contrast, cells expressing the T259A-HK1 mutant showed diminished ATP production rates, indicating the dependence of HK1 activity on O-GlcNAcylation. Moreover, the elevation in oxygen consumption rates (OCR) and extracellular acidification rates (ECAR) observed with O-GlcNAcylation upregulation was consistent with enhanced metabolic activity (Figure S5F). Notably, these effects were specific to O-GlcNAcylation status and not influenced by changes in endogenous HK1, WT-HK1, or T259A-HK1 expression levels (Figure S5G-I). These findings highlight the role of O-GlcNAcylation in modulating HK1-mediated ATP production rates through both glycolytic and mitochondrial pathways.

To rescue the impaired functional activity of the T259A-HK1, we employed a chemically induced protein dimerization strategy to promote mitochondrial recruitment. We utilized a dual-tagged construct, where truncated cytoplasmic HK1 (Tr-HK) was tagged with the FKBP-rapamycin-binding (FRB) domain at its N-terminus and GFP at its C-terminus. By co-expressing FRB-Tr-HK1-GFP and mCherry-mito-FKBP (FK506 binding protein domain), we could selectively recruit cytoplasmic HK1 to the mitochondria upon treatment with the dimerizer AP21967, as confirmed through time-lapse imaging (Figure 5D and 5E). To evaluate the functional rescue of T259A-HK1 activity, we measured ATP production rates in cells where T259A-HK1 was targeted to the mitochondrial outer membrane with the dimerizer. We observed a significant increase in both glycolytic and mitochondrial OXPHOS ATP production rates after recruiting cytoplasmic T259A-HK1 to the mitochondria (Figure 5F). This phenotypic rescue indicates that the impaired enzymatic activity of T259A-HK1 can be restored by targeted localization to the mitochondria, leading to enhanced ATP production rates. These results further highlight the significance of O-GlcNAcylation in modulating HK1 activity by regulating its mitochondrial localization.

### Hexokinase 1 O-GlcNAcylation promotes glycolytic metabolon formation on mitochondria

Enzymes involved in glycolysis are generally dispersed throughout the cytoplasm in cells^22^. However, under certain conditions such as hypoxia ^23-26^, these enzymes can compartmentalize to form metabolons, specialized microenvironments that enhance the transfer of metabolites from one enzyme to another^27,28^. This process enhances the efficiency of metabolic pathways and increases the overall metabolic output ^29^. Given this knowledge, we sought to investigate the potential role of O-GlcNAcylation in recruiting other glycolytic enzymes to mitochondria to form metabolon for metabolic efficiency (Figure 5G). We upregulated O-GlcNAcylation levels in HEK239T cells by ectopically expressing OGT and inhibiting OGA using Thiamet-G. We isolated whole cell lysate (Input), mitochondrial, and cytoplasmic fractions, which were then resolved on SDS-PAGE gels. We probed the gels with antibodies against endogenous HK1, pyruvate kinase M2 (PKM2), aldolase A (Aldo A), phosphoglycerate kinase (PGK), ATP5B (a mitochondrial marker), and tubulin alpha-4A (a cytoplasmic marker). Interestingly, our results indicated that enhanced O-GlcNAcylation not only increased the amount of HK1 in the mitochondria, but also augmented the levels of PKM2 and Aldo A on the mitochondria, a process that was parallel with their decrease in the cytoplasmic fraction (Figure 5H and 5I). However, the increased O-GlcNAcylation levels did not appear to significantly alter the localization of PGK (Figure 5H and 5I) in HEK293T cells.

We extended our analysis to cultured primary cortical neurons, probing for the presence of glycolytic enzymes in mitochondrial fractions under conditions of upregulated O-GlcNAcylation. Intriguingly, a significant portion of glycolytic enzymes are associated with purified mitochondria from cortical neurons. To account for the impact of upregulated O-GlcNAcylation, we normalized mitochondrial and cytoplasmic enzyme levels to the baseline. This showed that upregulated O-GlcNAcylation increased the mitochondrial localization of HK1, Aldo A, PGK, neuron-specific enolase (NSE), triosephosphate isomerase (TPI), and PKM in cortical neurons, while decreasing their presence in the cytoplasm (Figure S5J and S5K). Our results suggest that O-GlcNAcylation not only intensifies HK1 activity but also promotes the recruitment of rate-limiting glycolytic enzymes to mitochondria, resulting in increased ATP production rates. Collectively, our findings highlight HK1 O-GlcNAcylation’s crucial role in promoting the assembly of a glycolytic metabolon on mitochondria. Overall, our results demonstrate that O-GlcNAcylation plays a crucial role in facilitating the formation of a glycolytic metabolon on the mitochondria, where key enzymes involved in glycolysis are recruited, resulting in enhanced ATP production. This mechanism provides a new understanding of the coordination between glycolytic and mitochondrial metabolism and highlights the significance of O-GlcNAcylation in regulating cellular energy metabolism.

### O-GlcNAcylation Stabilizes HK1 on Mitochondria

Mitochondrial localization of HK1 is known to be modulated by G6P concentrations; elevated G6P concentrations can lead to the release of HK1, which acts as an inhibitory feedback mechanism^30^. We hypothesized that O-GlcNAcylation stabilizes HK1 on the mitochondria by counteracting this G6P-induced feedback loop. To test our hypothesis, we combined computational simulations and experimental assays. First, we utilized steered molecular dynamics (MD) simulations to understand the mechanism of G6P binding and unbinding from the N-terminal pocket of HK1. Since traditional MD simulations did not fully capture this process, we turned to steered MD simulations. These simulations demonstrated that the average work required to unbind G6P from the pocket was 13-fold lower for O-GlcNAc-modified HK1 compared to WT-HK1 (Figure 6A). It suggests that O-GlcNAcylation of the binding pocket reduces the affinity to G6P. To further corroborate our findings, we performed alchemical free energy perturbation calculations, which indicated a shift in the dissociation constant of G6P from 6.9 nM to 12 mM upon O-GlcNAcylation of the N-terminal binding pocket. The model was further supported by dynamic network analysis (Figure 6B and 6C).

**Figure 6.**
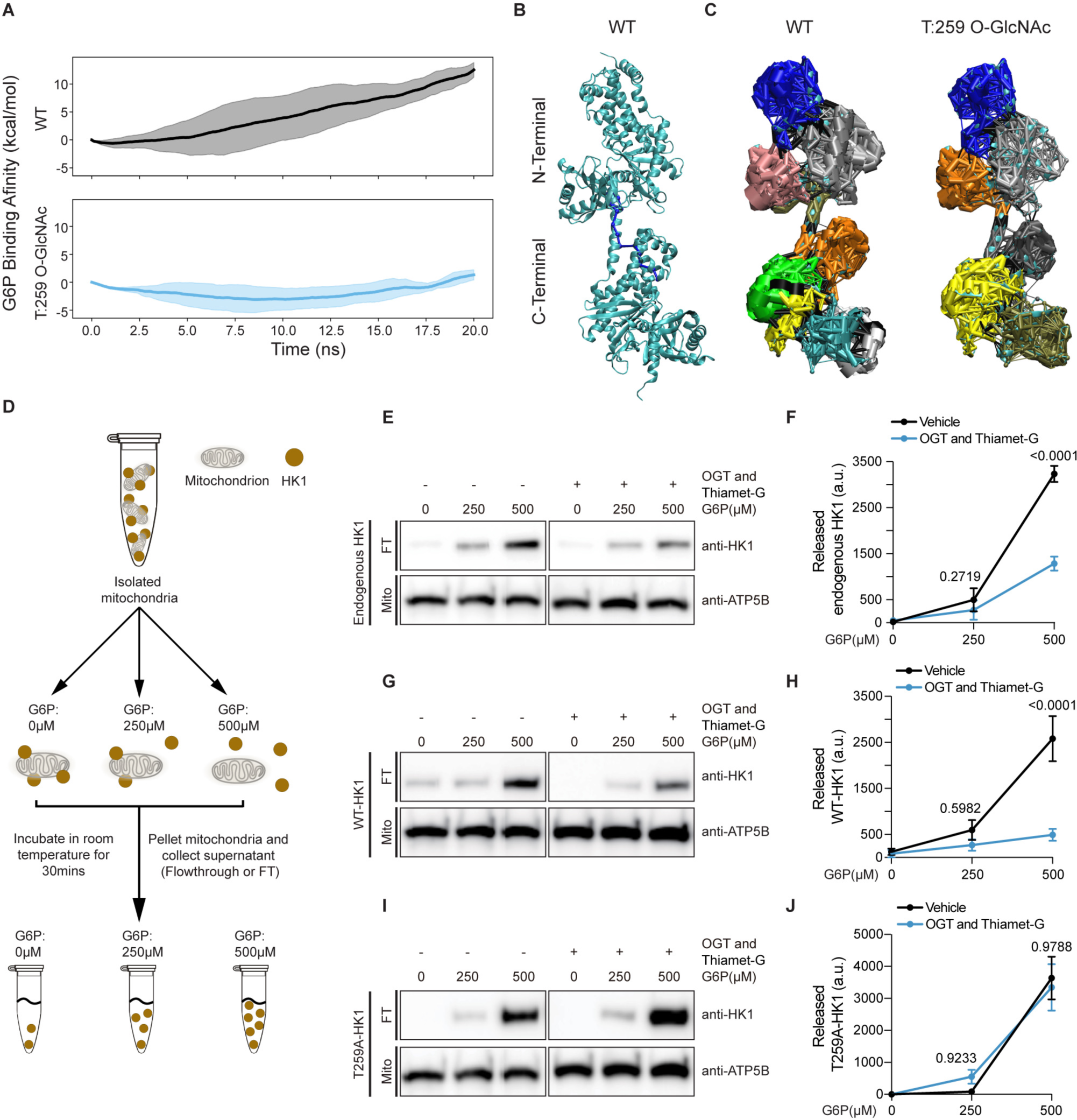
O-GlcNAcylation Modifies G6P Affinity and Stabilizes Hexokinase 1 on Mitochondria. (A) The average accumulated work profiles computed from steered MD simulations, in which Glucose-6-phosphate (G6P) was pushed out of the N-terminal binding pocket. The line represents the average accumulated work over 20 steered MD simulations, and the shading indicates one standard deviation from the mean accumulated work. (B) Optimal pathway of information transfer from the N-terminal Ser:449 to the C-terminal T:784 for the unmodified HK1. Dark blue nodes (spheres representing Cα atoms) and edges (cylinders representing allosteric interactions) illustrate the optimal pathway of allosteric interactions. (C) The community substructure for the network of allosteric interactions. Nodes within a community (single color) represent a network of interactions with more frequent and stronger connections with nodes in a community than with nodes outside of that community. Black edges represent connections between communities. (D) Schematic diagram illustrating the G6P titration assay for mitochondrial fractions. Isolated mitochondria, containing HK1, were exposed to three different concentrations of G6P for 30 minutes at room temperature. Following incubation, mitochondria and the supernatant, which contains the released HK1, were separated by centrifugation. Both the pellet (mitochondria) and supernatant (flowthrough containing released HK1) were collected and subsequently analyzed. (E-J) Mitochondrial fractions were prepared from HEK239T cells expressing control vector (E and F), eGFP-tagged WT (G and H) or T259A HK1 (I and J) with or without OGT co-expression and overnight Thiamet-G treatment. Flowthrough (FT) fractions were collected as described in (D) following 0, 250 and 500μM G6P treatments of the mitochondrial pellet (Mito). (E,G,I) Samples were then separated by SDS gel electrophoresis, and probed with anti-HK1, anti-ATPβ (mitochondrial marker) antibodies. (F,H,J) The total intensity of HK1 band was quantified for each condition. n = 3 independent experiments (All values are shown as mean ± SEM; Mann-Whitney *U* test). See also Figure S6.

To further investigate whether O-GlcNAcylation of HK1 strengthens its mitochondrial association by counteracting G6P-dependent feedback inhibition, we isolated mitochondria from HEK293T cells expressing endogenous HK1, WT-HK1-GFP or T259A-HK1-GFP. We then evenly distributed these mitochondrial fractions into three tubes containing 0, 250 and 500μM G6P (Figure 6D), concentrations that are within the physiological range for G6P^21,31^. After a 30-minute incubation at room temperature, mitochondria (Mito) were pelleted, separating them from the flow-through (FT) that contains proteins released from the mitochondria (Figure 6D). We observed that as G6P concentrations increased, more HK1 was released from the mitochondria (Figure 6E and 6F). Comparing the released HK1 from mitochondria at baseline and upregulated O-GlcNAcylation levels, we observed that increased O-GlcNAcylation resulted in less HK1 being released from mitochondria at the same G6P concentration (Figure 6E-H). We found up-regulating O-GlcNAcylation levels would stabilize WT-HK1 on mitochondria via counteracting G6P negative feedback (Figure 6G and 6H), but not for T259A-HK1 (Figure 6I and 6J). Thus O-GlcNAcylation is necessary for protecting HK1 from being released from mitochondria via G6P feedback inhibition.

Building on these molecular insights, we next delved into the broader cellular context to understand the implications of HK1 T259 O-GlcNAcylation on glycolytic metabolon formation. We analyzed the positioning of other glycolytic enzymes in HEK239T cells after expressing either WT-HK1 or T259A-HK1 from mitochondrial and cytoplasmic fractions. When we enhanced O-GlcNAcylation levels in cells overexpressing WT-HK1, we observed increased mitochondrial localization of WT-HK1, along with other glycolytic enzymes, such as Aldo A, PGK, and PKM2. At the same time, their cytoplasmic levels declined without a change in overall expression levels (Figure S6A and S6B). However, in cells overexpressing T259A-HK1 with increased O-GlcNAcylation levels, we did not detect a significant increase in mitochondrial localization of T259A-HK1, Aldo A, PGK, and PKM2. The cytoplasmic levels of these enzymes remained largely unchanged, as did their total expression levels (Figure S6C and S6D). These results further emphasize the specific role of HK1 O-GlcNAcylation in promoting the recruitment and localization of glycolytic enzymes on mitochondria, supporting the formation of the metabolon for efficient ATP production.

### Hexokinase 1 O-GlcNAcylation is essential for synaptic vesicle recycling

Given the role of HK1 O-GlcNAcylation in cellular metabolism, we next questioned how this modification might impact the high-energy demands of neurons. Neuronal activity and synaptic vesicle recycling heavily rely on ATP produced at presynaptic sites^32,33^. To investigate the impact of HK1 O-GlcNAcylation on neuronal function, we conducted live-cell imaging in rat hippocampal neurons. These neurons expressed either shRNA against HK1, shRNA-resistant WT-HK1-BFP, or T259A-HK1-BFP (O-GlcNAc silent HK1), along with mCherry as an axon filler and vGLUT1-pH, a pH-sensitive fluorescent probe attached to vesicular glutamate transporter (vGLUT1). We aimed to measure the recycling kinetics of presynaptic vesicles in response to neuronal activity and evaluate the role of HK1 O-GlcNAcylation in presynaptic function (Figure 7A and 7B). Under baseline conditions, neurons displayed spontaneous activity, with some vGLUT1-pHluorin on the axonal surface. Neuronal activation with 100 action potentials (APs) at 10 Hz significantly enhanced the fluorescence intensity of vGLUT1-pH, indicating the release of synaptic vesicles. The fluorescence signal quickly returned to baseline levels, reflecting the vesicle retrieval. The addition of NH4Cl at the end of the experiment allowed visualization of all presynaptic vesicles containing vGLUT1-pH (Figure 7C and 7D). Comparing neurons expressing with WT-HK1 and T259A-HK1 revealed lower baseline vGLUT1-pH signals in neurons expressing the O-GlcNAc silent T259A-HK1 (Figure 7D and 7E), independent of protein expression levels (Figure S7A-C). After applying 100 APs at 10 Hz, neurons expressing T259A-HK1 exhibited significantly lower vGLUT1-pH fluorescence intensity compared to WT-HK1 expressing neurons. Notably, there was no difference in the number of presynaptic release sites containing vGLUT1-pH between WT-HK1 and T259A-HK1-transfected neurons when treated with NH4Cl (Figure 7C). These results suggest that HK1 O-GlcNAcylation plays a crucial role in regulating energy-dependent pre-synaptic vesicle docking, formation of the readily releasable pool and retrieval. To assess the impact of HK1 O-GlcNAcylation on Ca^2+^ levels, we used GCaMP6s, a genetically encoded fluorescent Ca^2+^ indicator, in place of vGLUT1-pH. We did not observe significant differences in Ca^2+^ spikes in response to 100 APs at 10 Hz between WT-HK1 and T259A-HK1-expressing neurons (Figure S7D-I). These results indicate that the regulation of pre-synaptic vesicle recycling by HK1 O-GlcNAcylation is independent of Ca^2+^ flux.

**Figure 7.**
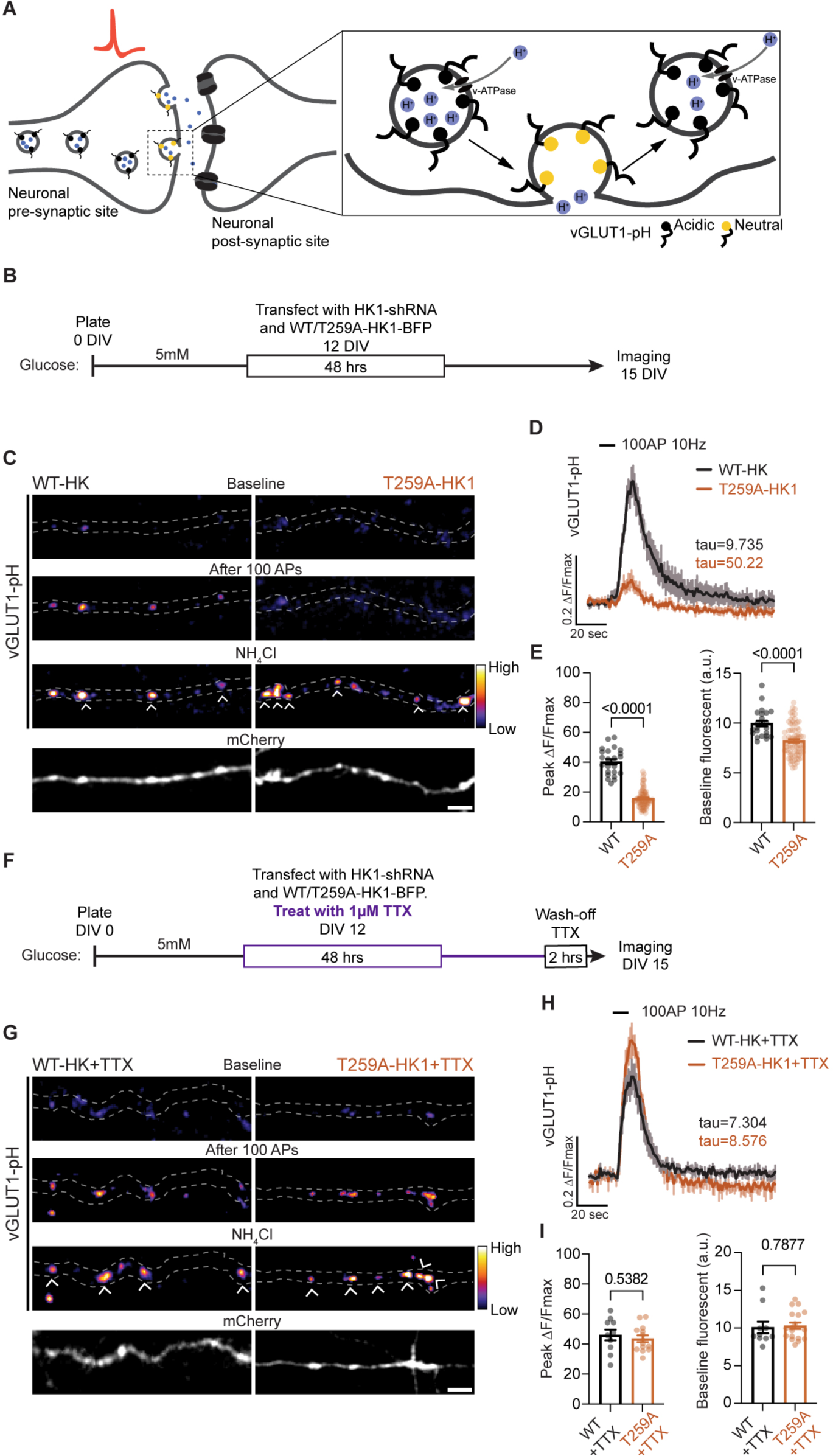
Presynaptic function relies on Hexokinase 1 O-GlcNAcylation. (A) Schematic of the pHluorin-tagged vesicular glutamate transporter 1 (vGLUT1-pH) located at the presynaptic release site of a neuron. The pHluorin protein, conjugated to the luminal domain of vGLUT1, exhibits fluorescence quenching at the acidic pH (∼5.5) inside synaptic vesicles. Upon neuronal stimulation and vesicle fusion to the plasma membrane, the luminal tag is exposed to the extracellular pH, leading to a significant increase in fluorescence. Post-endocytosis, pHluorin fluorescence is quenched again as the vesicle lumen becomes acidic by the activity of the vacuolar ATPase (v-ATPase). (B) The experimental design outlines the timeline for plating, transfection, and imaging of cultured rat hippocampal neurons. (C) Hippocampal neurons expressing either WT or T259A-HK1-BFP with vGLUT1-pH were electrically stimulated with 100 APs at 10 Hz. Representative images display vGLUT1-pH (pseudocolor, fire) and mCherry cell filler (gray) before and after stimulation in neuronal axons expressing either WT or T259A-HK1-BFP. Neutralization of vGLUT1-pH vesicles with NH4Cl treatment reveals total axonal vesicle pool. Scale bar represent 5µm. (D) Average trace of vGLUT1-pH with 100 APs 10Hz stimulation in WT-HK1 (black) or T259A-HK1 (orange) expressing neurons. ΔF values were normalized to maximal ΔF obtained from NH4Cl treatment. All values are shown as mean ± SEM. n = 10-11 neurons and 91-205 presynaptic boutons, four independent experiments. (E) Baseline and maximal (after electrical stimulation) vGLUT1-pH ΔF/F values. (F) The experimental design outlines the timeline for plating, transfection, Tetrodotoxin (TTX) treatment and imaging of cultured rat hippocampal neurons. (G) WT/T259A-HK1-BFP and vGLUT1-pH were expressed in hippocampal neuron. Following transfection, 1µM TTX was added to neuronal culture. Two hours before imaging, TTX was washed-off as shown in (F). Neurons were electrically stimulated with 100 APs 10Hz. Images showing vGLUT1-pH (pseudo-color, fire) and the cell filler mCherry (gray) before and after stimulation with WT-HK1 or T259A-HK1-BFP expressing neurons. Neutralization of vGLUT1-pH vesicles with NH4Cl reveals total axonal vesicle pool. (H) Average trace of vGLUT1-pH with 100 APs 10Hz stimulation in WT-HK1 (black) or T259A-HK1 (orange) expressing neurons, previously TTX treated. ΔF values were normalized to maximal ΔF obtained from NH4Cl treatment. Error bars represent SEM. n = 8-9 neurons and 20-55 presynaptic boutons from four independent experiments. (I) Baseline and maximal (after electrical stimulation) vGLUT1-pH ΔF/F values. All values are shown as mean ± SEM (unpaired t-test). See also Figure S7.

We further examined whether the observed reduction in pre-synaptic vesicle recycling in neurons expressing T259A-HK1 was due to an energy deficiency. We transfected neurons with either WT-HK1 or T259A-HK1-BFP, along with vGLUT1-pH. Following transfection, we added 1 µM tetrodotoxin (TTX), a blocker of voltage-gated sodium channels, to suppress spontaneous neuronal activity and conserve ATP levels. Two hours before imaging, TTX was washed off, allowing for the recovery of baseline pre-synaptic vesicle recycling (Figure 7F). By preserving ATP levels through TTX treatment, neurons expressing T259A-HK1 exhibited a restoration of pre-synaptic vesicle recycling comparable to WT-HK1 expressing neurons (Figure 7G-I). Furthermore, following one round of electrical stimulation, pre-synaptic vesicle recycling was successfully rescued in T259A-HK1 expressing neurons, demonstrating a similar performance to that of WT-HK1 expressing neurons (Figure 7G-I). Our findings highlight the essential role of O-GlcNAcylation of HK1 in addressing localized energy demands in neurons. Specifically, this post-translational modification of HK1 is indispensable for proper vesicle recycling at presynaptic terminals, a compartmentalized ATP-intensive process.

## DISCUSSION

Cells operate within a finely-tuned signaling network where metabolism, efficient energy management, and nutrient sensing work in unison to ensure optimal cellular function and viability under ever-changing nutrient environment. Among these intricate mechanisms, O-GlcNAcylation recently emerged as a pivotal nutrient-sensing posttranslational modification (PTM)^34^. Our study provides critical insights into the regulatory role of O-GlcNAcylation in cellular metabolism and energy homeostasis, specifically through its modulation of the positioning and activity of Hexokinase 1 (HK1), an enzyme central to glucose metabolism^21,35^. Our findings demonstrate a dynamic regulatory feedback system where glucose levels, through the mediation of O-GlcNAcylation, orchestrate the positioning of HK1 to mitochondria. This regulatory mechanism involves the O-GlcNAcylation of Threonine 259 on HK1, which we found to be crucial for HK1 localization and activity. Moreover, our study provides evidence of how HK1 O-GlcNAcylation promotes the formation of a glycolytic metabolon on the mitochondrial outer membrane. Metabolon formation increases the efficiency of glucose metabolism and ATP generation, crucial for high energy demanding areas such as the presynaptic sites.

The O-GlcNAcylation of HK1 is tightly correlated with its positioning on mitochondria, whether the level was manipulated by overexpression of OGT, pharmacological inhibition of OGT and OGA, or by nanobody-mediated targeting of O-GlcNAc cycling (Figure 2 and 3). OGT post translationally modifies over 8,000 proteins, including several glycolytic enzymes such as PKM2^36^, PKF^37^ and PGK1^38^, all of which potentially impact cellular metabolism. However, by mapping and mutating the only site of O-GlcNAcylation on HK1 (T259A-HK1), we demonstrated that HK1 is a pivotal substrate through which OGT can enhance ATP synthesis rate (Figure 4 and 5). While T259A-HK1 does not alter HK1 enzymatic activity, we found that it selectively prevents mitochondrial recruitment of HK1 and the formation of glycolytic metabolon on mitochondria (Figure 5 and S6). These findings reveal a novel role for HK1 in regulating cellular energy metabolism through glucose and O-GlcNAcylation-dependent mitochondrial localization.

Cells maintain metabolic homeostasis through the precise sensing and regulation of glucose fluctuations. Within this complex regulatory landscape, O-GlcNAcylation emerges as a crucial modulator^34^. In contrast to other forms of protein glycosylation, certain O-GlcNAcylation sites are dynamically regulated, undergoing rapid addition or removal in response to changes in the cellular state, such as excitatory neuronal stimuli^39,40^. Conversely, others sites primarily fulfill structural roles and exhibit slower turnover^41^. Only dynamic and substoichiometric O-GlcNAcylation is highly responsive to glucose concentrations, influx through HBP and OGA inhibition (with PUGNAC or Thiamet-G) in neurons and other cell types^12,13,42^. Our study shows that dynamic changes of HK1 O-GlcNAcylation on highly conserved T259 site allows metabolic sensing (Figure 2, 3 and 4). The widely expressed hexokinase isoform HK2, which lacks the T259 O-GlcNAcylation site, does not exhibit similar regulation in response to genetic or pharmacological OGT and OGA manipulations (Figure S3). Notably, HK1 contains an N-terminal regulatory domain instead of two catalytic domains found in HK2. Our study reveals that the affinity of G6P to the N-terminal domain binding pocket is markedly reduced upon T259 O-GlcNAcylation (Figure 6). Previous studies have suggested an allosteric communication from the non-catalytic N-terminal domain of HK1 to the catalytic C-terminal binding pocket, wherein binding G6P at the N-terminal promotes a conformational change at the C-terminal^18^. Hence, we postulate that O-GlcNAcylation of HK1 averts allosteric regulation by the N-terminal binding pocket on the C-terminal catalytic pocket, ensuring HK1 remains catalytically active despite the presence of G6P. Our observations indicate that the binding of G6P to the N-terminal pocket significantly reduces the dipole moment of HK1, causing its N-terminal domain to have a less preferential orientation in the electrostatic field created by the negatively charged outer mitochondrial membrane. The N-terminal domain of HK1 contains the mitochondrial targeting domain^11^. Potentially, an increase in the molecular dipole orients the N-terminal half of HK1 toward the mitochondrial outer membrane allowing mitochondrial targeting domain to insert itself into the membrane. In conclusion, our study offers a rare and compelling demonstration of how a singular post-translational modification, O-GlcNAcylation, profoundly influences the structure and function of HK1, unraveling critical insights into the complex and finely tuned metabolic regulation of this pacemaker enzyme.

Glucose regulates the localization of mitochondria in neurons through OGT/O-GlcNAcylation^12,13^, allowing the cell to adjust to spatial heterogeneity in glucose concentration and directing mitochondria to the glucose-rich areas^43,44^. We have now revealed another layer of regulation and elucidated that OGT/O-GlcNAcylation, by altering the HK1 structure and mitochondrial localization, also regulates the spatial organization of glycolytic enzymes, promoting “metabolon” formation on mitochondria to bolster metabolic efficacy (Figure 5 and S5). Assembling enzymes with their subsequent substrates in close proximity increases reaction rates through substrate channeling^45,46^. Substrate channels between glycolytic enzymes located on the mitochondrial surface in neurons could be a regulatory mechanism to ensure that adequate pyruvate is supplied into mitochondria when respiratory demand increases, rather than allowing glycolytic intermediates to be diffused or used for an alternative metabolic pathway. Conversely, the release of hexokinase into the cytoplasm would result in more spatially uniform glucose consumption, negating the metabolic enhancement achieved through mitochondrial localization. The formation of cytoplasmic glycolytic metabolon was previously observed in multiple cell types including in yeast^24,26,27,47,48^, cancer cells^28,49^, *Trypanosoma*^50^, *Drosophila* flight muscle^51,52^, also in response to hypoxia in *C. elegans* neurons^23,25^. Here we provide a molecular mechanism that promotes glycolytic metabolon formation on mitochondria via post-translational modification of HK1. This mechanism amplifies both glycolytic and mitochondrial ATP synthesis rate. Importantly, this regulation via O-GlcNAcylation of HK1 is dynamic, leading to the redistribution of glycolytic enzymes within minutes. This dynamic rearrangement underscores the existence of a spatially and temporally synchronized system for glucose uptake, processing, and ATP production, reinforcing cellular energy homeostasis.

In the context of neurons, a cell type with substantial and localized energy needs, coupling glycolysis to mitochondria via the O-GlcNAcylation mechanism becomes particularly crucial. Despite their significant energy demands, neurons lack stored ATP and predominantly metabolize glucose through glycolysis to meet acute energy requirements^32,33,53-55^. Synaptic activity and action potential firing also create a rapid and highly dynamic energetic demand for neurons. In response to periods of high activity neurons increase in both surface glucose transporter levels and glucose uptake—aligning with the intensified demand for ATP^32,33^. This dynamic is underscored by findings that synaptic activity boosts glucose consumption by a median of 1.9-fold^56^, also promotes O-GlcNAcylation^40^. Intriguingly, a study indicates that in a resting adult mouse brain, hexokinase operates at a mere 3% of its maximum potential, suggesting an available 33-fold upswing in activity^36^. Here we provide a molecular mechanism that the O-GlcNAcylation of HK1 would increase its activity and ATP production at pre-synaptic sites, thus supply the high energy demand during intense synaptic activity (Figure 7). Dysregulation of HK1 O-GlcNAcylation, resulting from the introduction of T259A mutation, likely impairs pre-synaptic vesicle recycling due to ATP insufficiency. The docking, release and retrieval of synaptic vesicles at the active zone are ATP-dependent processes^32^. The synaptic vesicles themselves also demand a significant amount of ATP, especially given their reliance on vesicular V-ATPase activity for filling vesicles with neurotransmitters^57^. Given this, it is consistent that by silencing neurons using TTX, which blocks spontaneous firing and conserves ATP levels, rescues the synaptic vesicle recycling defects observed with the T259A mutation (Figure 7).

Neurological disease including Alzheimer’s disease (AD)^58,59^, depression^60^ and schizophrenia^61^ have been correlated with metabolic dysfunction, such as the dissociation of HK1 from mitochondria and reduced glucose metabolism^62-64^. The prevalence of O-GlcNAcylation within the brain, coupled with the changes in glucose metabolism in AD brains, suggests that O-GlcNAcylation may play important roles in AD progression^65-69^. The precise regulation of glucose sensing, uptake, and processing is fundamentals to cellular metabolic adaptability. In this study, we elucidate how nutrient sensing influences HK1 O-GlcNAcylation and its interaction with neuronal mitochondria, optimizing metabolic efficiency through the PTM mechanism. Disruptions in this metabolic pathway correlate with various neurological disorders, suggesting a previously unidentified involvement in neuropathologies. By unraveling this essential feedback mechanism, which integrates synaptic activity with ATP synthesis, we set the stage for discovering new therapeutic targets in neuronal dysfunction.

## ACKNOWLEDGEMENTS

We gratefully acknowledge the invaluable contributions of the Pekkurnaz laboratory members, as well as the generous sharing of key instrument resources by Dr. Enfu Hui. We also extend our appreciation to the technical team members of the University of California San Diego BPMSF and Nikon Imaging Center for their expert assistance. This project was made possible by the support of a grant from the National Institutes of Health (NIH) to G.P. (R35GM128823), NIH (2T32GM007240) to S.B.Y., NIH (5T32GM133351) to A.A.A., NIH to M.H. (R01NS094219), University of California San Diego TRELS fellowship to A.Z., NIH (5T32NS007220) to V.L. and the San Diego IRACDA Scholars Program (K12GM068524) to R.S. We also acknowledge the Gordon and Betty Moore Foundation and the Chan Zuckerberg Initiative DAF, an advised fund of Silicon Valley Community Foundation (2020-222005) for their contributions to this project.

## AUTHOR CONTRIBUTIONS

Experiments were designed by G.P. and H.W., with technical contributions from H.W., R.S., M.L.M., A.Z., S.B.Y., V.L., M.J., and A.A.A. J.V. and A.S. contributed structural data and performed molecular simulations, while *C. elegans* experiments were carried out by Y.W. and

M.H. Mass spectrometry experiments and analysis were performed by M.G., and E.G. provided expert technical assistance for microscopy. The manuscript was written by H.W. and G.P., with input from all co-authors.

## DECLARATIONS OF INTERESTS

The authors declare no competing interests.

## METHODS

### EXPERIMENTAL MODEL AND SUBJECT DETAILS

#### Mice and Rat strains and maintenance

All animal experiments were conducted according to the NIH Guide for the Care and Use of Experimental Animals and approved by the University of California San Diego Animal Care and Use Committee. C57BL/6J strain wild-type male and female mice (Jackson Laboratory) were used for fasting/refeeding studies. Mice were housed at 22-24°C in a room using a 12-hour light/dark cycle with *ad libitum* access to standard chow diet and water. For experiments involving fasting and refeeding, food was removed, then replaced for 24 hours. Mouse blood glucose levels were measured using Accu-Chek® Aviva Plus Glucose Monitor. Food intake, body weight and blood glucose levels were monitored weekly throughout the experiments as previously described^12^. Sprague-Dawley strain wild-type rats (Envigo) were used for primary neuron cultures. Timed pregnant female rats (embryonic days 13-16) were singly housed in a room with a 12-hour light/dark cycle with *ad libitum* food, water access, and environmental enrichment. Primary hippocampal and cortical neuron cultures were generated as described in the section below.

#### *Caenorhabditis elegans* strains and maintenance

All strains were maintained at 20°C on nematode growth medium plates seeded with *E. coli* OP50 as previously described ^70^. Hermaphrodites were used for all experiments, and males were only used for crossing. To label mitochondria in cholinergic motor neuron DA9, mitochondria matrix localization signal (as in pPD96.32) was fused with TagRFP expressed as extrachromosomal arrays. The microinjection mix contained Pitr-1pB::mito::TagRFP::let858 3’UTR (pYW217) at 30 ng/μl to label the mitochondria in DA9 neuron, Podr-1::gfp at 20 ng/μl to label the AWB/C neurons as the co-injection marker, and Promega 1 kb DNA ladder as the filler. The hxk-1::7xgfp11 knock-in was generated by CRISPR/cas9 using preassembled Cas9 ribonucleoprotein complex ^71^ (Figure S2F). Briefly, the injection mix contained 0.25 μg/μl S.pyogenes Cas9 (IDT), 0.1 μg/μl tracrRNA (IDT), 0.056 μg/μl crRNA, 25 ng/μl pre-melted double-stranded DNA repair template and 40 ng/μl PRF4 (rol-6 (su1006)) plasmid. The mix was injected into a strain that expresses GFP1-10 fragment in the DA9 neuron and another neuron in the ventral cord^72^. Non-roller F1 progeny from P0 roller jackpot plates were screened by single worm PCR. Homozygotes from the F2 generation were verified by Sanger sequencing. The repair template contained 35 bp homology arms on both ends and a GSGGGG liker between hxk-1 and 7xgfp11. The repair template was synthesized as a Gblock by IDT, amplified by PCR, gel purified and further cleaned up with the MinElute PCR purification kit.

#### Primary neuronal culture

Primary neuron cultures were established from rat embryos (E18) as previously described ^73^. Hippocampal and cortical neurons were plated at a density of 5-7 × 10^4^ cells/cm^2^ on round (12mm diameter) coverslips (Carolina Biological Supplies) for imaging or on 6-well plates at a 1-2 × 10^5^ cells/cm^2^ for biochemical assays. Prior to plating, coverslips and plates were coated with 20 µg/mL Poly-L-Lysine and 3.5 µg/mL Laminin overnight at room temperature. Primary neuron cultures were maintained in Neurobasal medium containing 5mM glucose, supplemented with B27, GlutaMAX, and penicillin/streptomycin unless modified as specified. Glucose levels in neuron culture media were monitored every 24 hours to maintain concentration of 5mM glucose. For neuron culture media containing 1mM glucose, glucose levels were monitored every 12 hours. Each independent experiment was performed by the preparation of new primary neuron cultures. For biochemical assays, cortical neurons were treated with 1 µM Cytarabine (Ara-C) at 3 days in vitro (DIV) to prevent glia proliferation for 2 days. The Ara-C-containing medium was replaced with fresh neuron maintenance medium at 5 DIV. The primary neuron cultures were maintained for 11-15 DIV by replacing one-third of the culture medium with a fresh medium every three days. When indicated, hippocampal and cortical neurons were treated with Thiamet-G at a final concentration of 5 µM for 16–18 hours. Primary neuron cultures were transfected with indicated plasmid DNA constructs using Lipofectamine 2000 for each experiment, and imaged 2-3 days later. For each experiment, co-transfection efficiency of 80-90% (up to four plasmids) was confirmed retrospectively.

#### Cell line culture

Human embryonic kidney 293T (HEK293T), monkey kidney COS-7 and Neuro-2a cell lines were used to perform indicated experiments. The cell lines were obtained from American Type Culture Collection (ATCC), and were tested every 6 months for mycoplasma contamination using LookOut^®^ Mycoplasma PCR Detection Kit. To achieve consistency in cellular metabolic properties, the cell lines were used no more than 15 passages. HEK293T cells were CRISPR-edited to endogenously tag HK1, as described in the section below. Both wild-type and CRISPR-edited HEK293T cells were maintained in DMEM containing 5mM glucose, supplemented with Penicillin (100 U/ml)/Streptomycin (100 μg/ml)/L-glutamine (2mM), 10% FBS at 37°C in a 95% air/5% CO2 humidified incubator. For biochemistry experiments and respirometry measurements, HEK293T cells were plated on 6-well-plate at a density of 6.3x10^4^ cells/cm^2^. For microscopy experiments, HEK293T cells were plated on round (12mm diameter) coverslips at a density of 2.6x10^4^ cells/cm^2^. Plasmid DNA transfections were performed with calcium phosphate protocol ^74^ when HEK293T cells reached at least 30% confluency, using 0.5-1µg of DNA per well. When indicated, HEK239T cells were treated with Thiamet-G at a final concentration of 10 µM for 16–18 hours. COS-7 and Neuro-2a cell lines were maintained in DMEM containing 25mM glucose, supplemented with Penicillin (100 U/ml)/Streptomycin (100 μg/ml)/L-glutamine (2mM) and 10% FBS at 37°C in a 95% air/5% CO2 humidified incubator. For microscopy experiments, COS-7 cells were plated on round (12mm diameter) coverslips at a density of 2x10^4^ cells/well, and transfected with plasmid DNA (0.4-2.9 µg of DNA per well) confluency using calcium phosphate protocol ^74^ when they reached 30%. Neuro-2a cells were transfected with plasmid DNA (2.5 µg DNA per well) at 50% confluency using Lipofectamine™ 3000 transfection reagent in Opti-MEM medium. Cells were lysed for biochemical analysis or processed for microscopy 2-3 days after transfection, as indicated for each experiment.

### METHOD DETAILS

#### Plasmid Constructs

The following previously published or commercially available DNA constructs were used: vGLUT1-pHluorin^75^ (gift from Dr. Ghazal Ashrafi), mCherry^76^ (gift from Dr. Gentry Patrick), pRK5-Myc-OGA^77^ and nucleocytoplasmic OGT (ncOGT)^78^ (gift from Gerald Hart laboratory, Johns Hopkins and the NHLBI P01HL107153 Core C4), mCherry-mito-FKBP^79^ and pDsRed2-Mito^12^ (gift from Thomas Schwarz), pAAV.CAG.GCaMP6s.WPRE.SV40^80^ (gift from Douglas Kim and GENIE Project; Addgene plasmid 100844), FLHKI-pGFPN3, TrHKI-pGFPN3 and FLHKII-pGFPN3^11^ (gift from Hossein Ardehali; Addgene plasmid 21917, 21918 and 21920), (294) pcDNA3.1-myc-OGA(1-400) and (344)pcDNA3.1(+)-HA-nLaG6-(EAAAK)4-OGA(544-706)^15^ (gift from Christina Woo; Addgene plasmid 168095 and 168197). Additionally, shRNA plasmid against mouse HK1 and empty vector control were purchased from Sigma-Aldrich MISSION.

The following plasmids are generated in this study: T259A-HK1-GFP was PCR amplified from WT-HK1-GFP with the Threonine 259 to Alanine mutation via In-Fusion HD Cloning Kit. WT/T259A-HK1-BFP was generated from WT/T259A-HK1-GFP by excising GFP using ApaI and NotI digestion, and replacing it with gene block containing TagBFP. Furthermore, pEGFP-FRB*:Tr-hHK1 (Vector ID number VB181221-1086jen) and pEGFP-FRB*:Tr-T259A-hHK1 (The vector ID number VB181221-1158hep) used in this study was constructed by VectorBuilder. The vector ID numbers can be used to retrieve the detailed information.

#### CRISPR-Mediated EGFP Tagging of Endogenous Hexokinase 1 in HEK293T

To generate endogenously fluorescently tagged Hexokinase 1 (with EGFP) expressing HEK293T cells (HEK293T-EGFP-HK1), single guide RNAs (sgRNAs) were designed to target the region between exon 18 and 3’UTR of the human transcript HK1-202 (NM_000188). Candidate sgRNAs were designed by the CRISPR design tool from the Wellcome Sanger Institute. To confirm the on-target activity of the designed sgRNAs, Universal CRISPR Activity Assay (UCATM), a sgRNA activity detection system developed by Biocytogen, was used. The targeting donor vector used in this study contained a PuroDeltatk resistant cassette flanked by loxP sites in intron17 and a linker-EGFP cassette inserted before the TAA stop codon of the human HK1 gene. Thus, the fluorescent protein EGFP is fused to and under control of the endogenous HK1 promoter (Figure S3C). To introduce the sgRNA2 and targeting vector into HEK293T cell line, electroporation was used. After the selection of cells for drug resistance and performing PCR, mixed positive clones were further electroporated with a Cre expressing vector to remove the antibiotic resistant cassette. Subsequently, fourteen clones were picked and expanded in semisolid media. Finally, two homozygous clones were obtained by further genotyping characterization and PCR product sequencing.

#### *C.elegans* Imaging and Analysis

ST045849 was used to inhibit the O-GlcNAc Transferase (OGT) in *C. elegans* ^14^. A 20 mM stock solution was prepared in DMSO and added to NGM media to achieve a final concentration of 50 μM. Media was poured into 12-well plates at a volume of 2 ml per well. The plate was left to dry overnight at room temperature with a loose cover. The following day, 50 μl of concentrated OP50 bacteria containing 50 μM ST045849 was added to each well. After the plate had dried for approximately 30 minutes, L2 worms were transferred onto the plate immediately and incubated at 20°C. Worms at L4/adult transition stage was imaged the next day (approximately 24 hours later). For the control group, the same amount of DMSO was added to both NGM and OP50.

For imaging, L4 animals were immobilized with 50 mM Muscimol and mounted on 3% agarose pads. Animals oriented with the ventral side facing the coverslip were selected for imaging the cell body. Images were acquired on a VT-iSIM system (BioVision) built around a Leica Laser Safe DMi8 inverted scope with a HC PL APO 63x/1.40 OIL CS2 objective and an ORCA-Flash4.0 camera (Hamamatsu). Emission filters 525/50nm and 605/52 nm were used for 488 nm and 561 nm illumination, respectively. Z-stacks were acquired with 0.2 μm step size on an ASI-XYpZ Piezo stage. Images were taken sequentially for 488 nm and 561 nm at each Z position, and 400 ms exposure time was used for each channel. Image acquisition was controlled by the MetaMorph Advanced Confocal Acquisition Software Package. Maximum projections were generated in Fiji^81^ from microcopy data. To calculate the ratio of HXK-1::7xSplitGFP intensity on mitochondria versus the cytoplasm in the cell body, all images were blinded and randomized. Mitochondria were traced manually in the TagRFP channel, and average intensities from the GFP channel in areas with mitochondria or without mitochondria were recorded. The auto fluorescent signal from the surrounding area was selected as background and subtracted from the GFP intensity. The subtracted values were then used to calculate the mitochondrial and cytoplasmic (mito/cyto) ratio.

#### Brain Slice Immunohistochemistry

Immediately after the end of the fasting/refeeding experiments, mice were anesthetized by inhalation of isoflurane and perfused transcardially first with ice-cold phosphate-buffered saline (PBS, pH 7.4) containing heparin (19.5 U ml^-1^), followed by freshly prepared 4% paraformaldehyde (PFA) in PBS. Brains were extracted and post-fixed with 4%PFA solution overnight, rinsed with PBS, cryoprotected in 30% sucrose, and embedded with Tissue-Tek® O.C.T. Compound for cryosectioning. Brain sections of 20 µm thickness were obtained using a cryostat. The sections were placed in Mouse-on-Mouse blocking solution (1 drop in 2.5 ml of 1x PBS) for 30 mins at room temperature to reduce nonspecific binding. To further prevent nonspecific binding, the sections were incubated in blocking buffer (3% normal goat serum, 1% bovine serum albumin, 1% fish gelatin, 0.1% Triton X-100, 1X PBS, ddH2O) for 1 hour at room temperature. The sections were then incubated overnight at 4 °C with the following antibodies in staining solution (10% blocking buffer, 0.1% Triton X-100, 1X PBS, ddH2O): rabbit anti-HK1 (1:500), mouse anti-Pyruvate dehydrogenase (PDH) (1:500), chicken anti-NeuN (1:500). After washing with 1X PBS three times, the sections were incubated with secondary antibodies conjugated with Alexa Fluor™ 488, 568 and 647 (1:500 dilution for each) for 1 hour at room temperature. The sections were then washed twice with 1X PBS before incubating with DAPI 4’,6-diamidino-phenyindole (1:1000 in 1X PBS) for 10 mins at room temperature to label nuclei. Lastly, the sections were washed twice with 1X PBS and mounted on Superfrost™ Plus Microscope slides and #1.5 rectangle Platinum Line cover glass using VECTASHIELD^®^ Vibrance™ antifade mounting medium. Images were acquired using both Zeiss Axio Imager Z2 upright microscope with ZEN pro Elements, and the fluorophores were excited using the 385nm, 475nm, 555nm, 630nm lines of the six-line (385nm, 430nm, 475nm, 511nm, 555nm, 630nm), Plan-Apochromat objective of 20x/0.4 objective, the camera used was AxioCam 712 mono, pixel size 3.45 μm × 3.45 μm, 1,388 × 1,040 px, and Zeiss LSM 780 confocal laser scanning microscope, Plan-Apochromat 100x/1.40 Oil DIC M27 objective with the highly sensitive low dark noise PMTs (2x) and GaAsP (32x) array. Image acquisition settings were kept constant across different conditions and sections. The acquired images were analyzed using Fiji/ImageJ software^81^, and linear adjustments of brightness and contrast were made only for visualization purposes. To quantify colocalization between HK1 and mitochondria first all images were blinded and randomized, then Fiji/ImageJ’s COLOC2 plugin was used to calculate Manders’ coefficients^82^. Indicated regions from mice hippocampus were used for this analysis. The calculated values were extracted, and statistically analyzed using GraphPad Prism version 7.0 for Mac OS.

#### Immunocytochemistry

HEK293T and COS-7 cells were fixed with 4% PFA in 1X PBS for 10 mins at room temperature. Fixed cells were washed three times with 1X PBS, followed by blocking with 0.5% Saponin and 1% BSA in 1X PBS for 1hr at room temperature. The cells were then immunostained with primary antibodies at 4°C overnight in blocking buffer. The primary antibodies used were rabbit anti-GFP, mouse anti-GFP, rabbit anti-Tomm20, rat anti-HA, and chicken anti-myc (1:500 dilution for each). After washing cells three times with 1xPBS, they were incubated with secondary antibodies conjugated with Alexa Fluor™ dyes at 1:500 dilution for 1 hour at room temperature. The secondary antibodies used were goat anti-rabbit Alexa Fluor™ 488, goat anti-chicken Alexa Fluor™ 405, goat anti-mouse Alexa Fluor™ 488, goat anti-rabbit Alexa Fluor™ 568, and goat anti-rat Alexa Fluor™ 647. Following another three washes with 1xPBS, coverslips were mounted using DAPI Fluoromount-G. Images were acquired using a Zeiss LSM 780 confocal laser scanning microscope equipped with Plan-Apochromat 100x/1.40 Oil DIC M27 objective.

Immunocytochemistry of 11-15 DIV disassociated primary neuron cultures was performed as previously described^12^. Briefly, neurons were fixed with 4% PFA and 4% sucrose in 1X PBS for 10 mins at room temperature. The fixed neurons then washed three times in 1X PBS, followed by blocking in 1x GBD (10% goat serum, 1% bovine serum albumin, and 0.1% Triton X-100 in 1X PBS) for 1 hour at room temperature. Primary antibodies were diluted in 1X GBD and incubated with the neurons overnight at 4°C. Secondary antibodies were also diluted in 1X GBD and incubated with the neurons for 1 hour at room temperature. The following primary and secondary antibodies were used: rabbit anti-HK1 (Invitrogen, 1:500) or rabbit anti-HK1 (Cell Signaling, 1:250), goat anti-rabbit Alexa Fluor™ 647 (1:500). The coverslips were mounted using Fluoromount-G. Images were acquired using a Zeiss LSM 780 confocal laser scanning microscope equipped with C-Apochromat 40x/1.20 W Korr FCS M27 objective. The image acquisition settings were kept consistent across different conditions and coverslips for HEK293T, COS-7 and primary neuron immunostaining. The acquired images were analyzed using Fiji^81^, and only linear adjustments of brightness and contrast were made for visualization purposes. All experiments were performed at least three times, and representative images are shown. Quantification of staining intensity and colocalization was performed on Fiji/ImageJ, and statistical analysis was conducted using GraphPadPrism version 7.0 for Mac OS.

#### Mitochondria Isolation

Mitochondrial fractions were prepared from 11-15 DIV primary cortical neuron cultures or HEK293T cells, plated on 6-well plates, by homogenization in mitochondrial isolation buffer (MIB) (10 mM Tris-HCl (pH 7.4), 10 mM KCl, 250mM Sucrose, 1 mM EDTA (pH 8.0), 1x Protease Inhibitor Cocktail Set III EDTA Free, 0.1 mM phenylmethylsulfonyl fluoride (PMSF), 4 µM Thiamet-G and 2 mM Dithiothreitol (DTT)) and differential centrifugation. Each well containing cells (∼1.5 x 10^7^ cells/condition) were first washed with 1 ml ice cold 1X PBS and then incubated with 330 µl of freshly prepared MIB on ice for 10 mins with gentle agitation. Cells were detached with cell scraper and homogenized with 20-30 strokes using a tight-fitting B pestle in a 1 ml Dounce homogenizer. The homogenate was centrifuged at 700xg for 10 minutes at 4°C to pellet nuclei and large cell debris. 20 µl of the first supernatant (Input, which containing mitochondria) was saved for western blot analysis as whole cell lysate. The supernatant centrifuged again at 10,000xg for 10 minutes at 4°C to pellet the crude mitochondrial fraction. The second supernatant (cytoplasmic fraction) was collected and concentrated using 0.5 ml Centrifugal Filters, Ultracel-10K. 50% of mitochondrial fraction and 40% of cytoplastic fraction were then loaded in SDS-PAGE and analyzed by western blotting. For western blot analysis, samples were resuspended in 1xLaemmli sample buffer, and denatured at 95°C for 5 minutes before loading onto an SDS-PAGE. After separation, the proteins were transferred to nitrocellulose membranes and stained with primary/secondary antibodies as previously described^12^. Stripping buffer was used to reprobe the western blots with different antibodies, and blots were only reprobed after confirming the absence of the previous signal. Chemiluminescent detection with SuperSignal™ West Dura Extended Duration Substrate. For quantitative western blot measurements, image exposure times were optimized for the linear range of detection using Azure C600 Biosystem gel documentation system. All experiments were performed at least three times. The images were further analyzed using Fiji gel analyzer^81^, using only linear adjustments of brightness and contrast for visualization.

The following antibodies were used for probing blots to analyze mitochondrial and cytoplasmic fractions: rabbit anti-HK1 (Invitrogen) at 1:500, rabbit anti-PKM2 at 1:500, mouse anti-PKM at 1:250, rabbit anti-Aldolase A used at 1:250, mouse anti-Aldolase A at 1:500, mouse anti-PGK at 1:250, rabbit anti-GPI at 1:500, mouse anti-PFKM at 1:500, mouse anti-GAPDH at 1:1,000, mouse anti-PGAM1 at 1:500, mouse anti-NSE at 1:500, mouse anti-TPI at 1:500, rabbit anti-ATP5B at 1:1000, mouse anti-Tubulin alpha-4A chain at 1:1,000, goat anti-mouse horseradish peroxidase at 1:2,000, goat anti-rabbit horseradish peroxidase-conjugated peroxidase at 1:2,000.

#### Hexokinase 1 Release Assay

To evaluate the release of HK1 from mitochondria as a function of Glucose-6-Phosphate (G6P) concentration, mitochondria were isolated from HEK239T cells (expressing WT or T259A-HK1-GFP together with OGT overexpression and 10 µM Thiamet-G treatment or under control conditions) using modified mitochondria isolation buffer (5 mM HEPES (pH 7.4), 250 mM Sucrose, 5 mM D-Glucose, 1X Protease Inhibitor Cocktail Set III EDTA Free, 0.1 mM PMSF, 2 mM DTT, 10 µM Thiamet-G, 40 mM N-Acetyl-D-glucosamine), as described above. To perform the HK1 release assay, equal amounts of purified mitochondria were resuspended in a release buffer (5 mM MgCl2·6H2O added into modified mitochondria isolation buffer) containing 0, 250 or 500 µM G6P (Figure 6D). The samples were incubated for 30 minutes at room temperature. The mitochondria were pelleted by centrifugation at 10,000xg for 10 minutes at 4°C. The supernatant containing the HK1 fraction released by various concentrations of G6P (flow through (FT)) was collected. The mitochondrial pellet was resuspended in modified mitochondria isolation buffer and centrifuged again. The mitochondria and FT samples were loaded into SDS-PAGE for western blot analysis and evaluated with indicated antibodies.

#### O-GlcNAcylation measurements

HEK293T or HEK293T-EGFP-HK1 cells were plated in a 6-well plate as mentioned above and transfected with the indicated plasmid constructs the next day. Three days after reaching confluency, cells were washed once with ice-cold 1X phosphate-buffered saline (PBS) containing 8 μM Thiamet-G and lysed in 500 µl buffer containing: 2% NP-40 Alternative, 50 mM Tris-HCl (pH 7.5), 150 mM NaCl, 1 mM EDTA, 40 mM N-Acetyl-D-glucosamine, 8 mM Thiamet-G, 2 mM DTT, 0.1 mg/ml PMSF and Protease Inhibitor Cocktail at 1:1000. Lysates were centrifuged 10 mins at 13,000 xg at 4°C, and the supernatants were collected. For immunoprecipitations of HK2-GFP, and HK1-GFP WT or GlcNAc site mutant (T259A), 2 µg anti-GFP incubated for 2 hours at 4°C with 500 µl of whole-cell lysates, then for 1 hour at 4°C with Protein A Sepharose beads. Beads were washed three times with lysis buffer and resuspended with 1x Laemmli buffer. 80%–90% of immunoprecipitants were separated by SDS-PAGE and transferred to nitrocellulose membranes. For O-GlcNAcylation measurements, blots were first incubated with blocking buffer containing 3% bovine serum albumin (BSA) in 1X TBST, then probed with anti-O-GlcNAc antibody (RL2) overnight^83^. To demonstrate O-GlcNAc modification of endogenous HK1 in neurons, we immunoprecipitated O-GlcNAcylated proteins from mitochondrial fractions using 5 µg anti-O-GlcNAc (RL2) antibody and probed with Anti-HK1 antibody (Figure S3A and B). All buffers have to be made/added fresh and all steps mentioned above have to be done in the same day to preserve O-GlcNAcylation. The same blot was re-probed with rabbit anti-GFP used at 1:1,000, rabbit anti-HK1 (Invitrogen) used at 1:500 or rabbit anti-OGT antibody used at 1:1,000 after washing thoroughly and blocking with 5% non-fat milk in 1X PBST (1X PBS with 0.1% Tween20) for 1-2 hrs at room temperature. For quantitative western blot measurements, image exposure times were optimized for the linear range of detection using Azure C600 Biosystem gel documentation system. The images were further analyzed using Fiji gel analyzer function was used to quantify the intensity of each band.

#### Mass Spectrometry Analysis

HK1-GFP immunoprecipitated from HEK293T cells as described above. HEK293T cells were treated either with 5 µM Thiamet-G or vehicle overnight. 80-90% of each sample were separated by pre-cast 7.5% Mini-PROTEAN TGX precast gel. SimpleBlue stained gel band corresponding to HK1-GFP (as well as the control lane) were excised, minced and prepared for mass spectrometry analysis as previously described^84^. Briefly, the gel was cut to 1 mm by 1 mm cubes and destained three times by first washing with 100 µl of 100 mM ammonium bicarbonate for 15 mins, followed by addition of the same volume of acetonitrile (ACN) for 15 mins. The supernatant was removed, and samples were dried in a speedvac. Samples were then reduced by mixing with 200 µl of 100 mM ammonium bicarbonate-10 mM DTT and incubated at 56°C for 30 mins. The liquid was removed and 200 µl of 100 mM ammonium bicarbonate and 55 mM iodoacetamide was added to gel pieces and incubated at room temperature in the dark for 20 mins. After the removal of the supernatant and one wash with 100 mM ammonium bicarbonate for 15 mins, same volume of ACN was added to dehydrate the gel pieces. The solution was then removed, and samples were dried in a speedvac. For digestion, enough solution of ice-cold trypsin (0.01 µg/µl) in 50 mM ammonium bicarbonate was added to cover the gel pieces and set on ice for 30 mins. After complete rehydration, the excess trypsin solution was removed, replaced with fresh 50 mM ammonium bicarbonate, and left overnight at 37°C. The peptides were extracted twice by the addition of 50 µl of 0.2% formic acid and 5% ACN and vortex mixing at room temperature for 30 mins. The supernatant was removed and saved. A total of 50 µl of 50% ACN-0.2% formic acid was added to the sample, which was vortexed again at room temperature for 30 mins. The supernatant was removed and combined with the supernatant from the first extraction^84^. The combined extractions are analyzed directly by liquid chromatography (LC) in combination with tandem mass spectroscopy (MS/MS) using electrospray ionization.

Trypsin/Gluc-digested peptides were analyzed by ultra-high-pressure liquid chromatography (UPLC) coupled with tandem mass spectroscopy (LC-MS/MS) using nano-spray ionization. The nanospray ionization experiments were performed using a Orbitrap fusion Lumos hybrid mass spectrometer (Thermo) interfaced with nano-scale reversed-phase UPLC (Thermo Dionex UltiMate™ 3000 RSLC nano System) using a 25 cm, 75-micron ID glass capillary packed with 1.7-µm C18 (130) BEHTM beads (Waters corporation). Peptides were eluted from the C18 column into the mass spectrometer using a linear gradient (5–80%) of ACN (Acetonitrile) at a flow rate of 375 μl/min for 1 hr. The buffers used to create the ACN gradient were: Buffer A (98% H2O, 2% ACN, 0.1% formic acid) and Buffer B (100% ACN, 0.1% formic acid). Mass spectrometer parameters are as follows; an MS1 survey scan using the orbitrap detector (mass range (m/z): 400-1500 (using quadrupole isolation), 60000 resolution setting, spray voltage of 2400 V, Ion transfer tube temperature of 285°C, AGC target of 400000, and maximum injection time of 50 ms) was followed by data dependent scans (top speed for most intense ions, with charge state set to only include +2-5 ions, and 5 seconds exclusion time, while selecting ions with minimal intensities of 50000 at in which the collision event was carried out in A-high energy collision cell (HCD Collision Energy of 30%), and the fragment masses where analyzed in the ion trap mass analyzer (With ion trap scan rate of turbo, first mass m/z was 100, AGC Target 5000 and maximum injection time of 35ms), followed by B) Electron Transfer Dissociation (ETD Collision Energy of 25%, and EThcD setting active (SA Collision Energy of %25)), and the fragment masses where analyzed in the ion trap mass analyzer (With ion trap scan rate of turbo, first mass m/z was 100, AGC Target 5000 and maximum injection time of 35 ms), Data analysis was carried out using the Byonic™ (Protein Metrics Inc.) or Peaks 8.5 (Bioinformatics solutions).

#### Structural Analysis of Hexokinase 1

X-ray crystallography structure of HK1 (PDB ID: 1CZA) was used for the structural analysis^18^. The secondary structure of the unmodeled portion of HK1 (amino acid residue 1-15) was predicted with Iterative Threading ASSEmbly Refinement (I-TASSER)^85^, and manually appended to the crystal structure using Visual Molecular Dynamics (VMD). The full-length structure was then minimized using stochastic gradient descent (SGD) and equilibrated for 200 ns in explicit water. The equilibrated HK1 structure was then used to construct a variant with the T:259 O-GlcNAc post-translational modification using Charmm-GUI’s standard procedures^86^. The O-GlcNAc modified variant of HK1 was minimized using SGD and equilibrated for 200 ns in water and counter ions at a concentration of 0.15 M. Finally, a conformational search was performed using Replica Exchange Solute Tempering (REST2) in Nanoscale Molecular Dynamics (NAMD)^20,87^. The REST2 simulations were performed with 30 replicas over a linear temperature range from 300 to 500 K. The number of exchange attempts was 5000 with 20000 integration steps between each exchange for 200 ns for each of the 30 replicas totaling 6 µs of sampling.

To determine how T259 O-GlcNAcylation alters HK1 function, Poisson-Boltzmann calculations were performed using the APBS software suite^88^ with the following parameters. First, the linearized Poisson-Boltzmann equation was solved with the single Debye-Huckel boundary condition and the *smol* option for the molecular surface’s representation. The *cglen* and *fglen* options were chosen to ensure a 1 Å resolution electrostatic potential grid. Temperature and ion concentrations were set with 300 K and 0.15 M. The protein and solvent dielectric constants were set to 12.0 and 78, respectively. The radius of the solvent was set to that of water, 1.4 Å. Finally, the atomic radii and charges were set as defined in the CHARMM36 force field^89^.

#### Glucose-6-Phosphate Measurements

To measure glucose-6-phosphate (G6P) levels, HEK293T cells were transfected with indicated constructs and detached from 6-well-plate with cell scraper. Cell lysates were prepared in 500 µl ice-cold 1X PBS using 25G 5/8 and 30G 1/2 syringe needle. Lysates were centrifuged 5 minutes at 13,000 xg in 4°C and the supernatants were collected. The supernatant was filtered through Amicon® Ultra-0.5 ml Centrifugal Filters, Ultracel-10K by centrifuged 15 mins at 14,000 xg at 4°C. The concentrated samples were used to measure total protein concentrations via BCA Protein Assay and the deproteinized samples were used to measure G6P levels by following the protocol provided in the G6P Assay Kit. Colorimetric measurements were made at room temperature using Spark 20M. Expression level of wild-type (WT) or GlcNAc site mutant (T259A) HK1-GFP were analyzed by western blotting from cell lysates.

#### Respirometry Measurements with HEK293T

Oxygen consumption (OCR) and extracellular acidification rates (ECAR) were measured using an Agilent Seahorse XFe96 Analyzer with Seahorse XF Mito Stress Test and Seahorse XF Real-Time ATP Rate protocol from HEK293T cells. HEK239T cells were initially cultured on 6-well-plate as described above, after 16-18 hrs of transfection, cells were re-plated on XF96e plates at 2.5x10^3^/mm^2^ density. After 2-3 hrs of replating, cells were treated with 10 µM OGA inhibitor Thiamet-G or vehicle control (DMSO) overnight. For respirometry measurements, the DMEM was exchanged with XF DMEM Base Medium (pH 7.4) with no phenol red supplemented with 5 mM glucose and 1 mM pyruvate. NucBlue live cell stain was included to stain nuclei for cell counting immediately after the assay. Respiration was measured under basal conditions as well as after injections of 2 µM (for ATP production rate assay) /0.5 µM (for mitochondria stress test) Oligomycin, 0.5 µM FCCP, and 0.5 µM Rotenone/Antimycin A. To recruit HK1 to mitochondria surface using chemically induced dimerization strategy^79^ (Figure 5D-F), AP21967 was applied at final concentration of 500 nM to HEK239T for 15-20 minutes prior to respirometry measurements. ATP production rates was calculated according to protocol provided by Agilent. Basal and maximal respiration (OCR) as well glycolytic capacity (ECAR) values were plotted after cell count normalization with FluxNorm Normalization with cell count was done by FluxNorm ^40^.

#### Verification of Hexokinase 1 knock-down with shRNA and rescue

For the HK1 knockdown experiments, 14 shRNA constructs from the Sigma MISSION shRNA library were screened. The most effective shRNA construct (TRCN0000297076) was selected and subsequently used to examine the HK1 knockdown efficiency. Because the target sequence was conserved between mouse and rat HK1, the knockdown efficiency was first validated in Neuro-2a cells. The specificity of the HK1 shRNA construct was verified by co-expressing shRNA resistant human WT or T259A HK1-GFP for 3 days in and then measuring the level of GFP signal from cell lysates (Figure S4B and C). After confirming the mouse HK1 knockdown efficiency, HK1 shRNA construct was further characterized in rat primary neuron cultures. Rat hippocampal neurons were transfected with either TRC2 pLKO.5-puro empty vector control plasmid DNA or HK1 shRNA construct together and Mito-DsRed. Given the half-life of HK1 protein was estimated to be ∼72 hours^90^, shRNA construct was expressed in rat neurons for 3 days, allowing efficient knockdown of endogenous HK1 to be achieved. The efficiency of rat HK1 knockdown was verified retrospectively for each transfection by immunocytochemistry using an antibody against endogenous HK1 (Figure S4D). shRNA-resistant human GFP or BFP-tagged WT-HK1 and T259A O-GlcNAc silent base mutation constructs were used together with shRNA construct to maintain endogenous HK1 levels for all experiments (Figures S4D-F).

#### Live HEK293T cells imaging and analysis

HEK293T cells were transfected with a plasmid expressing Tom20-mCherry-FKBP^79^ and FRB-Tr-HK1-GFP. After 24 hours, cells were imaged using a Zeiss LSM 780 confocal laser scanning microscope equipped with a C-Apochromat 40x/1.20 W Korr FCS M27 objective with excitation at 488 nm and 561 nm separately. To investigate the time course of rapalog (AP21967)-induced dimerization, cells were treated with 500 nM AP21967 or DMSO as a vehicle control, and live-cell imaging was performed for 20 minutes. It was found that 10-15 minutes of AP21967 treatment was sufficient to induce HK1 recruitment to mitochondria. HEK293T cells were continuously perfused at 0.2-0.25 ml/min with Hibernate E low fluorescent, 5 mM glucose during image acquisition at 37°C. Time-lapse images were acquired every 5 seconds at <1.5% laser power for each channel to minimize phototoxicity.

#### Live neuron imaging and analysis

*vGLUT1-pH and GCaMP live imaging with electrical stimulation:* Primary hippocampal neurons were transfected with indicated DNA constructs using Lipofectamine 2000. Live-cell imaging of neurons were performed 2-3 days after transfections at 11-14 DIV. Coverslips were mounted on a stimulation chamber with laminar flow for perfusion and imaged at 37°C using Zeiss LSM 780 laser scanning confocal microscope equipped with heated stage and C-Apochromat 40x/1.20 W Korr FCS M27 objective with highly sensitive low dark noise PMTs (2x) and GaAsP (32x) array. Laser power was set to <1% for each channel to minimize phototoxicity during time-lapse image acquisition. For all experiments, neurons were continuously perfused at 0.2-0.25 ml/minute with modified Tyrode buffer (50 mM HEPES (pH 7.4), 119 mM NaCl, 2.5 mM KCl, 2 mM CaCl2, 2 mM MgCl2, 5 mM D-glucose, 2 mM pyruvate, and 2mM lactate, supplemented with 0.01 mM CNQX, 0.05 mM APV to suppress postsynaptic responses), pH measured at 37°C. Trains of action potentials were evoked by current pulses of 100 mA, at 10 Hz for 10 seconds with 5 minutes of recovery between runs for live-cell imaging to measure synaptic vesicle recycling rates with vGLUT1-pH (pHluorin-tagged vesicular glutamate transporter 1) ^91^, and calcium dynamics with GCaMP6s. For these measurements, endogenous HK1 was knocked down with shRNA, and rescued with BFP tagged WT and T259A-HK1 to avoid overexpression. The stimulus pattern was generated with an Arduino connected to an isolated bipolar stimulator.

For vGLUT1-pH imaging, time-lapse movies were acquired from 11-14 DIV neurons for a total of 120 seconds at 10 Hz. For each imaging session, the first pre-stimulus fluorescence baseline (F0) measurements were performed for 20 seconds in the absence of electrical activity then action potentials were evoked by electrical field stimulation (100 mA,10 Hz,100 APs). At the end of experiments, neurons were perfused with Tyrode buffer containing 50 mM NH4Cl at pH 7.4, which rapidly equilibrates pH and increase the vGLUT1-pH signal, and imaging continued with 1s interval to obtain Fmax value for each imaged axonal segment. When indicated, 1 µM Tetrodotoxin (TTX) were added to neuron maintenance media right after transfection with WT/T259A-HK1-BFP constructs to preserve ATP use due to spontaneous neuronal activity. Two hours before live imaging, TTX was removed and washed off twice using culture media.

For calcium imaging, time-lapse movies were acquired from 11-12DIV neurons expressing GCaMP6s^80^ and indicated HK1 constructs for a total of 30 seconds at 10 Hz. For each imaging session, first pre-stimulus fluorescence baseline (F0) measurements were performed for 5 seconds in the absence of electrical activity then action potentials were evoked by electrical field stimulation (100mA,10 Hz,100 APs). At the end of each experiment, neurons were perfused with Tyrode buffer containing 50mM KCl to achieve maximal response (Fmax) for each imaged axonal segment.

Images were analyzed using Fiji plugin Time Series Analyzer (v3.0). ∼2 μm regions of interests from 8-11 neurons corresponding to presynaptic boutons were selected for image analysis. ΔF values for vGLUT1-pH and GCaMP6s images were calculated as previously described after background substructions ^75^. F0 values were defined by averaging data points from pre-stimulus period, while Fmax values were determined by averaging data points following NH4Cl applications for vGLUT1-pH and KCl application for GCaMP6s images. Endocytic time constants (τ) were calculated by fitting the fluorescent change after the electrical stimulus to a single exponential decay ^91^.

*Glucose level-dependent cytoplasmic and mitochondrial HK1 imaging:* For live cell imaging, endogenous HK1 was knocked down with shRNA, and rescued with GFP tagged WT and T259A-HK1 to avoid over expression. Primary hippocampal neurons were transfected with indicated DNA constructs using Lipofectamine 2000. Live-cell imaging of neurons were performed 2-3 days after transfections at 11-14 DIV. Yokogawa W1 SoRa scan head on the Nikon Ti-2E microscope, equipped with NIS Elements software and Photometrics Prime 95B camera, was used in SoRa mode with a 2.8x zoom lens. The fluorophores were excited using the 488 nm and 561 nm lines of a six-line (405 nm, 445 nm, 488 nm, 515 nm, 561 nm, and 640 nm) LUN-F-XL laser engine. For simultaneous acquisition, a quad bandpass filter (Chroma ZET405/488/561/640mv2) was placed in the emission path of the W1 scan head, and the emission was split with a Cairn TwinCam with a 580LP filter. The GFP emission was reflected and passed through a 514/30 BP filter onto camera 2. The mCherry fluorescence was passed through a 617/73 and an additional 600/50 filter to camera 1. Camera alignment was carried out by parking the W1 disk and projecting transmitted light through the pinholes of the disk onto both cameras. Camera 2’s image of the pinholes was shifted relative to camera 1 until the patterns matched at 1200% zoom. An Okolab Bold Line stage-top incubator that was designed to fit in the piezo Z-stage (MadCity Labs) was used to maintain 37°C and 5% CO2. For all the experiments, neurons were continuously perfused with 0mM (also containing 1mM lactate and pyruvate), 1mM (also containing 1mM lactate and pyruvate) or 5mM glucose containing Hibernate E low fluorescent imaging media as indicated. Distal axons were selected for live-cell imaging. Images were analyzed using Fiji intensity plot function.

#### Statistical Analysis

Throughout the paper, data are expressed as mean ± SEM unless otherwise noted. All p-values and number of replicas were indicated in the figure legend for each experiment. Statistical analysis was performed with GraphPad Prism v7.0 for MacOS. The Mann-Whitney U test was used to determine the significance of differences between two unpaired conditions. Multiple conditions were compared by Kruskal-Wallis nonparametric ANOVA test, which was followed by Dunn’s multiple comparisons test or by one-way ANOVA with post hoc Tukey’s test as appropriate to determine significance of differences across every condition to control condition.

### RESOURCES TABLE

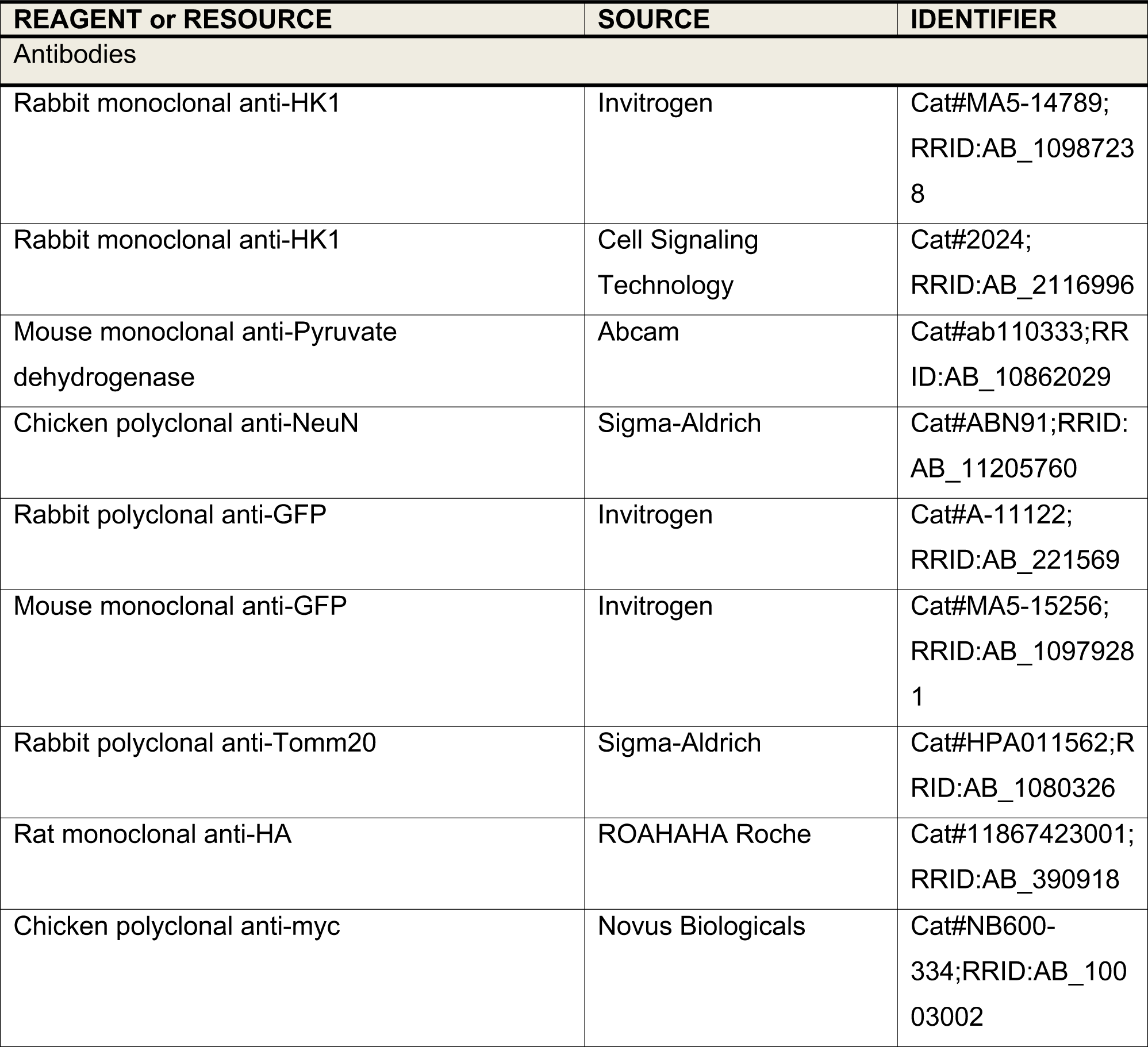
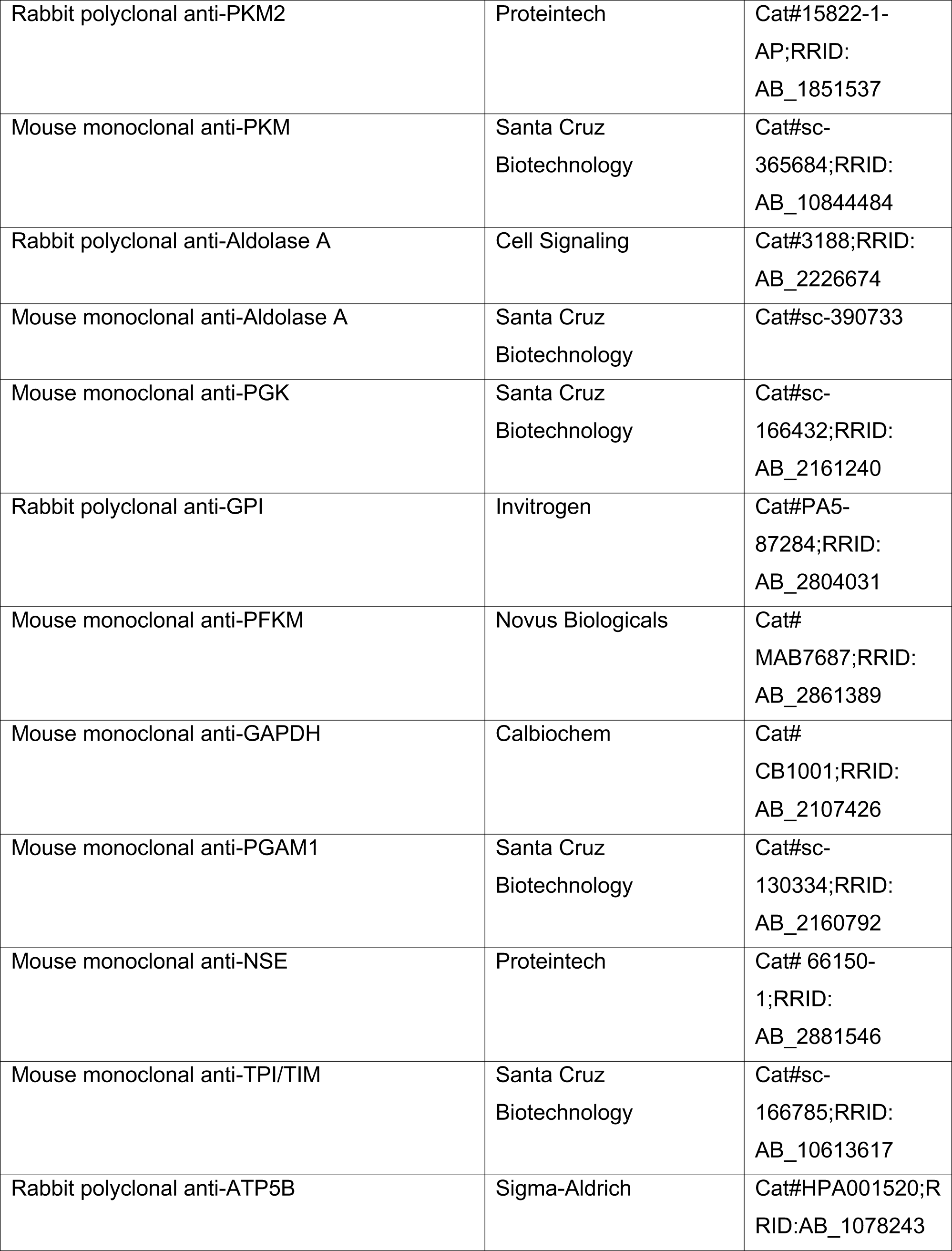
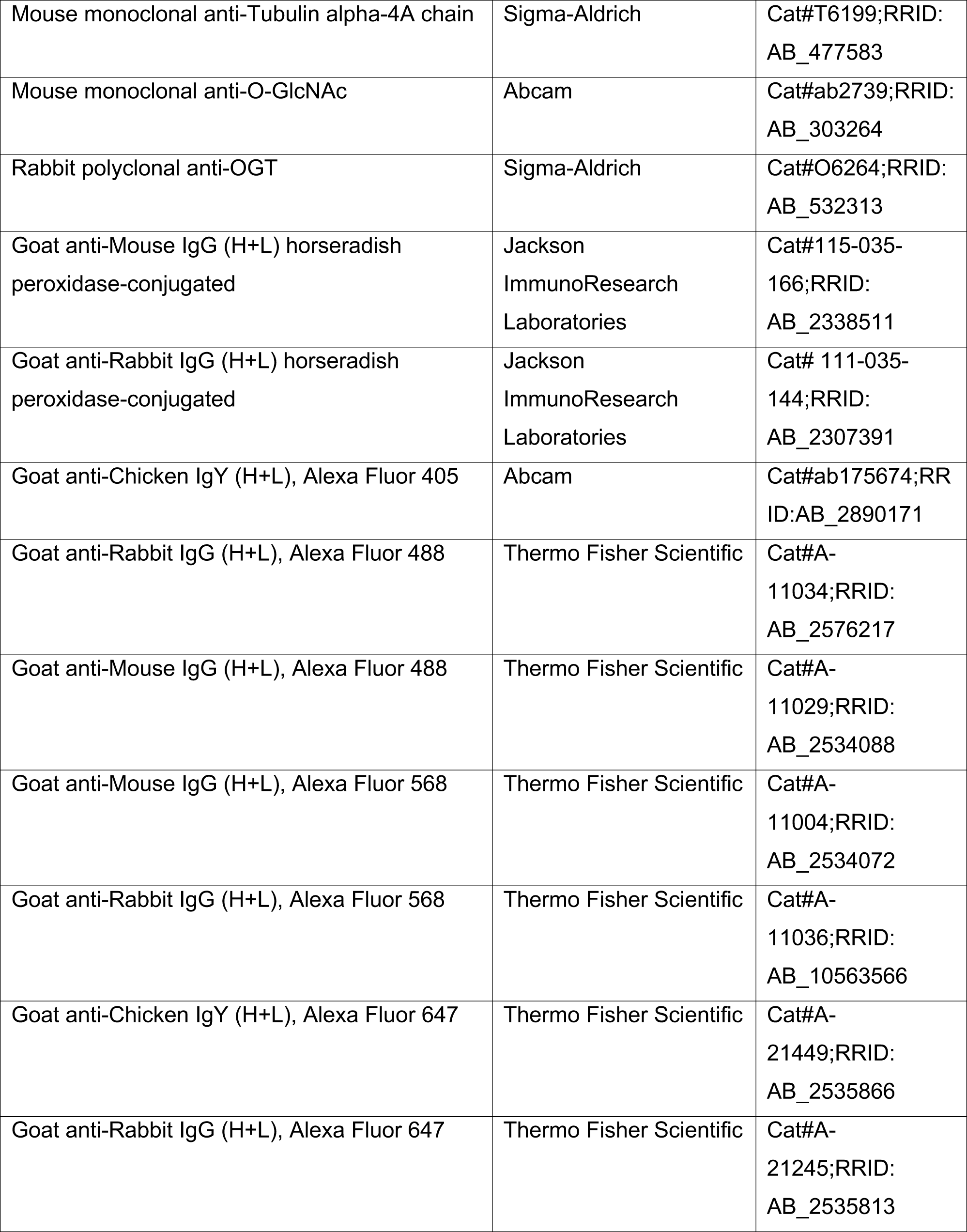
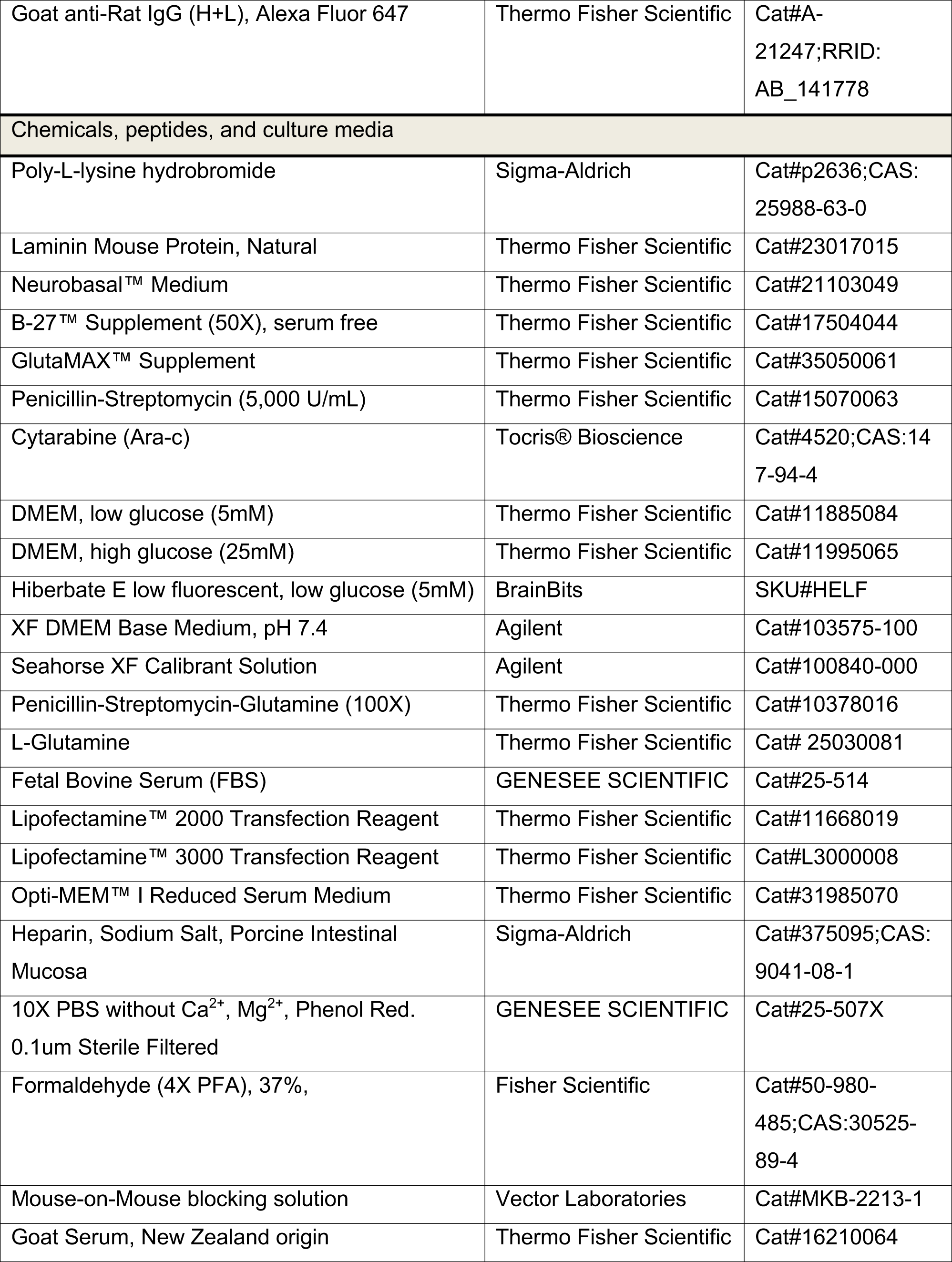
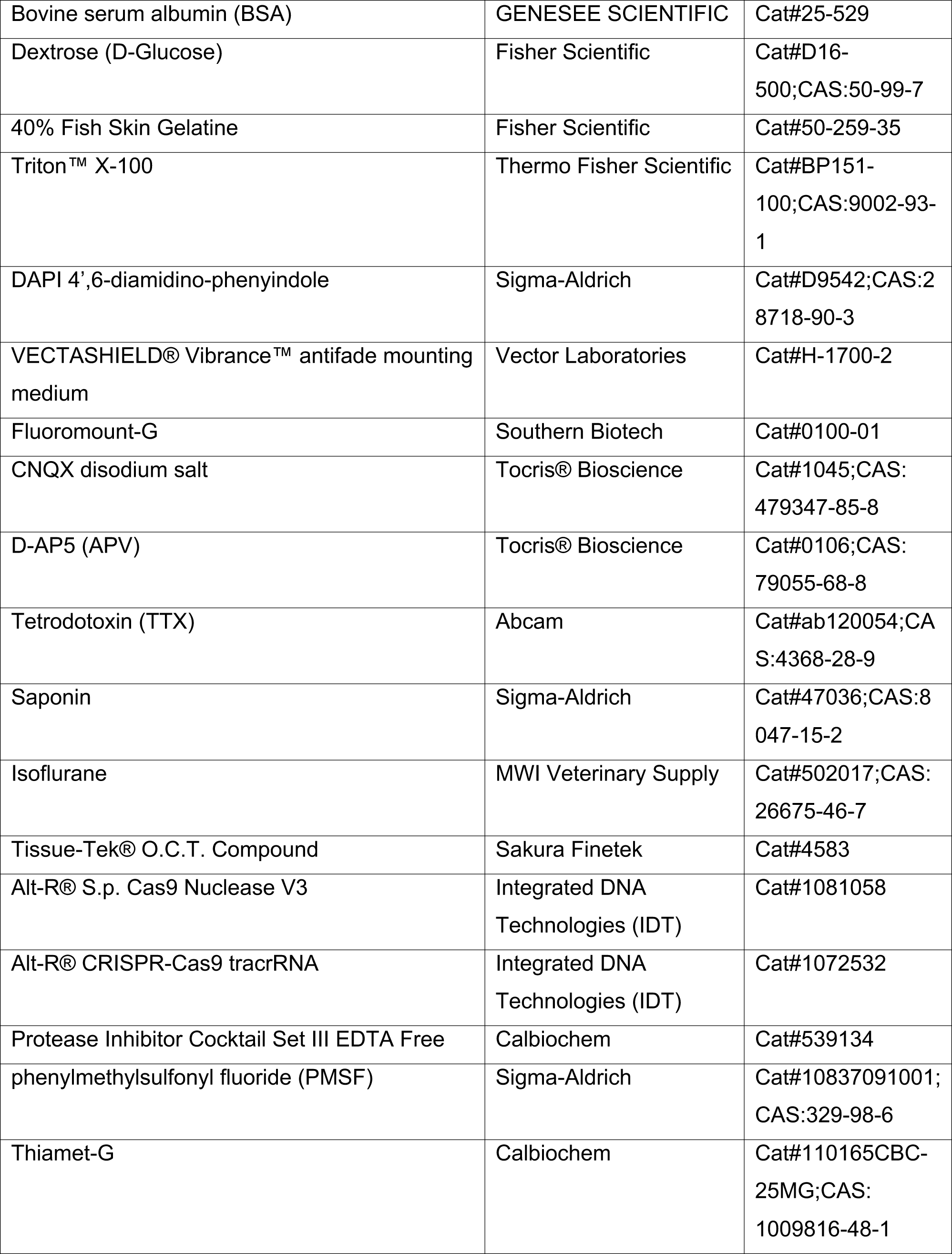
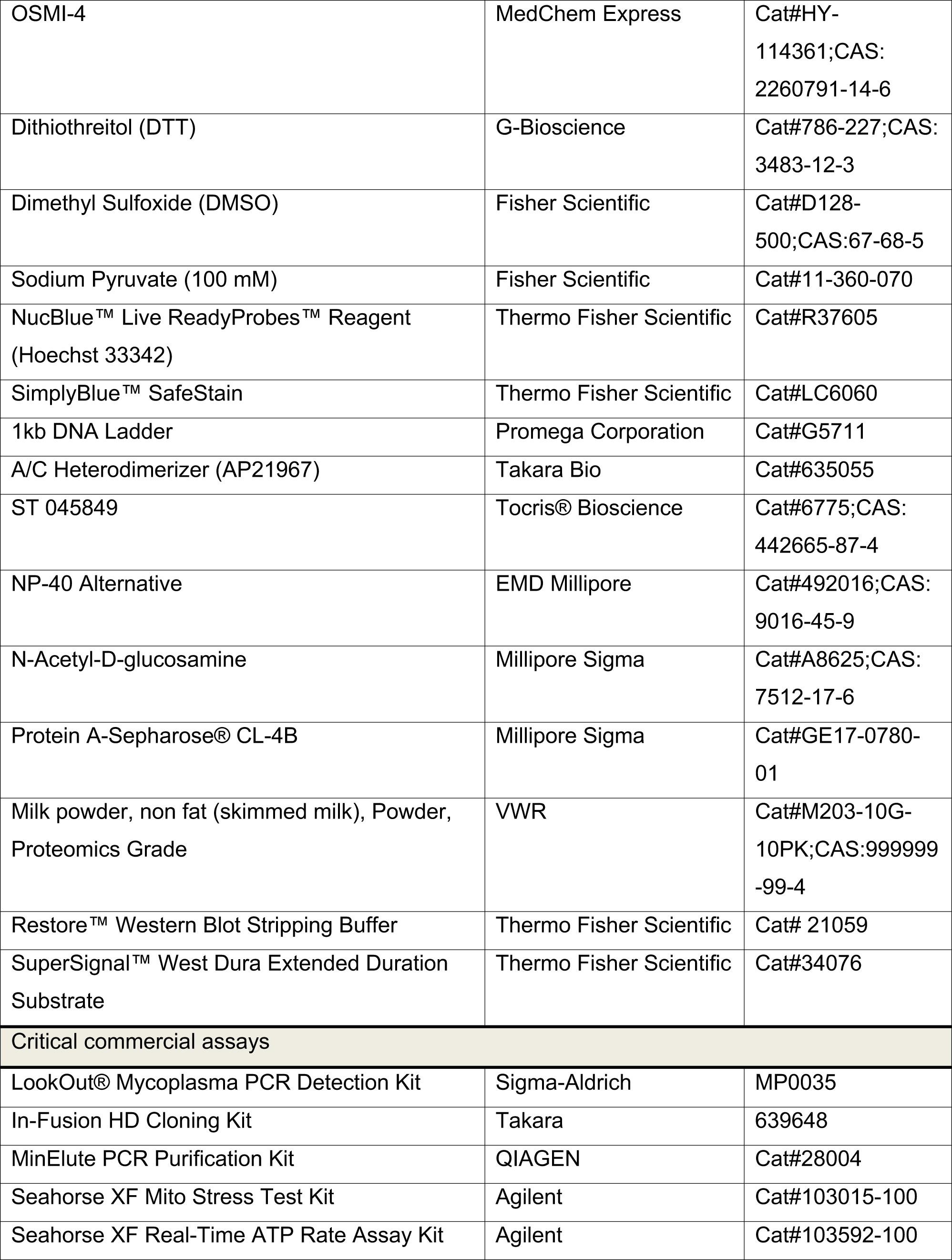
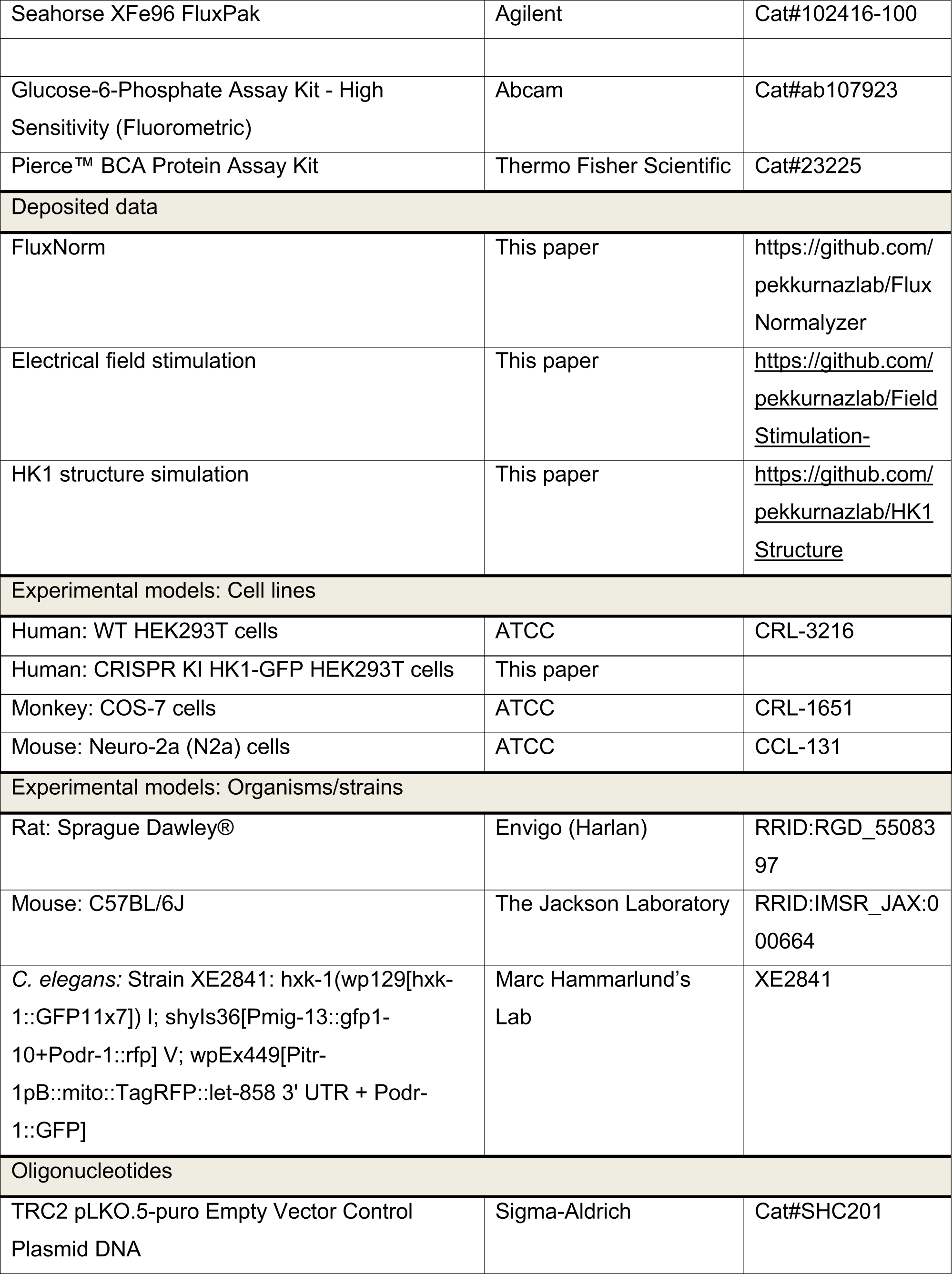
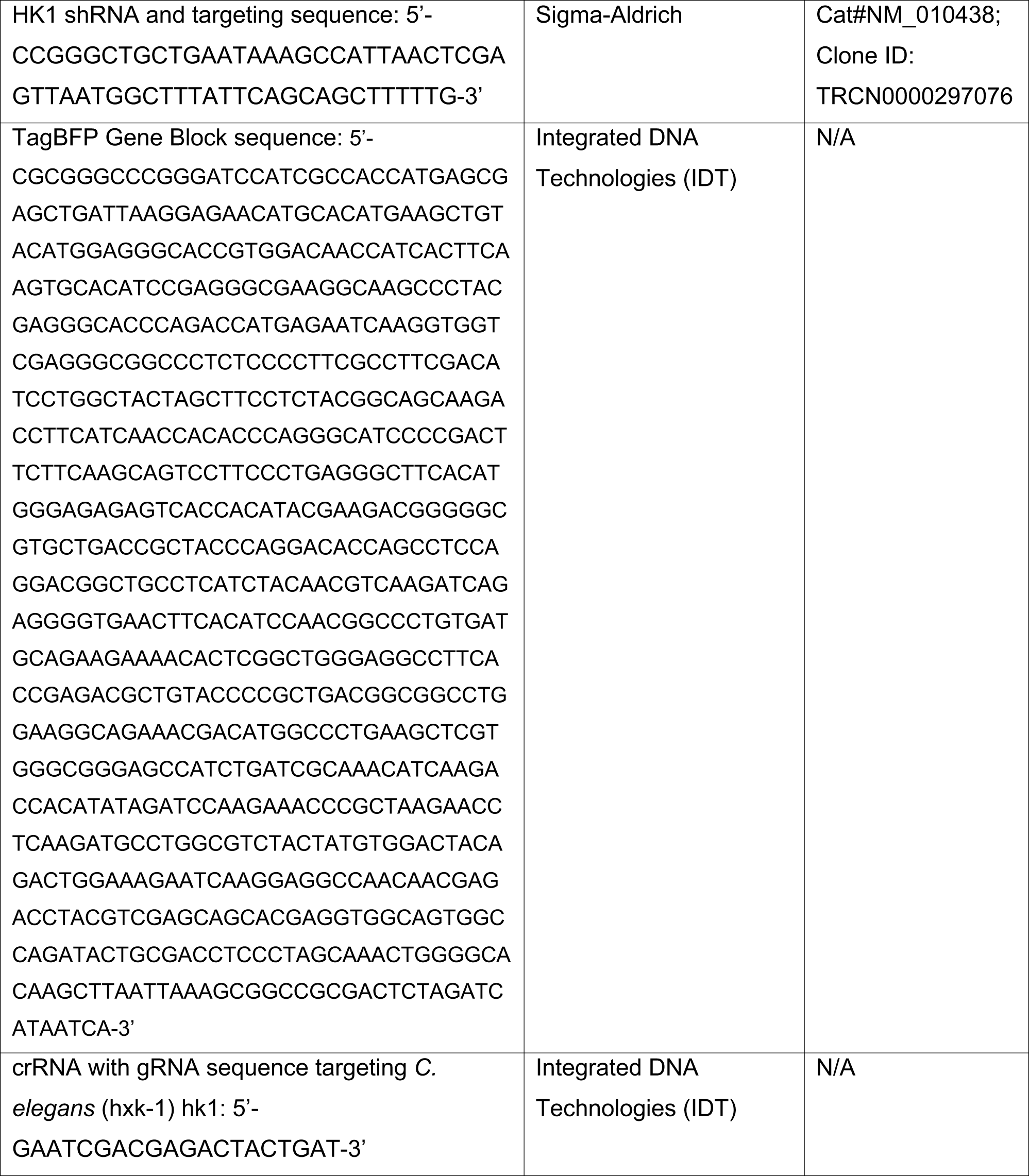
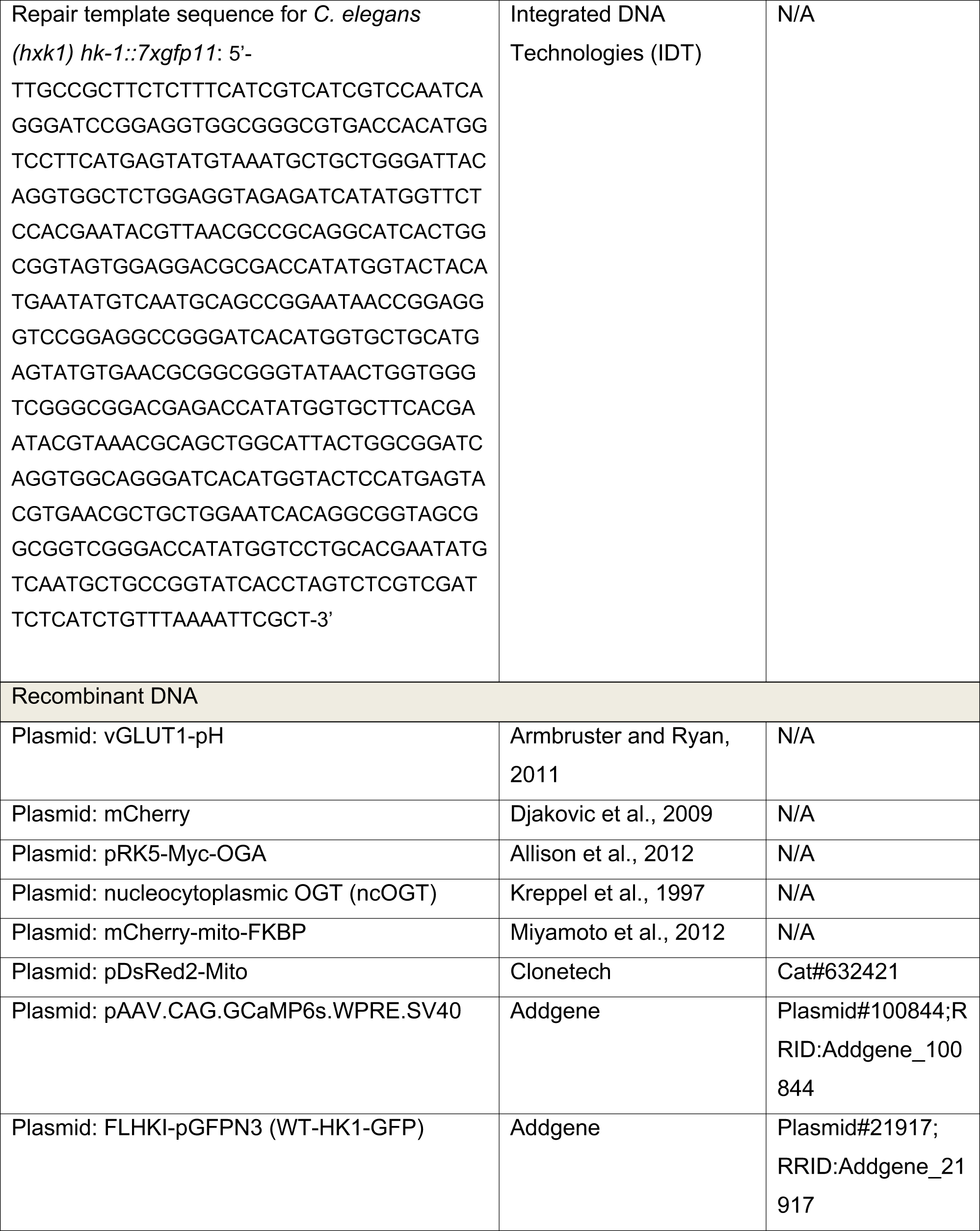
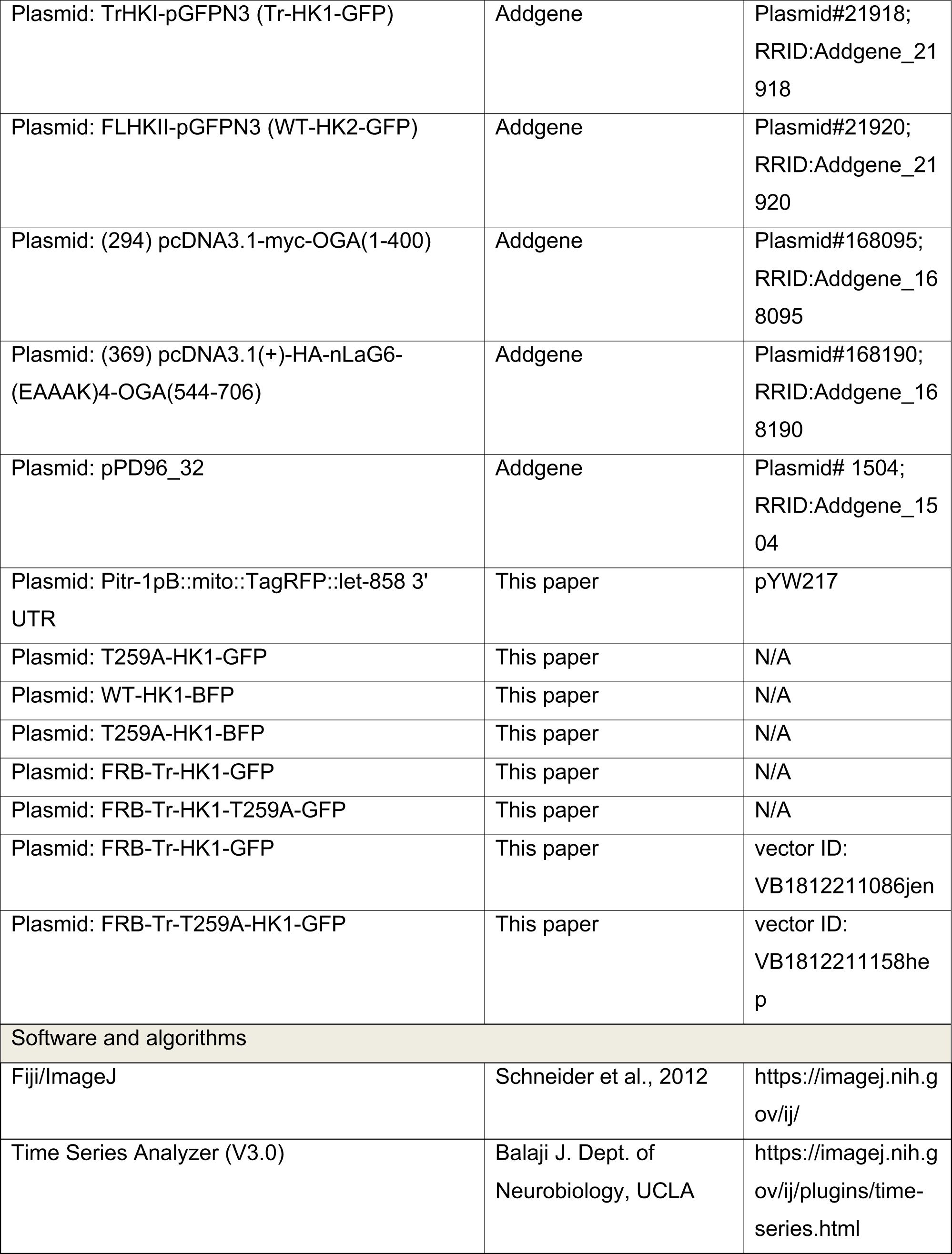
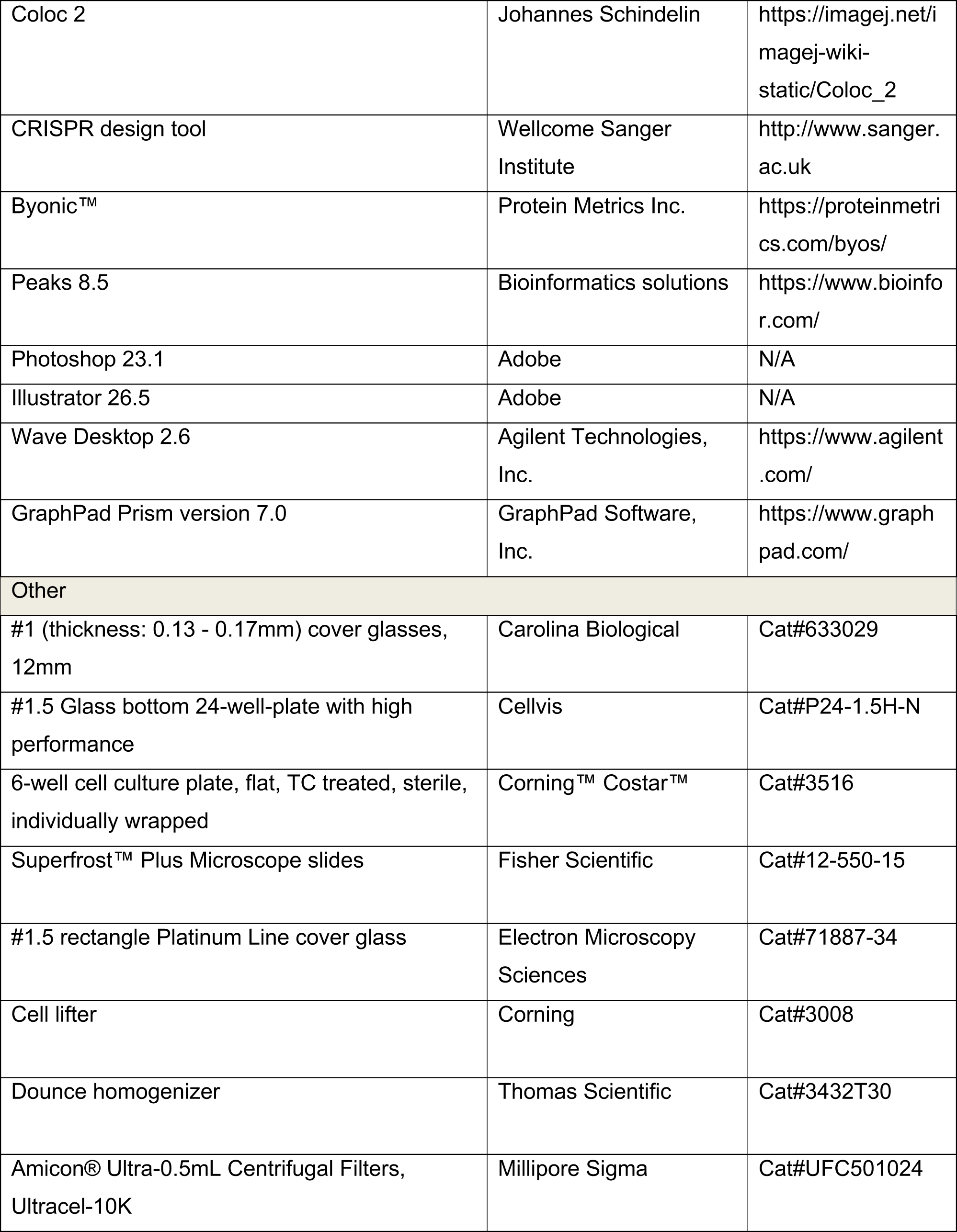
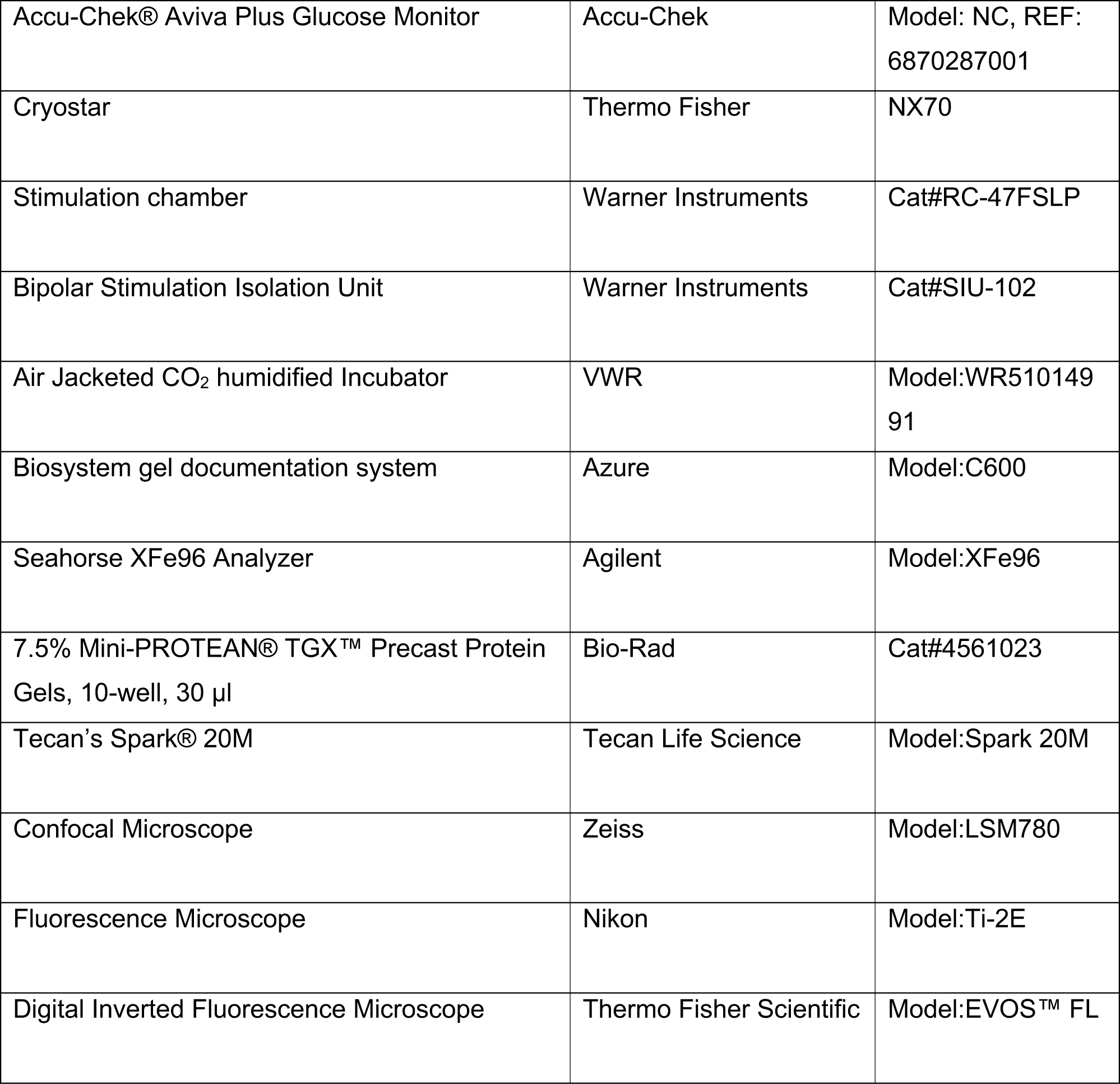

### RESOURCE AVAILABILITY

#### Lead contact

Further information and requests for resources should be directed and will be fulfilled by the lead contact, Gulcin Pekkurnaz.

#### Materials availability

All biological resources and tools are either available from commercial sources or lead contact.

#### Data and code availability

Agilent Seahorse XF96e Metabolic Flux Analyzer data normalization is performed via the custom-written macro “FluxNorm”, available at https://github.com/pekkurnazlab/FluxNormalyzer. Electrical field stimulation code is available at https://github.com/pekkurnazlab/FieldStimulation-. HK1 structure simulation data are available at https://github.com/pekkurnazlab/HK1Structure. Any additional information required to reanalyze the data reported in this paper is available from the lead contact upon request.

---

**Figure S1.**
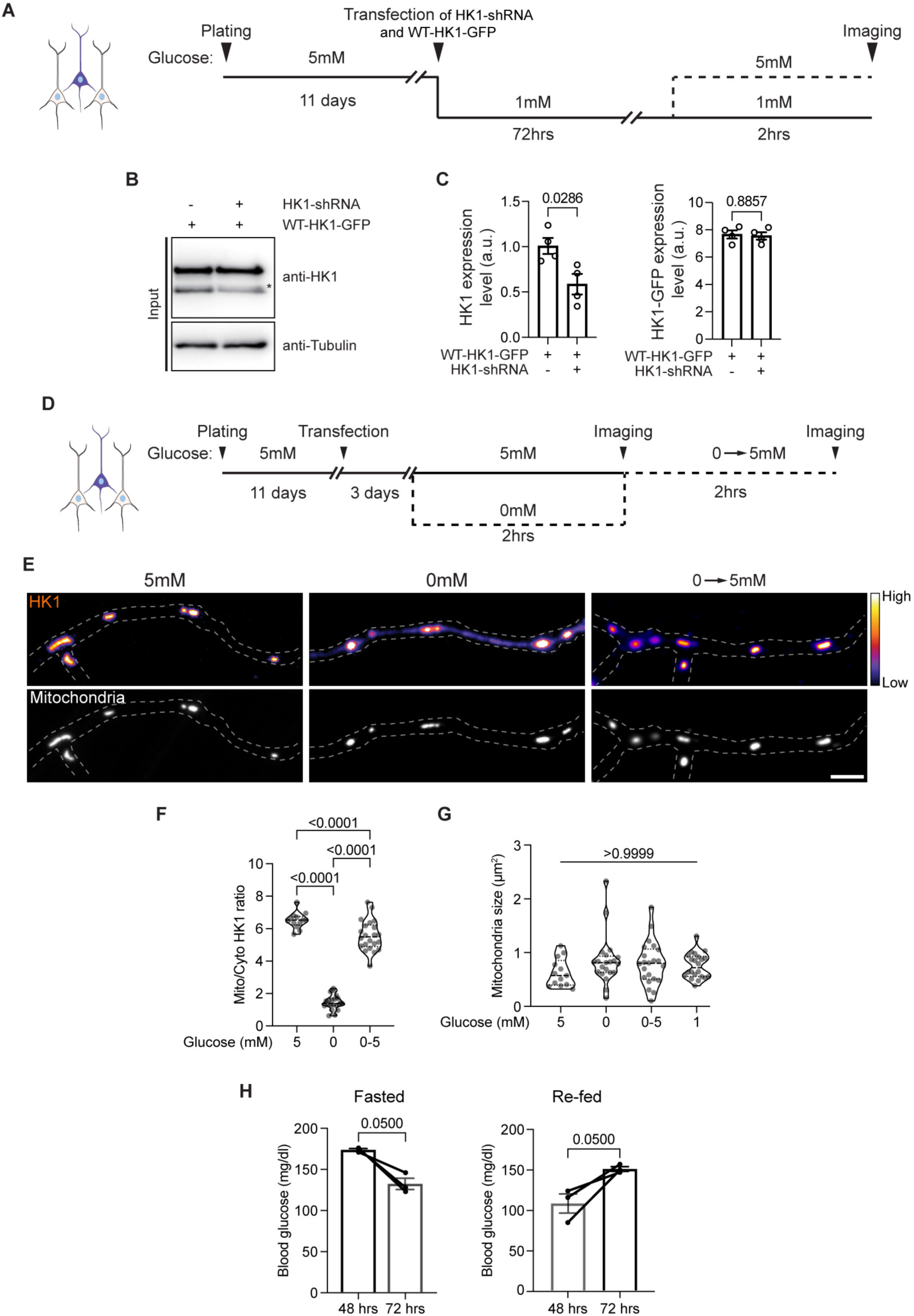
Glucose level modulates Hexokinase 1 localization. (A) Experimental scheme detailing the sequence of plating, transfection, imaging, and alteration of extracellular glucose levels in cultured rat hippocampal neurons. (B) Evaluation of HK1-shRNA knock-down efficiency in Neuro-2a cells. HK1-shRNA and shRNA resistant eGFP tagged HK1 (HK1-GFP) were expressed for 48-72 hrs, and whole cell lysate (Input) were probed with anti-GFP and anti-Tubulin (loading control) antibodies. Asterisk indicates endogenous HK1. (C) Quantification of endogenous HK1 (left) and eGFP-tagged HK1 (WT-HK1-GFP) (right) expression levels as shown in (B). All values are shown as mean ± SEM. n= 4 (Mann-Whitney U test). (D) Experimental scheme detailing the sequence of plating, transfection, and imaging conditions for the 0mM glucose experiments in cultured rat hippocampal neurons. (E) Axonal localization of HK1 in cultured rat hippocampal neurons transfected with HK1-shRNA, shRNA-resistant eGFP-tagged HK1 (pseudo-color, fire), and Mito-DsRed (gray). Representative axonal images were captured at 5mM glucose, following a 2-hour exposure to 0mM glucose, and at 5mM glucose after 2 hours exposure to 0mM glucose (1mM lactate and pyruvate), as depicted in (C). Scale bar represents 5µm. (F) The mitochondrial (Mito) and cytoplasmic (Cyto) HK1 intensity ratios were quantified along axons. Data are presented as a violin plot with individual data points and associated p-value. (G) Mitochondrial size measurements along the axons under varying extracellular glucose levels. n = 94-117 mitochondria, 9-10 axons from three independent experiments (one-way ANOVA with post hoc Tukey’s multiple comparison test). (H) Blood glucose measurements from the Fasted and Re-fed mice used for comparing subcellular localization of Hexokinase 1 as shown in Figure 1. n = 3 mice for each condition, three independent experiments. All values are shown as mean ± SEM (Mann-Whitney *U* test).

**Figure S2.**
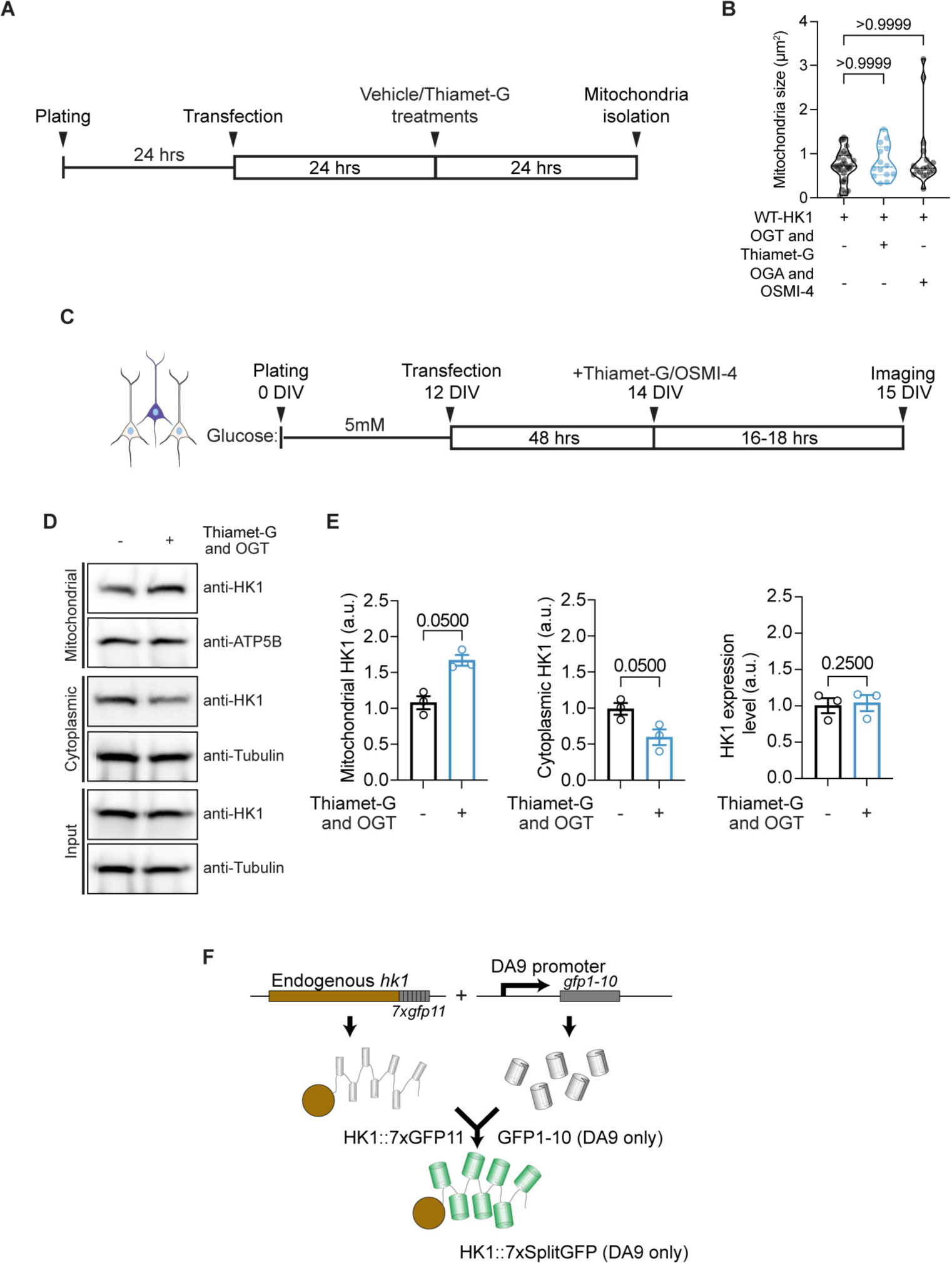
O-GlcNAcylation promotes mitochondrial enrichment of Hexokinase 1 in various cell types. (A) The experimental timeline illustrating the sequence of HEK293T cell plating, OGT transfection, administration of vehicle or Thiamet-G, and mitochondrial isolation. (B) Analysis of mitochondrial size in cultured rat hippocampal neurons co-transfected with HK1-shRNA and eGFP-tagged HK1 to achieve endogenous HK1 levels. O-GlcNAcylation level was upregulated by ectopic OGT expression and Thiamet-G treatment, and downregulated by OGA expression and OSMI-4 treatment. Data are presented as a violin plot with individual data points and associated p-values. n= 83-120 mitochondria, 11-13 neurons, three independent experiments (one-way ANOVA with post hoc Kruskal-Wallis multiple comparison test). (C) Experimental timeline outlining the sequence of plating, transfection, Thiamet-G and OSMI-4 treatments, and imaging of cultured rat hippocampal neurons in 5mM glucose for experiments illustrated in Figure 2D. (D) Western blot analysis of whole cell lysate (Input), isolated mitochondrial and cytoplasmic fractions from HEK293T. The samples were probed with antibodies against HK1, ATP5β (mitochondrial marker), and Tubulin (cytoplasmic marker) with or without ectopic OGT expression and Thiamet-G or vehicle treatments. (E) Quantification of HK1 levels in mitochondrial (left), cytoplasmic (middle), and whole cell lysate (right) under indicated different conditions. n = 3 (all values are shown as mean ± SEM, Mann-Whitney *U* test). (F) A schematic illustration of Native and Tissue-specific Fluorescence (NATF) method used for endogenous labeling of HK1 in *C. elegans* DA9 neuron.

**Figure S3.**
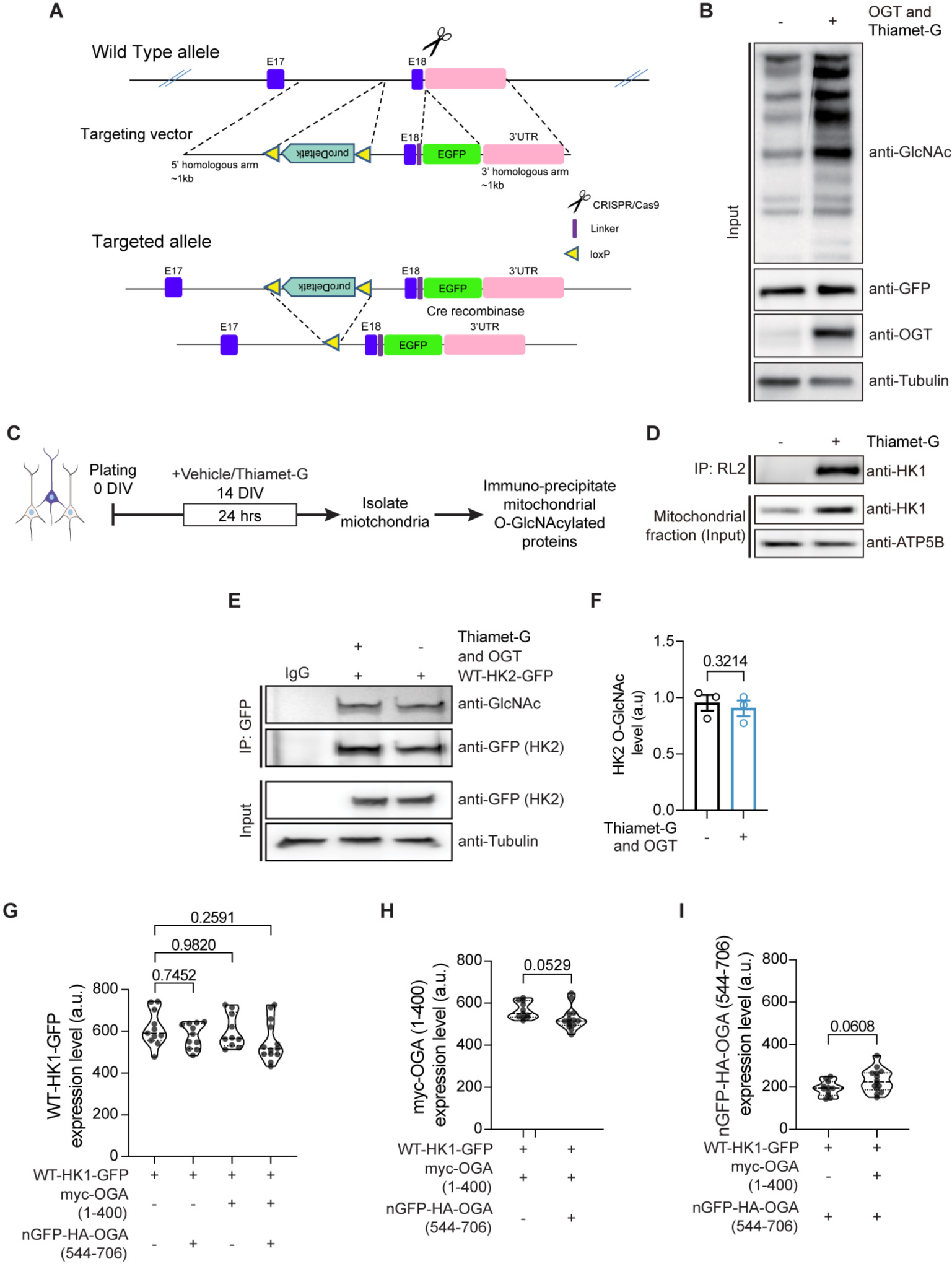
Elucidating the O-GlcNAc modification of Hexokinase 1 and 2. (A) Illustration of CRISPR-based approach to add eGFP tag at the C-terminal of HK1 in HEK293T cells. The strategy is based on transcript-202 (NM_000188.2) and was implemented by BioCytogen. (B) Western blot analysis of the whole cell lysate (input), used for the generation of mitochondrial and cytoplasmic fractions as shown in Figure 3, from HEK293T cells. The whole cell lysate (Input) from CRISPR edited HEK293T cells was probed with antibodies against O-GlcNAc (RL2), GFP (HK1), OGT and tubulin (loading control), with or without OGT overexpression and Thiamet-G treatments. (C) Schematic demonstrating the sequence of mitochondrial isolation and O-GlcNAc immunoprecipitation (IP) using the anti-O-GlcNAc antibody RL2 from cultured rat hippocampal neurons. (D) Western blot analysis of mitochondrial fraction (Input) and O-GlcNAc IP using antibody against HK1 and ATP5B (mitochondrial marker). (E) eGFP tagged Hexokinase 2 (HK2) was expressed in HEK293T cells. GFP antibody was used to immunoprecipitate (IP) HK2, with or without OGT overexpression and Thiamet-G treatments. The IPs were probed with anti-GlcNAc (RL2) and anti-GFP antibodies. Whole cell lysates (Input) were probed with anti-GFP and anti-tubulin antibodies. Rabbit IgG serves as an IP control. (F) Quantification of HK2 O-GlcNAcylation levels. All values are shown as mean ± SEM, unpaired *t*-test. n= 3. (G-I) Quantification of the expression levels of HK1-GFP (G), myc-OGA (1-400) (H) and nGFP-HA-OGA (544-706) (I) in COS-7 cells, as shown in Figure 3D. Data are presented as a violin plot with individual data points and associated p-value. n= 42 cells, three independent experiments (Unpaired *t*-test, and one-way ANOVA with post hoc Tukey’s multiple comparison test).

**Figure S4.**
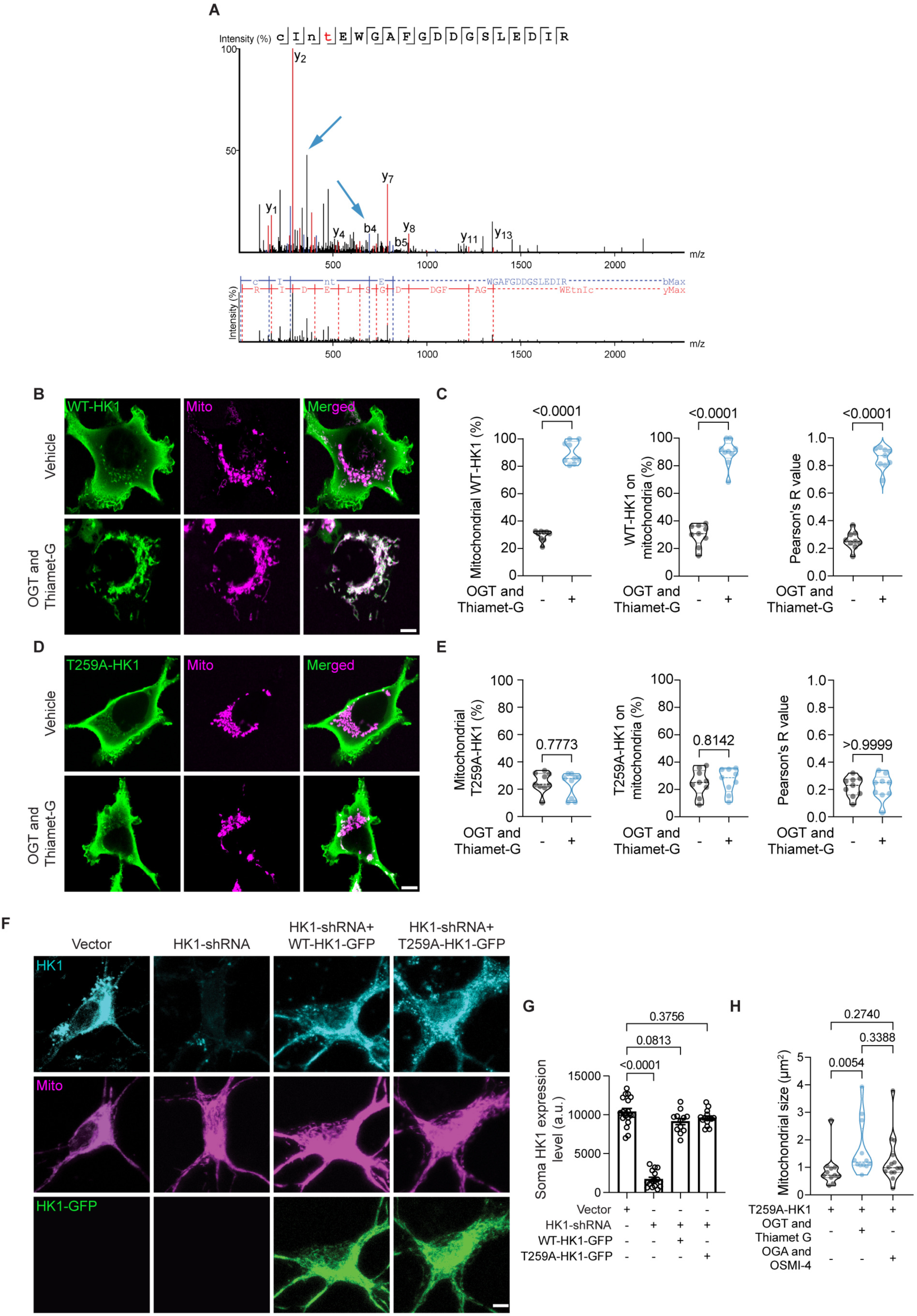
Identification and functional analysis of Hexokinase 1 T259 O-GlcNAcylation Site. (A) Tandem mass spectra showing O-GlcNAc on peptides derived from human Hexokinase 1. Data were acquired using HCD fragmentation and prominent y and b-type ions are labeled. Blue arrow indicates the O-GlcNAc modified threonine (T). Bottom figure demonstrating the survey scan and prominent y/b-type ions. (B-E) Quantitative analysis of HK1 co-localization to measure the percentage of mitochondrial HK1 intensity in HEK293T cells, cultured in 5mM glucose containing media. (B and D) Representative images of HEK293T cells expressing WT-HK1-GFP or T259A-HK1-GFP (green) and Mito-DsRed (magenta) with or without OGT overexpression and Thiamet-G treatments. (C and E) WT and T259A HK1 intensity on mitochondria, percentage of total WT and T259A HK1 on mitochondria and the Pearson’s correlation coefficient (R value) for each condition. Data are presented as violin plots with individual data points and associated p-values. n = 9 cells, three independent experiments (one-way ANOVA with post hoc Tukey’s multiple comparison test). (F) Representative images of hippocampal neurons expressing HK1-shRNA, WT-HK1-GFP, and the O-GlcNAc mutant T259A HK1-GFP (T259-HK1-GFP) are shown in green, along with Mito-DsRed in magenta. These images were stained with an HK1 antibody (in cyan) to visualize the total HK1 expression. (G) Quantification of HK1 expression levels was performed retrospectively for all experiments by analyzing the anti-HK1 staining to ensure consistent endogenous HK1 levels throughout all experiments. (H) Quantification of the size of the mitochondria along the axons as depicted in Figure 4G. n= 81-86 mitochondria from 10-13 axons from three independent experiments.

**Figure S5.**
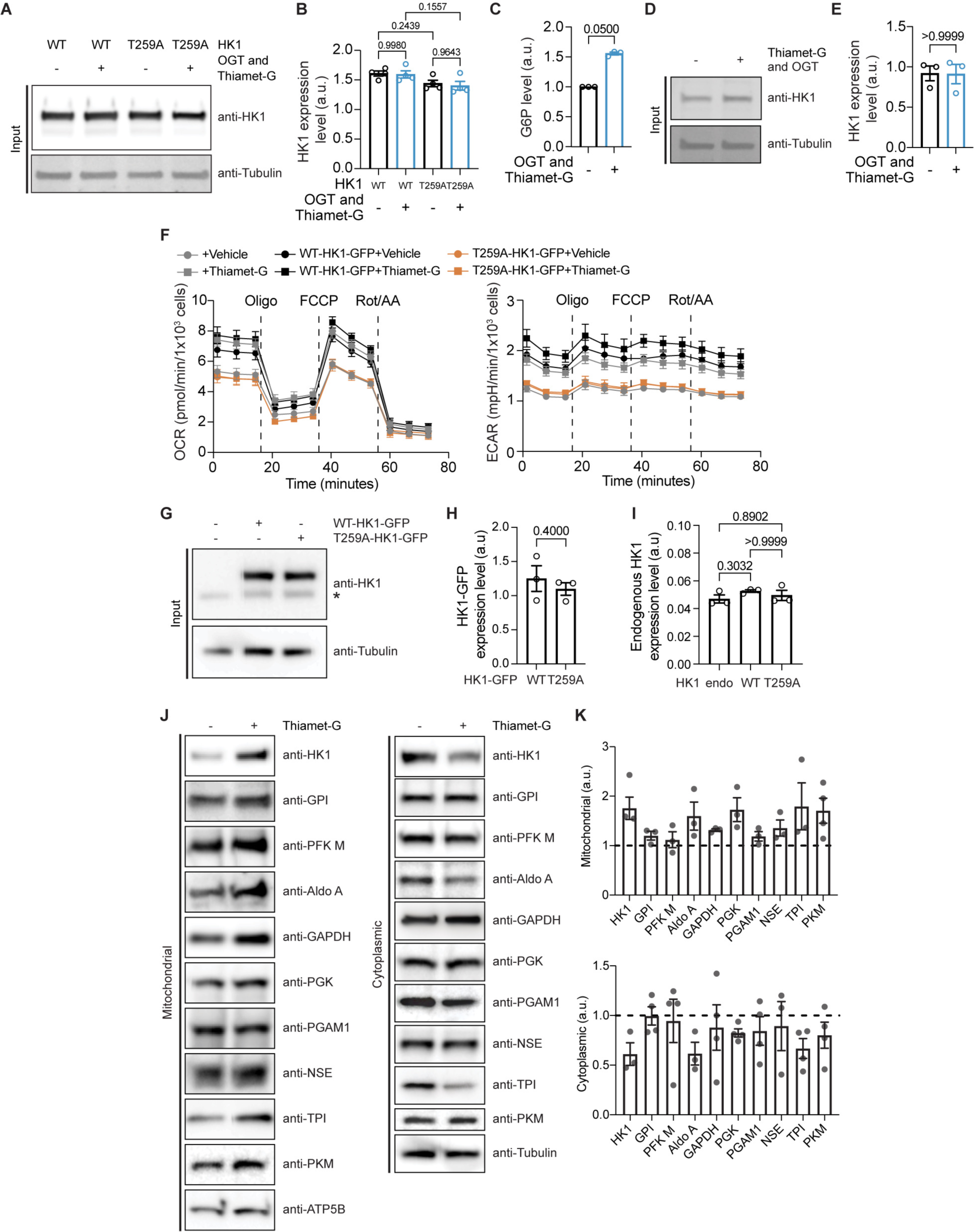
O-GlcNAcylation modifies Hexokinase 1 activity and contributes to the formation of mitochondrial glycosome. (A-B) Quantification of WT and T259A-HK1 expression levels in HEK293T cells, corresponding to the IPs presented in Figure 5C. Whole cell lysates (Input) were probed with anti-HK1 and anti-tubulin (loading control) antibodies. All values are shown as mean ± SEM. n= 4 (Mann-Whitney *U* test). (C-E) Glucose-6-phosphate (G6P) levels were measured in HEK293T cells (maintaining endogenous HK1 levels) following OGT overexpression and treatment with either Thiamet-G or vehicle. G6P levels in untreated cells were set as 1, and fold changes in response to Thiamet-G treatment and OGT overexpression were calculated. (D and E) Endogenous HK1 levels were quantified from whole cell lysates (Input) using anti-HK1 and anti-tubulin (loading control) antibodies. (F) Mitochondrial oxygen consumption rates (left) and extracellular acidification rates (right) were measured in HEK293T cells expressing control vector, eGFP-tagged WT, or T259A HK1, following treatment with either vehicle (DMSO) or overnight Thiamet-G treatment. The subsequent injections of oligomycin (Oligo, 2 μM), FCCP (2 μM), and a combination of rotenone (Rot, 0.5 μM) and antimycin A (AA, 0.5 μM) were used to calculate ATP production rates. All values are presented as the mean ± SEM. (G-I) HK1 levels in HEK293T cells used for metabolic measurements were quantified using western blot analysis of whole cell lysates, probed with antibodies against HK1 and tubulin (serving as a loading control). The asterisk indicates the presence of endogenous HK1. All values are shown as mean ± SEM. n= 3 (Mann-Whitney *U* test). (J and K) Analysis of glycolytic enzymes in mitochondrial and cytoplasmic fractions from rat cortical neurons. Mitochondrial (left) and cytoplasmic fractions (right) from rat cortical neurons, treated overnight with vehicle or Thiamet-G to upregulate O-GlcNAcylation, were analyzed for all glycolytic enzymes using the following antibodies: HK1, Glucose-6-phosphate isomerase (GPI), phosphofructokinase muscle isoform (PFK M), Aldolase A (Aldo A), Glyceraldehyde 3-phosphate dehydrogenase (GAPDH), Phosphoglycerate kinase (PGK), Phosphoglycerate mutase 1 (PGAM1), neuron-specific enolase (NSE), Triosephosphate isomerase (TPI), Pyruvate kinase (PKM), ATP5B (mitochondrial loading control), and Tubulin (cytoplasmic loading control) (J). Quantified enzyme levels under baseline conditions (K) were normalized to 1 (dashed line), and fold changes in response to Thiamet-G treatment were calculated. All values are shown as mean ± SEM. n = 3-4 independent experiments.

**Figure S6.**
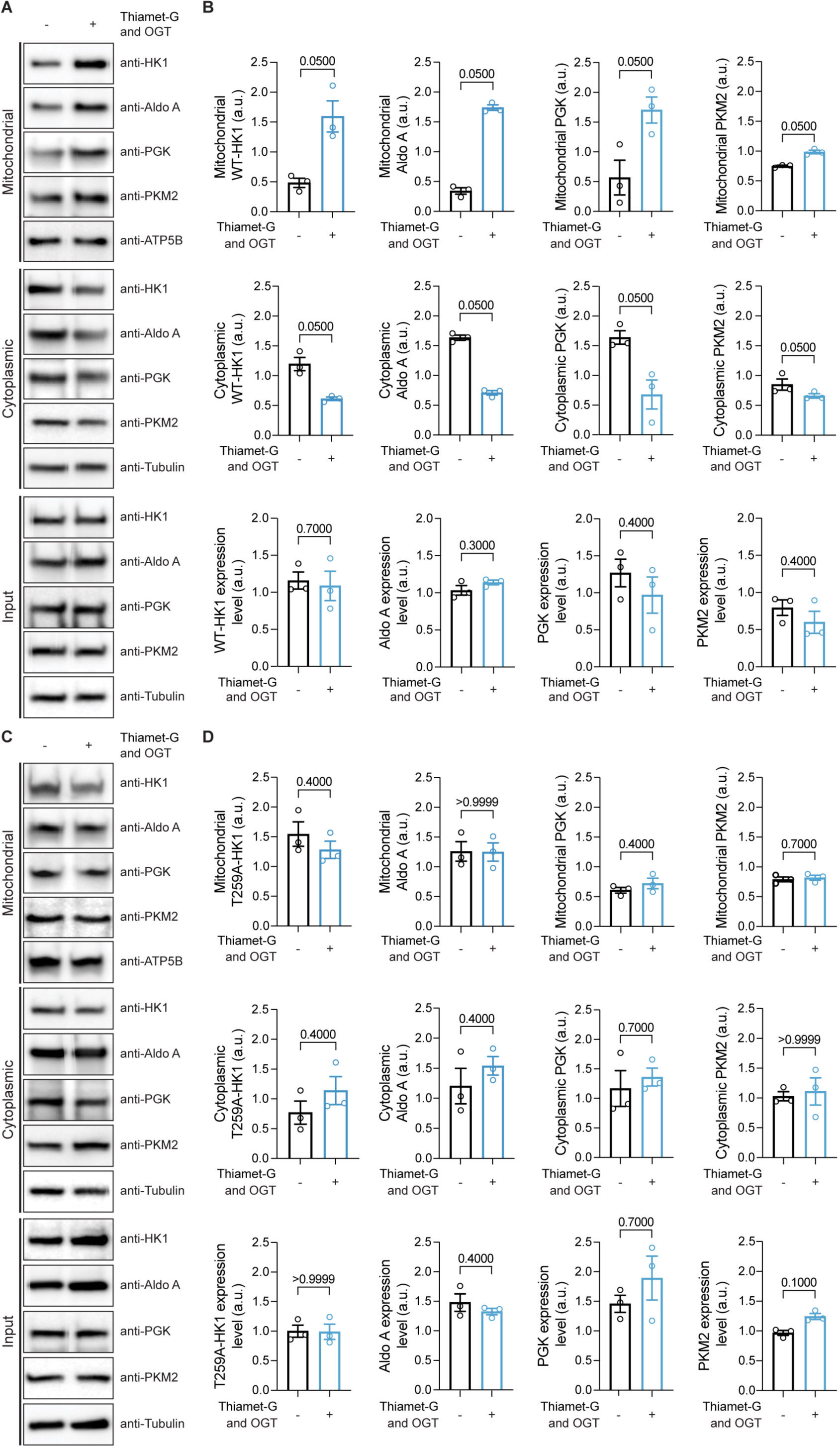
Hexokinase 1 O-GlcNAcylation is required for glycosome formation. (A-D) Analysis of glycolytic enzymes in mitochondrial and cytoplasmic fractions from HEK293T cells expressing WT-HK1-GFP or T259-HK1-GFP. Whole cell lysates, as well as mitochondrial and cytoplasmic fractions from HEK293T cells expressing WT (A and B) and T259A HK1 (C and D) with or without ectopic OGT expression and overnight vehicle (DMSO) or Thiamet-G treatment, were analyzed to quantify the glycolytic enzyme levels. The following antibodies were used: HK1, Aldolase A (Aldo A), Phosphoglycerate kinase (PGK), Pyruvate kinase 2 (PKM2), ATP5B (mitochondrial loading control), and Tubulin (cytoplasmic loading control). Quantified enzyme levels under baseline conditions were normalized to 1, and fold changes in response to OGT overexpression and Thiamet-G treatment were calculated. All values are shown as mean ± SEM. n = 3 independent experiments (Mann-Whitney *U* test).

**Figure S7.**
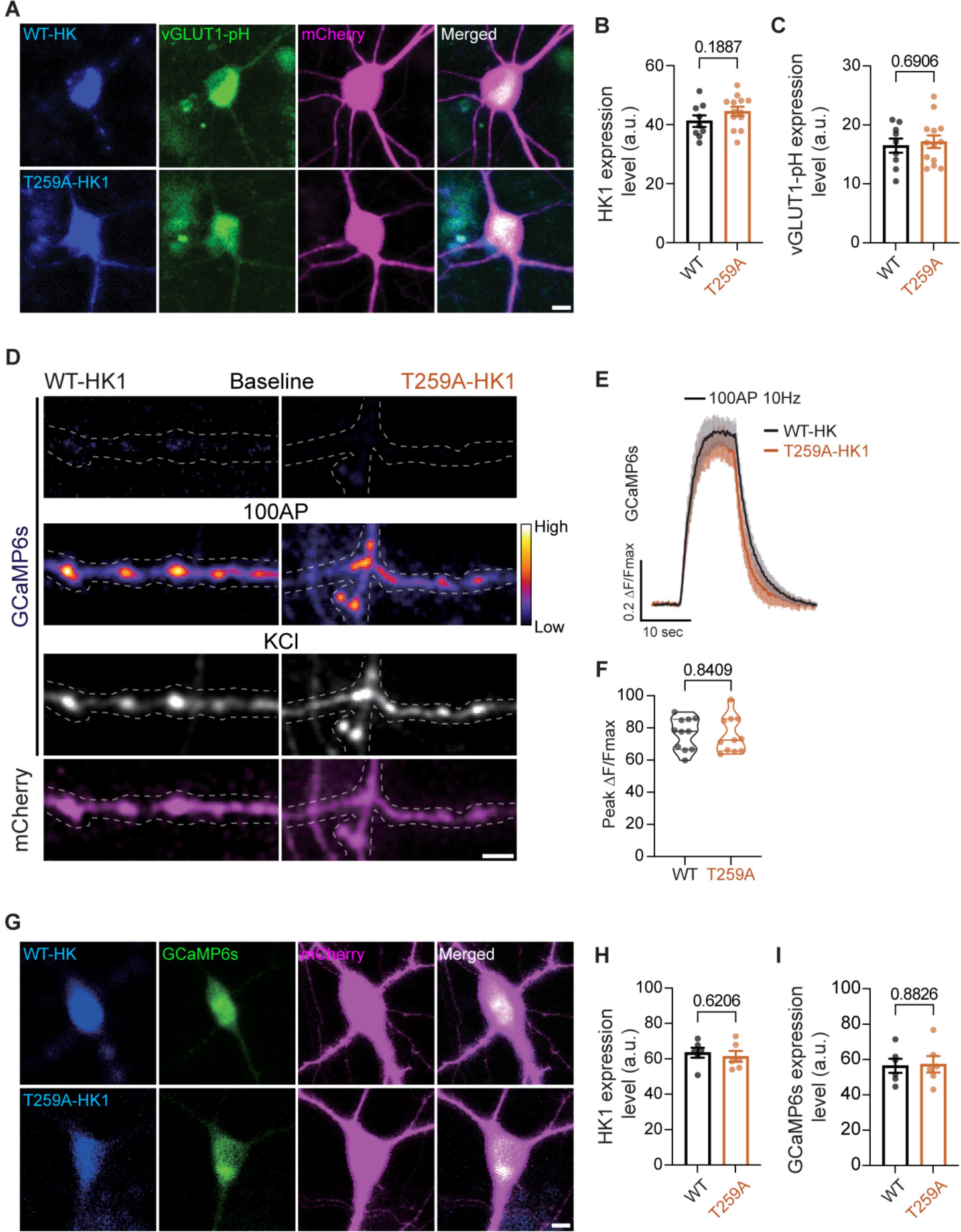
Impact of Hexokinase 1 O-GlcNAcylation on Presynaptic Calcium Dynamics. (A-C) Retrospective quantification of WT-HK1 and T259A-HK1-BFP (blue), vGLUT1-pH (green), and mCherry cell filler (magenta) in rat hippocampal neurons used for imaging experiments depicted in Figure 7. Scale bar represents 10µm. All values are presented as mean ± SEM. n= 12-13 neurons from three independent experiments (unpaired t-test). (D) WT-HK1 or T259A-HK1 together with GCaMP6s were expressed in hippocampal neurons. Neurons were subjected to electrical stimulation with 100 action potentials (APs) at 10Hz. The displayed images illustrate the GCaMP6s signal (pseudo-colored, fire) and the cell filler mCherry (magenta) prior to and following stimulation in neurons transfected with either WT-HK1 or T259A-HK1-BFP with HK1-shRNA to achieve endogenous levels of HK1. The peak of the Ca2+ response elicited by KCl defines the maximum GCaMP6s intensity. The scale bar represents 5µm. (E) The average trace of GCaMP6s during the 100 APs at 10Hz stimulation in neurons transfected with either WT-HK1 or T259A-HK1 was shown. ΔF values were normalized to the maximal ΔF observed during KCl treatment. All values are presented as mean ± SEM. n = 132 ROIs, 12 neurons from three independent experiments. (F) Maximum GCaMP6s ΔF/F values. Data are presented as a violin plot with individual data points and associated p-value. (G-I) Retrospective quantification of WT-HK1 and T259A-HK1-BFP (blue), GCaMP6s, and mCherry cell filler (magenta) in rat hippocampal neurons used for imaging experiments depicted in Figure S7D-F. Scale bar represents 10µm. All values are presented as mean ± SEM. n= 12-13 neurons from three independent experiments (unpaired t-test).

## Notes

### Competing Interest Statement

The authors have declared no competing interest.

### Summary of Updates

This version of the manuscript has been revised to enhance the clarity of the figure legends.

## REFERENCES

1. Welch, G.R. (1987). The living cell as an ecosystem: Hierarchical analog and symmetry. Trends Ecol Evol 2, 305–309. 10.1016/0169-5347(87)90084-X.

2. Patrick, M., Gu, Z., Zhang, G., Wynn, R.M., Kaphle, P., Cao, H., Vu, H., Cai, F., Gao, X., Zhang, Y., et al. (2022). Metabolon formation regulates branched-chain amino acid oxidation and homeostasis. Nat Metab 4, 1775–1791. 10.1038/s42255-022-00689-4.

3. Pedley, A.M., Pareek, V., and Benkovic, S.J. (2022). The Purinosome: A Case Study for a Mammalian Metabolon. Annu Rev Biochem 91, 89–106. 10.1146/annurev-biochem-032620-105728.

4. Alam, M.T., Olin-Sandoval, V., Stincone, A., Keller, M.A., Zelezniak, A., Luisi, B.F., and Ralser, M. (2017). The self-inhibitory nature of metabolic networks and its alleviation through compartmentalization. Nat Commun 8, 16018. 10.1038/ncomms16018.

5. Srere, P.A. (1985). The Metabolon. Trends in Biochem Sci 10, 109–110. 10.1016/0968-0004(85)90266-X.

6. Crane, R.K., and Sols, A. (1953). The association of hexokinase with particulate fractions of brain and other tissue homogenates. J Biol Chem 203, 273–292.

7. Tuttle, J.P., and Wilson, J.E. (1970). Brain hexokinase: a kinetic comparison of soluble and particulate forms. Biochim Biophys Acta 212, 185–188. 10.1016/0005-2744(70)90195-6.

8. Wilson, J.E. (1968). Brain hexokinase. A proposed relation between soluble-particulate distribution and activity in vivo. J Biol Chem 243, 3640–3647.

9. Wilson, J.E. (2003). Isozymes of mammalian hexokinase: structure, subcellular localization and metabolic function. J Exp Biol 206, 2049–2057. 10.1242/jeb.00241.

10. Vaidyanathan, K., Durning, S., and Wells, L. (2014). Functional O-GlcNAc modifications: implications in molecular regulation and pathophysiology. Crit Rev Biochem Mol Biol 49, 140–163. 10.3109/10409238.2014.884535.

11. Sun, L., Shukair, S., Naik, T.J., Moazed, F., and Ardehali, H. (2008). Glucose phosphorylation and mitochondrial binding are required for the protective effects of hexokinases I and II. Mol Cell Biol 28, 1007–1017. 10.1128/MCB.00224-07.

12. Pekkurnaz, G., Trinidad, J.C., Wang, X., Kong, D., and Schwarz, T.L. (2014). Glucose regulates mitochondrial motility via Milton modification by O-GlcNAc transferase. Cell 158, 54–68. 10.1016/j.cell.2014.06.007.

13. Basu, H., Pekkurnaz, G., Falk, J., Wei, W., Chin, M., Steen, J., and Schwarz, T.L. (2021). FHL2 anchors mitochondria to actin and adapts mitochondrial dynamics to glucose supply. J Cell Biol 220. 10.1083/jcb.201912077.

14. Taub, D.G., Awal, M.R., and Gabel, C.V. (2018). O-GlcNAc Signaling Orchestrates the Regenerative Response to Neuronal Injury in Caenorhabditis elegans. Cell Rep 24, 1931–1938 e1933. 10.1016/j.celrep.2018.07.078.

15. Ge, Y., Ramirez, D.H., Yang, B., D’Souza, A.K., Aonbangkhen, C., Wong, S., and Woo, C.M. (2021). Target protein deglycosylation in living cells by a nanobody-fused split O-GlcNAcase. Nat Chem Biol 17, 593–600. 10.1038/s41589-021-00757-y.

16. Ramirez, D.H., Aonbangkhen, C., Wu, H.Y., Naftaly, J.A., Tang, S., O’Meara, T.R., and Woo, C.M. (2020). Engineering a Proximity-Directed O-GlcNAc Transferase for Selective Protein O-GlcNAcylation in Cells. ACS Chem Biol 15, 1059–1066. 10.1021/acschembio.0c00074.

17. Ramirez, D.H., Ge, Y., and Woo, C.M. (2021). O-GlcNAc Engineering on a Target Protein in Cells with Nanobody-OGT and Nanobody-splitOGA. Curr Protoc 1, e117. 10.1002/cpz1.117.

18. Aleshin, A.E., Kirby, C., Liu, X., Bourenkov, G.P., Bartunik, H.D., Fromm, H.J., and Honzatko, R.B. (2000). Crystal structures of mutant monomeric hexokinase I reveal multiple ADP binding sites and conformational changes relevant to allosteric regulation. J Mol Biol 296, 1001–1015. 10.1006/jmbi.1999.3494.

19. Jo, S., and Jiang, W. (2015). A generic implementation of replica exchange with solute tempering (REST2) algorithm in NAMD for complex biophysical simulations. Computer Physics Communications 197, 304–311. 10.1016/j.cpc.2015.08.030.

20. Wang, L., Friesner, R.A., and Berne, B.J. (2011). Replica exchange with solute scaling: a more efficient version of replica exchange with solute tempering (REST2). J Phys Chem B 115, 9431–9438. 10.1021/jp204407d.

21. Rose, I.A., and Warms, J.V. (1967). Mitochondrial hexokinase. Release, rebinding, and location. J Biol Chem 242, 1635–1645.

22. Huh, W.K., Falvo, J.V., Gerke, L.C., Carroll, A.S., Howson, R.W., Weissman, J.S., and O’Shea, E.K. (2003). Global analysis of protein localization in budding yeast. Nature 425, 686–691. 10.1038/nature02026.

23. Jang, S., Xuan, Z., Lagoy, R.C., Jawerth, L.M., Gonzalez, I.J., Singh, M., Prashad, S., Kim, H.S., Patel, A., Albrecht, D.R., et al. (2021). Phosphofructokinase relocalizes into subcellular compartments with liquid-like properties in vivo. Biophys J 120, 1170–1186. 10.1016/j.bpj.2020.08.002.

24. Jin, M., Fuller, G.G., Han, T., Yao, Y., Alessi, A.F., Freeberg, M.A., Roach, N.P., Moresco, J.J., Karnovsky, A., Baba, M., et al. (2017). Glycolytic Enzymes Coalesce in G Bodies under Hypoxic Stress. Cell Rep 20, 895–908. 10.1016/j.celrep.2017.06.082.

25. Jang, S., Nelson, J.C., Bend, E.G., Rodriguez-Laureano, L., Tueros, F.G., Cartagenova, L., Underwood, K., Jorgensen, E.M., and Colon-Ramos, D.A. (2016). Glycolytic Enzymes Localize to Synapses under Energy Stress to Support Synaptic Function. Neuron 90, 278–291. 10.1016/j.neuron.2016.03.011.

26. Miura, N., Shinohara, M., Tatsukami, Y., Sato, Y., Morisaka, H., Kuroda, K., and Ueda, M. (2013). Spatial reorganization of Saccharomyces cerevisiae enolase to alter carbon metabolism under hypoxia. Eukaryot Cell 12, 1106–1119. 10.1128/EC.00093-13.

27. Noree, C., Begovich, K., Samilo, D., Broyer, R., Monfort, E., and Wilhelm, J.E. (2019). A quantitative screen for metabolic enzyme structures reveals patterns of assembly across the yeast metabolic network. Mol Biol Cell 30, 2721–2736. 10.1091/mbc.E19-04-0224.

28. Kohnhorst, C.L., Kyoung, M., Jeon, M., Schmitt, D.L., Kennedy, E.L., Ramirez, J., Bracey, S.M., Luu, B.T., Russell, S.J., and An, S. (2017). Identification of a multienzyme complex for glucose metabolism in living cells. J Biol Chem 292, 9191–9203. 10.1074/jbc.M117.783050.

29. Schmitt, D.L., and An, S. (2017). Spatial Organization of Metabolic Enzyme Complexes in Cells. Biochemistry 56, 3184–3196. 10.1021/acs.biochem.7b00249.

30. Ehsani-Zonouz, A., Golestani, A., and Nemat-Gorgani, M. (2001). Interaction of hexokinase with the outer mitochondrial membrane and a hydrophobic matrix. Mol Cell Biochem 223, 81–87. 10.1023/a:1017952827675.

31. Skaff, D.A., Kim, C.S., Tsai, H.J., Honzatko, R.B., and Fromm, H.J. (2005). Glucose 6-phosphate release of wild-type and mutant human brain hexokinases from mitochondria. J Biol Chem 280, 38403–38409. 10.1074/jbc.M506943200.

32. Rangaraju, V., Calloway, N., and Ryan, T.A. (2014). Activity-driven local ATP synthesis is required for synaptic function. Cell 156, 825–835. 10.1016/j.cell.2013.12.042.

33. Ashrafi, G., Wu, Z., Farrell, R.J., and Ryan, T.A. (2017). GLUT4 Mobilization Supports Energetic Demands of Active Synapses. Neuron 93, 606–615 e603. 10.1016/j.neuron.2016.12.020.

34. Yang, X., and Qian, K. (2017). Protein O-GlcNAcylation: emerging mechanisms and functions. Nat Rev Mol Cell Biol 18, 452–465. 10.1038/nrm.2017.22.

35. De Jesus, A., Keyhani-Nejad, F., Pusec, C.M., Goodman, L., Geier, J.A., Stoolman, J.S., Stanczyk, P.J., Nguyen, T., Xu, K., Suresh, K.V., et al. (2022). Hexokinase 1 cellular localization regulates the metabolic fate of glucose. Mol Cell 82, 1261–1277 e1269. 10.1016/j.molcel.2022.02.028.

36. Singh, J.P., Qian, K., Lee, J.S., Zhou, J., Han, X., Zhang, B., Ong, Q., Ni, W., Jiang, M., Ruan, H.B., et al. (2020). O-GlcNAcase targets pyruvate kinase M2 to regulate tumor growth. Oncogene 39, 560–573. 10.1038/s41388-019-0975-3.

37. Yi, W., Clark, P.M., Mason, D.E., Keenan, M.C., Hill, C., Goddard, W.A., 3rd, Peters, E.C., Driggers, E.M., and Hsieh-Wilson, L.C. (2012). Phosphofructokinase 1 glycosylation regulates cell growth and metabolism. Science 337, 975–980. 10.1126/science.1222278.

38. Nie, H., Ju, H., Fan, J., Shi, X., Cheng, Y., Cang, X., Zheng, Z., Duan, X., and Yi, W. (2020). O-GlcNAcylation of PGK1 coordinates glycolysis and TCA cycle to promote tumor growth. Nat Commun 11, 36. 10.1038/s41467-019-13601-8.

39. Rexach, J.E., Clark, P.M., Mason, D.E., Neve, R.L., Peters, E.C., and Hsieh-Wilson, L.C. (2012). Dynamic O-GlcNAc modification regulates CREB-mediated gene expression and memory formation. Nat Chem Biol 8, 253–261. 10.1038/nchembio.770.

40. Yu, S.B., Sanchez, R.G., Papich, Z.D., Whisenant, T.C., Ghassemian, M., Koberstein, J.N., Stewart, M.L., and Pekkurnaz, G. (2023). Neuronal activity-driven O-GlcNAcylation promotes mitochondrial plasticity. bioRxiv. 10.1101/2023.01.11.523512.

41. Khidekel, N., Ficarro, S.B., Clark, P.M., Bryan, M.C., Swaney, D.L., Rexach, J.E., Sun, Y.E., Coon, J.J., Peters, E.C., and Hsieh-Wilson, L.C. (2007). Probing the dynamics of O-GlcNAc glycosylation in the brain using quantitative proteomics. Nat Chem Biol 3, 339–348. 10.1038/nchembio881.

42. Rexach, J.E., Clark, P.M., and Hsieh-Wilson, L.C. (2008). Chemical approaches to understanding O-GlcNAc glycosylation in the brain. Nat Chem Biol 4, 97–106. 10.1038/nchembio.68.

43. Agrawal, A., Pekkurnaz, G., and Koslover, E.F. (2018). Spatial control of neuronal metabolism through glucose-mediated mitochondrial transport regulation. Elife 7. 10.7554/eLife.40986.

44. Garde, A., Kenny, I.W., Kelley, L.C., Chi, Q., Mutlu, A.S., Wang, M.C., and Sherwood, D.R. (2022). Localized glucose import, glycolytic processing, and mitochondria generate a focused ATP burst to power basement-membrane invasion. Dev Cell 57, 732–749 e737. 10.1016/j.devcel.2022.02.019.

45. Castellana, M., Wilson, M.Z., Xu, Y., Joshi, P., Cristea, I.M., Rabinowitz, J.D., Gitai, Z., and Wingreen, N.S. (2014). Enzyme clustering accelerates processing of intermediates through metabolic channeling. Nat Biotechnol 32, 1011–1018. 10.1038/nbt.3018.

46. Conrado, R.J., Varner, J.D., and DeLisa, M.P. (2008). Engineering the spatial organization of metabolic enzymes: mimicking nature’s synergy. Curr Opin Biotechnol 19, 492–499. 10.1016/j.copbio.2008.07.006.

47. Brandina, I., Graham, J., Lemaitre-Guillier, C., Entelis, N., Krasheninnikov, I., Sweetlove, L., Tarassov, I., and Martin, R.P. (2006). Enolase takes part in a macromolecular complex associated to mitochondria in yeast. Biochim Biophys Acta 1757, 1217–1228. 10.1016/j.bbabio.2006.07.001.

48. Araiza-Olivera, D., Chiquete-Felix, N., Rosas-Lemus, M., Sampedro, J.G., Pena, A., Mujica, A., and Uribe-Carvajal, S. (2013). A glycolytic metabolon in Saccharomyces cerevisiae is stabilized by F-actin. FEBS J 280, 3887–3905. 10.1111/febs.12387.

49. An, S., Parajuli, P., Kennedy, E.L., and Kyoung, M. (2022). Multi-dimensional Fluorescence Live-Cell Imaging for Glucosome Dynamics in Living Human Cells. Methods Mol Biol 2487, 15–26. 10.1007/978-1-0716-2269-8_2.

50. Opperdoes, F.R., and Borst, P. (1977). Localization of nine glycolytic enzymes in a microbody-like organelle in Trypanosoma brucei: the glycosome. FEBS Lett 80, 360–364. 10.1016/0014-5793(77)80476-6.

51. Wojtas, K., Slepecky, N., von Kalm, L., and Sullivan, D. (1997). Flight muscle function in Drosophila requires colocalization of glycolytic enzymes. Mol Biol Cell 8, 1665–1675. 10.1091/mbc.8.9.1665.

52. Sullivan, D.T., MacIntyre, R., Fuda, N., Fiori, J., Barrilla, J., and Ramizel, L. (2003). Analysis of glycolytic enzyme co-localization in Drosophila flight muscle. J Exp Biol 206, 2031–2038. 10.1242/jeb.00367.

53. Diaz-Garcia, C.M., Mongeon, R., Lahmann, C., Koveal, D., Zucker, H., and Yellen, G. (2017). Neuronal Stimulation Triggers Neuronal Glycolysis and Not Lactate Uptake. Cell Metab 26, 361–374 e364. 10.1016/j.cmet.2017.06.021.

54. Diaz-Garcia, C.M., and Yellen, G. (2019). Neurons rely on glucose rather than astrocytic lactate during stimulation. J Neurosci Res 97, 883–889. 10.1002/jnr.24374.

55. Li, H., Guglielmetti, C., Sei, Y.J., Zilberter, M., Le Page, L.M., Shields, L., Yang, J., Nguyen, K., Tiret, B., Gao, X., et al. (2023). Neurons require glucose uptake and glycolysis in vivo. Cell Rep 42, 112335. 10.1016/j.celrep.2023.112335.

56. Baeza-Lehnert, F., Saab, A.S., Gutierrez, R., Larenas, V., Diaz, E., Horn, M., Vargas, M., Hosli, L., Stobart, J., Hirrlinger, J., et al. (2019). Non-Canonical Control of Neuronal Energy Status by the Na(+) Pump. Cell Metab 29, 668–680 e664. 10.1016/j.cmet.2018.11.005.

57. Pulido, C., and Ryan, T.A. (2021). Synaptic vesicle pools are a major hidden resting metabolic burden of nerve terminals. Sci Adv 7, eabi9027. 10.1126/sciadv.abi9027.

58. Rossi, A., Rigotto, G., Valente, G., Giorgio, V., Basso, E., Filadi, R., and Pizzo, P. (2020). Defective Mitochondrial Pyruvate Flux Affects Cell Bioenergetics in Alzheimer’s Disease-Related Models. Cell Rep 30, 2332–2348 e2310. 10.1016/j.celrep.2020.01.060.

59. Han, S., He, Z., Jacob, C., Hu, X., Liang, X., Xiao, W., Wan, L., Xiao, P., D’Ascenzo, N., Ni, J., et al. (2021). Effect of Increased IL-1beta on Expression of HK in Alzheimer’s Disease. Int J Mol Sci 22. 10.3390/ijms22031306.

60. Regenold, W.T., Pratt, M., Nekkalapu, S., Shapiro, P.S., Kristian, T., and Fiskum, G. (2012). Mitochondrial detachment of hexokinase 1 in mood and psychotic disorders: implications for brain energy metabolism and neurotrophic signaling. J Psychiatr Res 46, 95–104. 10.1016/j.jpsychires.2011.09.018.

61. Shan, D., Mount, D., Moore, S., Haroutunian, V., Meador-Woodruff, J.H., and McCullumsmith, R.E. (2014). Abnormal partitioning of hexokinase 1 suggests disruption of a glutamate transport protein complex in schizophrenia. Schizophr Res 154, 1–13. 10.1016/j.schres.2014.01.028.

62. Chen, Z., and Zhong, C. (2013). Decoding Alzheimer’s disease from perturbed cerebral glucose metabolism: implications for diagnostic and therapeutic strategies. Prog Neurobiol 108, 21–43. 10.1016/j.pneurobio.2013.06.004.

63. Huang, C.W., Rust, N.C., Wu, H.F., Yin, A., Zeltner, N., Yin, H., and Hart, G.W. (2023). Low glucose induced Alzheimer’s disease-like biochemical changes in human induced pluripotent stem cell-derived neurons is due to dysregulated O-GlcNAcylation. Alzheimers Dement. 10.1002/alz.13058.

64. Lauretti, E., Li, J.G., Di Meco, A., and Pratico, D. (2017). Glucose deficit triggers tau pathology and synaptic dysfunction in a tauopathy mouse model. Transl Psychiatry 7, e1020. 10.1038/tp.2016.296.

65. Liu, F., Shi, J., Tanimukai, H., Gu, J., Gu, J., Grundke-Iqbal, I., Iqbal, K., and Gong, C.X. (2009). Reduced O-GlcNAcylation links lower brain glucose metabolism and tau pathology in Alzheimer’s disease. Brain 132, 1820–1832. 10.1093/brain/awp099.

66. Zhu, Y., Shan, X., Yuzwa, S.A., and Vocadlo, D.J. (2014). The emerging link between O-GlcNAc and Alzheimer disease. J Biol Chem 289, 34472–34481. 10.1074/jbc.R114.601351.

67. Lagerlof, O. (2018). O-GlcNAc cycling in the developing, adult and geriatric brain. J Bioenerg Biomembr 50, 241–261. 10.1007/s10863-018-9760-1.

68. Wang, A.C., Jensen, E.H., Rexach, J.E., Vinters, H.V., and Hsieh-Wilson, L.C. (2016). Loss of O-GlcNAc glycosylation in forebrain excitatory neurons induces neurodegeneration. Proc Natl Acad Sci U S A 113, 15120–15125. 10.1073/pnas.1606899113.

69. Balana, A.T., and Pratt, M.R. (2021). Mechanistic roles for altered O-GlcNAcylation in neurodegenerative disorders. Biochem J 478, 2733–2758. 10.1042/BCJ20200609.

70. Brenner, S. (1974). The genetics of Caenorhabditis elegans. Genetics 77, 71–94. 10.1093/genetics/77.1.71.

71. Ghanta, K.S., and Mello, C.C. (2020). Melting dsDNA Donor Molecules Greatly Improves Precision Genome Editing in Caenorhabditis elegans. Genetics 216, 643–650. 10.1534/genetics.120.303564.

72. Glomb, O., Swaim, G., LLancao, P.M., Lovejoy, C., Sutradhar, S., Park, J., Wu, Y., Hammarlund, M., Howard, J., Ferguson, S.M., and Yogev, S. (2022). Scaled-expansion of the membrane associated cytoskeleton requires conserved kinesin adaptors. bioRxiv, 2022.2005.2009.491207. 10.1101/2022.05.09.491207.

73. Nie, D., and Sahin, M. (2012). A genetic model to dissect the role of Tsc-mTORC1 in neuronal cultures. Methods Mol Biol 821, 393–405. 10.1007/978-1-61779-430-8_25.

74. Kingston, R.E., Chen, C.A., and Okayama, H. (2001). Calcium phosphate transfection. Curr Protoc Immunol Chapter 10, Unit 10 13. 10.1002/0471142735.im1013s31.

75. Ashrafi, G., de Juan-Sanz, J., Farrell, R.J., and Ryan, T.A. (2020). Molecular Tuning of the Axonal Mitochondrial Ca(2+) Uniporter Ensures Metabolic Flexibility of Neurotransmission. Neuron 105, 678-687 e675. 10.1016/j.neuron.2019.11.020.

76. Djakovic, S.N., Schwarz, L.A., Barylko, B., DeMartino, G.N., and Patrick, G.N. (2009). Regulation of the proteasome by neuronal activity and calcium/calmodulin-dependent protein kinase II. J Biol Chem 284, 26655–26665. 10.1074/jbc.M109.021956.

77. Allison, D.F., Wamsley, J.J., Kumar, M., Li, D., Gray, L.G., Hart, G.W., Jones, D.R., and Mayo, M.W. (2012). Modification of RelA by O-linked N-acetylglucosamine links glucose metabolism to NF-kappaB acetylation and transcription. Proc Natl Acad Sci U S A 109, 16888–16893. 10.1073/pnas.1208468109.

78. Kreppel, L.K., Blomberg, M.A., and Hart, G.W. (1997). Dynamic glycosylation of nuclear and cytosolic proteins. Cloning and characterization of a unique O-GlcNAc transferase with multiple tetratricopeptide repeats. J Biol Chem 272, 9308–9315. 10.1074/jbc.272.14.9308.

79. Chung, J.Y., Steen, J.A., and Schwarz, T.L. (2016). Phosphorylation-Induced Motor Shedding Is Required at Mitosis for Proper Distribution and Passive Inheritance of Mitochondria. Cell Rep 16, 2142–2155. 10.1016/j.celrep.2016.07.055.

80. Chen, T.W., Wardill, T.J., Sun, Y., Pulver, S.R., Renninger, S.L., Baohan, A., Schreiter, E.R., Kerr, R.A., Orger, M.B., Jayaraman, V., et al. (2013). Ultrasensitive fluorescent proteins for imaging neuronal activity. Nature 499, 295–300. 10.1038/nature12354.

81. Schindelin, J., Arganda-Carreras, I., Frise, E., Kaynig, V., Longair, M., Pietzsch, T., Preibisch, S., Rueden, C., Saalfeld, S., Schmid, B., et al. (2012). Fiji: an open-source platform for biological-image analysis. Nat Methods 9, 676–682. 10.1038/nmeth.2019.

82. Manders, E.M.M., Verbeek, F.J., and Aten, J.A. (1993). Measurement of co-localization of objects in dual-colour confocal images. J Microsc 169, 375–382. 10.1111/j.1365-2818.1993.tb03313.x.

83. Whelan, S.A., Lane, M.D., and Hart, G.W. (2008). Regulation of the O-linked beta-N-acetylglucosamine transferase by insulin signaling. J Biol Chem 283, 21411–21417. 10.1074/jbc.M800677200.

84. Shevchenko, A., Wilm, M., Vorm, O., and Mann, M. (1996). Mass spectrometric sequencing of proteins silver-stained polyacrylamide gels. Anal Chem 68, 850–858. 10.1021/ac950914h.

85. Roy, A., Kucukural, A., and Zhang, Y. (2010). I-TASSER: a unified platform for automated protein structure and function prediction. Nat Protoc 5, 725–738. 10.1038/nprot.2010.5.

86. Jo, S., Kim, T., Iyer, V.G., and Im, W. (2008). CHARMM-GUI: a web-based graphical user interface for CHARMM. J Comput Chem 29, 1859–1865. 10.1002/jcc.20945.

87. Terakawa, T., Kameda, T., and Takada, S. (2011). On easy implementation of a variant of the replica exchange with solute tempering in GROMACS. J Comput Chem 32, 1228–1234. 10.1002/jcc.21703.

88. Jurrus, E., Engel, D., Star, K., Monson, K., Brandi, J., Felberg, L.E., Brookes, D.H., Wilson, L., Chen, J., Liles, K., et al. (2018). Improvements to the APBS biomolecular solvation software suite. Protein Sci 27, 112–128. 10.1002/pro.3280.

89. Huang, J., and MacKerell, A.D., Jr. (2013). CHARMM36 all-atom additive protein force field: validation based on comparison to NMR data. J Comput Chem 34, 2135–2145. 10.1002/jcc.23354.

90. Heo, S., Diering, G.H., Na, C.H., Nirujogi, R.S., Bachman, J.L., Pandey, A., and Huganir, R.L. (2018). Identification of long-lived synaptic proteins by proteomic analysis of synaptosome protein turnover. Proc Natl Acad Sci U S A 115, E3827–E3836. 10.1073/pnas.1720956115.

91. Armbruster, M., and Ryan, T.A. (2011). Synaptic vesicle retrieval time is a cell-wide rather than individual-synapse property. Nat Neurosci 14, 824–826. 10.1038/nn.2828.

